# Type 2 diabetes promotes cell centrosome amplification and the role of AKT-ROS-dependent signalling of ROCK1 and 14-3-3σ

**DOI:** 10.1101/181214

**Authors:** Pu Wang, Yu Cheng Lu, Jie Wang, Lan Wang, Hanry Yu, Yuan Fei Li, Alice Kong, Juliana Chan, Shao Chin Lee

**Author notes:** Correspondence: Dr. Shao Chin Lee, School of Life Sciences, Shanxi University, Taiyuan, Shanxi, PR China.

## Abstract

Type2 diabetes is associated with oxidative stress which can cause cell centrosome amplification. The study investigated centrosome amplification in type 2 diabetes and the underlying mechanisms. We found that centrosome amplification was increased in the peripheral blood mononuclear cells (PBMC) from the type 2 diabetic patients, which correlated with the levels of fasting blood glucose and HbA1c. High glucose, insulin and palmitic acid, alone or in combinations, induced ROS production and centrosome amplification. Together, they increased AKT activation as well as the expression, binding and centrosome translation of ROCK1 and 14-3-3σ. Results from further analyses showed that AKT-ROS-dependent upregulations of expression, binding and centrosome translocation of ROCK1 and 14-3-3σ was the molecular pathway underlying the centrosome amplification induced by high glucose, insulin and palmitic acid. Moreover, the increases in AKT activation and ROS production as well as expression, binding and centrosome distribution of ROCK1 and 14-3-3σ were confirmed in the PBMC from the patients with type 2 diabetes. In conclusion, our results show that type 2 diabetes promotes cell centrosome amplification, and suggest that the diabetic pathophysiological factors-activated AKT-ROS-dependent signalling of ROCK1 and 14-3-3σ is the underlying molecular mechanism.

## INTRODUCTION

Type 2 diabetes is a common non-communicable disease. At certain stage of the disease development, patients present pathophysiological features such as hyperglycemia, hyperinsulinemia and increased plasma level of free fatty acids. At latter stage, patients may develop insulin deficiency, but continue to have hyperglycemia and increased plasma level of free fatty acids. These pathophysiological factors are able to trigger the production of reactive oxygen species (ROS) (Inoguchi et al., 2000; May and de Haen, 1979) and DNA damage (Lee and Chan, 2015) *in vitro*, which may explain why oxidative stress and DNA damage are increased *in vivo* in patients with type 2 diabetes (Lee and Chan, 2015; Orie et al., 1999).

Cell centrosome amplification refers to a cell with more than two centrosomes. There is evidence that oxidative stress and DNA damage can cause centrosome amplification (Chae et al., 2005; Dodson et al., 2004). At the molecular level, many signal mediators for cell centrosome amplification have been identified, which include those inside or outside the centrosomes. Overexpression of PLK4 (Basto et al., 2008) or aurora A (Castellanos et al., 2008), both are centrosome proteins, can cause centrosome amplification. Wang and co-workers show that Chk2 is located in centrosome, which controls centrosome amplification induced by sub-toxic concentrations of hydroxyurea (Wang et al., 2015). Overexpression of cyclin E (Bagheri-Yarmand et al., 2010) which can be in centrosome was also to promote centrosome amplification. Ras (Zen et al., 2010) which is a protein outside the centrosomes has been shown to cause centrosome amplification. Interestingly, BRCA1 promotes centrosome amplification when is translocated to centrosome (Zou et al., 2014), which shows a dynamic communication between centrosome and its surrounding environment for centrosome amplification. At the cellular and organismal levels, centrosome amplification regulates cell division, movement and intracellular transport (Anderhub et al., 2012), and increases cancer invasion potential (Godinho et al., 2014). While robust centrosome amplification results in multipolar division follow by mitosis catastrophe (Pannu et al., 2012), there is evidence that moderate centrosome amplification can induce genome instability (Ganem et al., 2009) and tumorigenesis (Levine et al., 2017).

Since type 2 diabetes is associated with oxidative stress (Inoguchi et al., 2000; May and de Haen, 1979; Lee and Chan, 2015; Orie et al., 1999) which can cause centrosome amplification (Chae et al., 2005; Dodson et al., 2004), in the present study, we investigate cell centrosome amplification in type 2 diabetes and the underlying mechanisms.

## RESULTS

### Cell centrosome amplification is increased in patients with type 2 diabetes, which is in correlation with fasting blood glucose and HbA1c

To investigate whether centrosome amplification is associated with type 2 diabetes, first of all, we compared the extent of centrosome amplification in PBMCs from 32 patients with type 2 diabetes and 12 healthy subjects. The clinical characteristics of the volunteers were summarized in Table 1. Compared with the healthy subjects, the diabetic patients had higher body mass index, fasting blood glucose, HbA1c and triglyceride. Moreover, we compared the level of centrosome amplification between the healthy subject and diabetic patients. The images of centrosome and centrosome amplification were shown in Fig. 1A. The patients had a greater degree of cell centrosome amplification than the healthy subjects (8.7 % v.s.3.4%; p<0.01) (Fig. 1B); there was a 2.6-fold increase in centrosome amplification in PBMCs from the patients. When the diabetic patients were divided into two groups according to their fasting blood glucose level less or greater than 10mM, there was a step-wise increase in centrosome amplification (p<0.01; Fig. 1C). Indeed, in all the volunteers, correlation analysis showed that the extent of cell centrosome amplification was correlated with the fasting blood glucose level (R^2^ = 0.7; p<0.01; Fig. 1D) and HbA1c (R^2^=0.7; p<0.01; Fig. 1E). Similarly, in the patients alone, the level of centrosome amplification was also correlated with fasting blood glucose (R^2^ = 0.5; p <0.01; Fig. 1F) and HbA1c (R^2^=0.4; p<0.01; Fig. 1G).

**Table 1.** Clinical and biochemical characteristics of the volunteers

|  | Healthy<br>subject | Diabetic<br>patient |
| --- | --- | --- |
| n | 12 | 32 |
| Age (years) | 57±2 | 55±2 |
| Sex ratio (male:female) | 1:1 | 1:1 |
| Body mass index (kg/m<sup>2</sup>) | 19.8±2.7 | 26.7±5* |
| Systolic blood pressure (mmHg) | 133.3±5.7 | 131.7±17.2 |
| Diastolic blood pressure (mmHg) | 76.6±11.5 | 83.3±10.3 |
| Fasting blood glucose (mmol/l) | 5.3±0.3 | 10.1±3.7* |
| HbA1c (%) | 5.6± | 8.5±2.2* |
| Total cholesterol (mmol/l) | 2.8±1.4 | 4.9±1.4 |
| Triglyceride (mmol/l) | 0.66(0.2-1.1) | 2.3(1.2-3.5)* |

Data are means ± SD, or medians (range). \*p<0.05

**Fig. 1.**
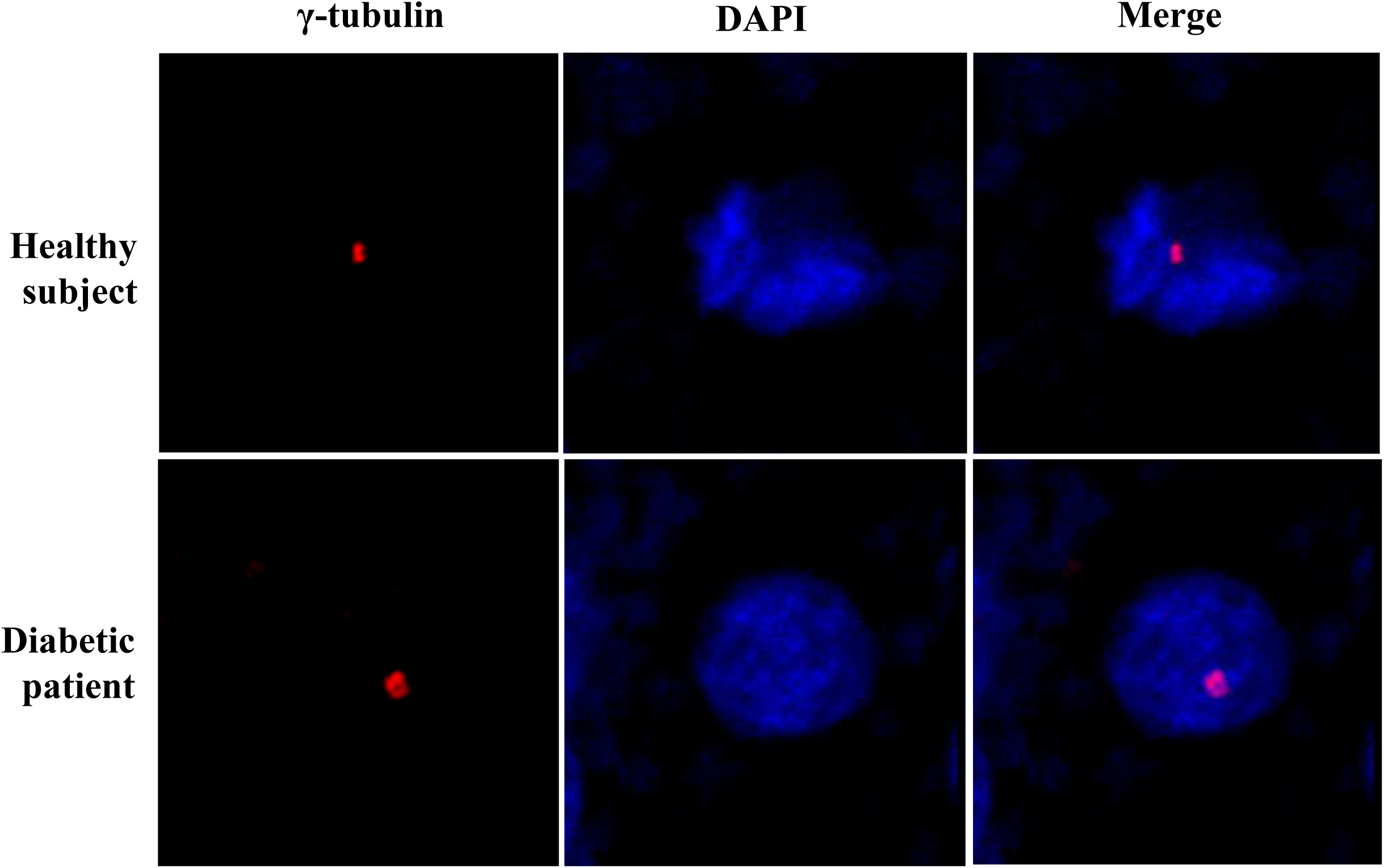
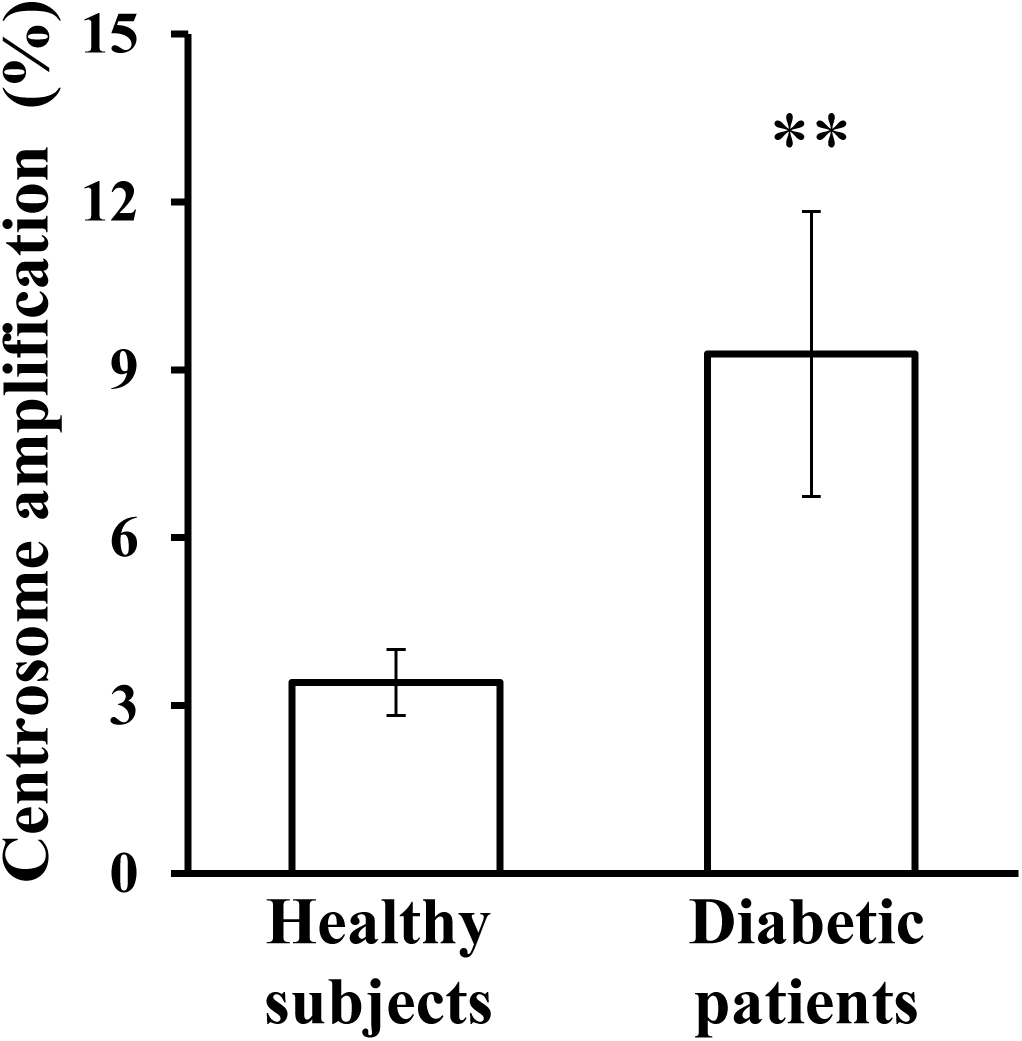
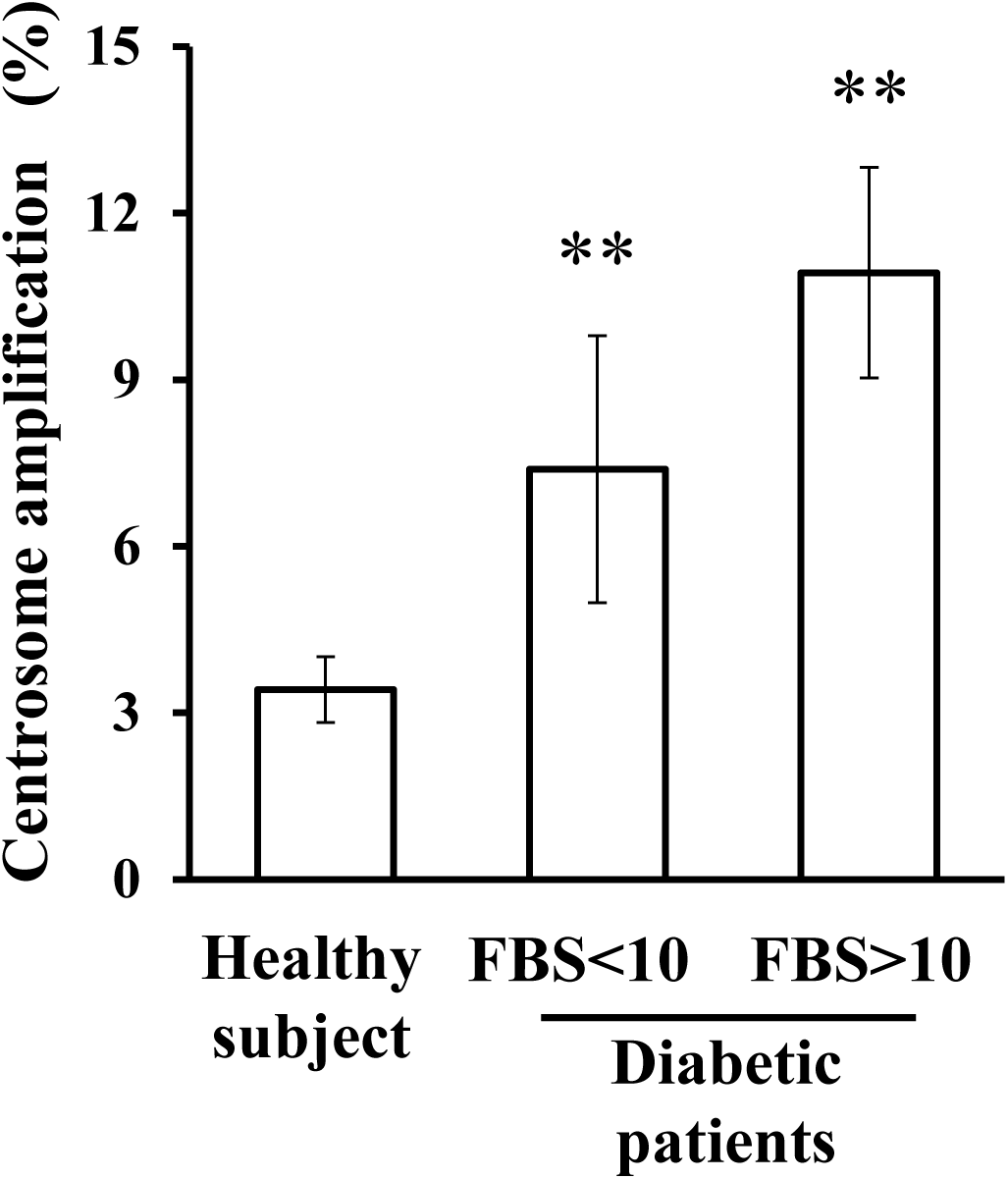
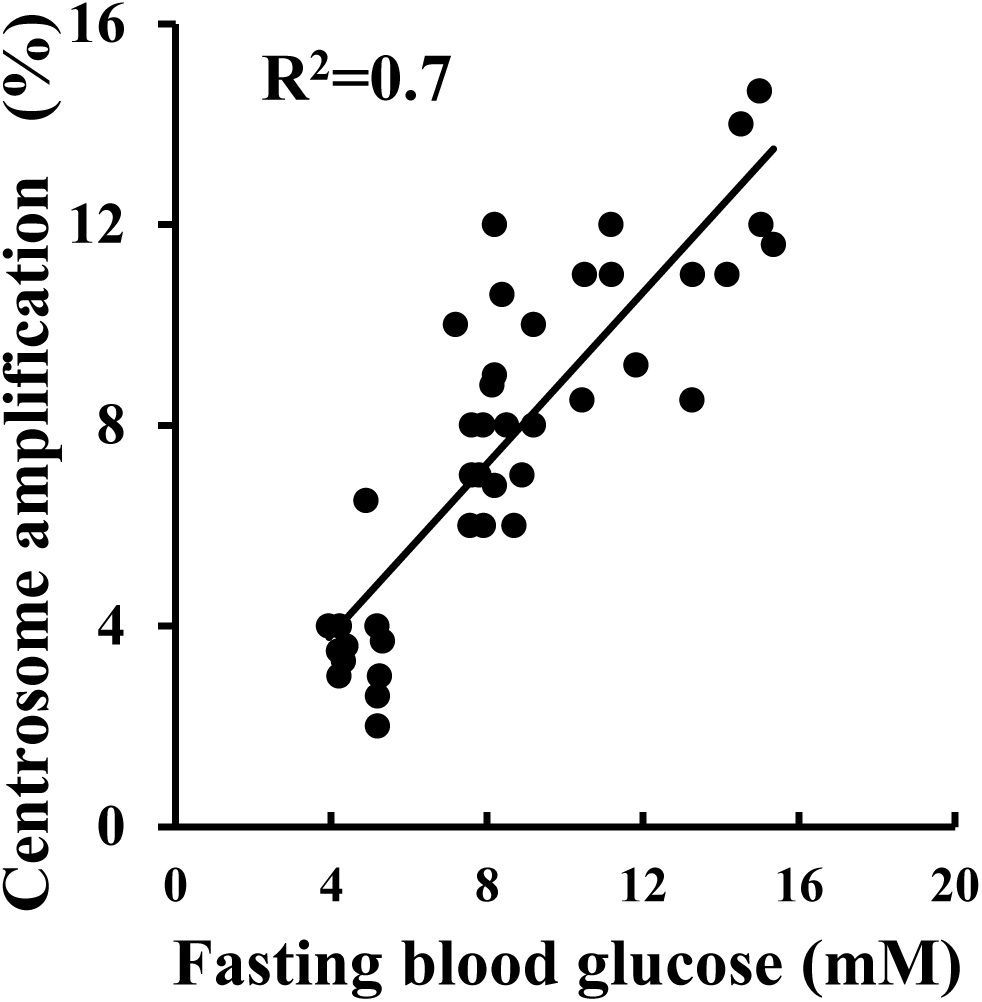
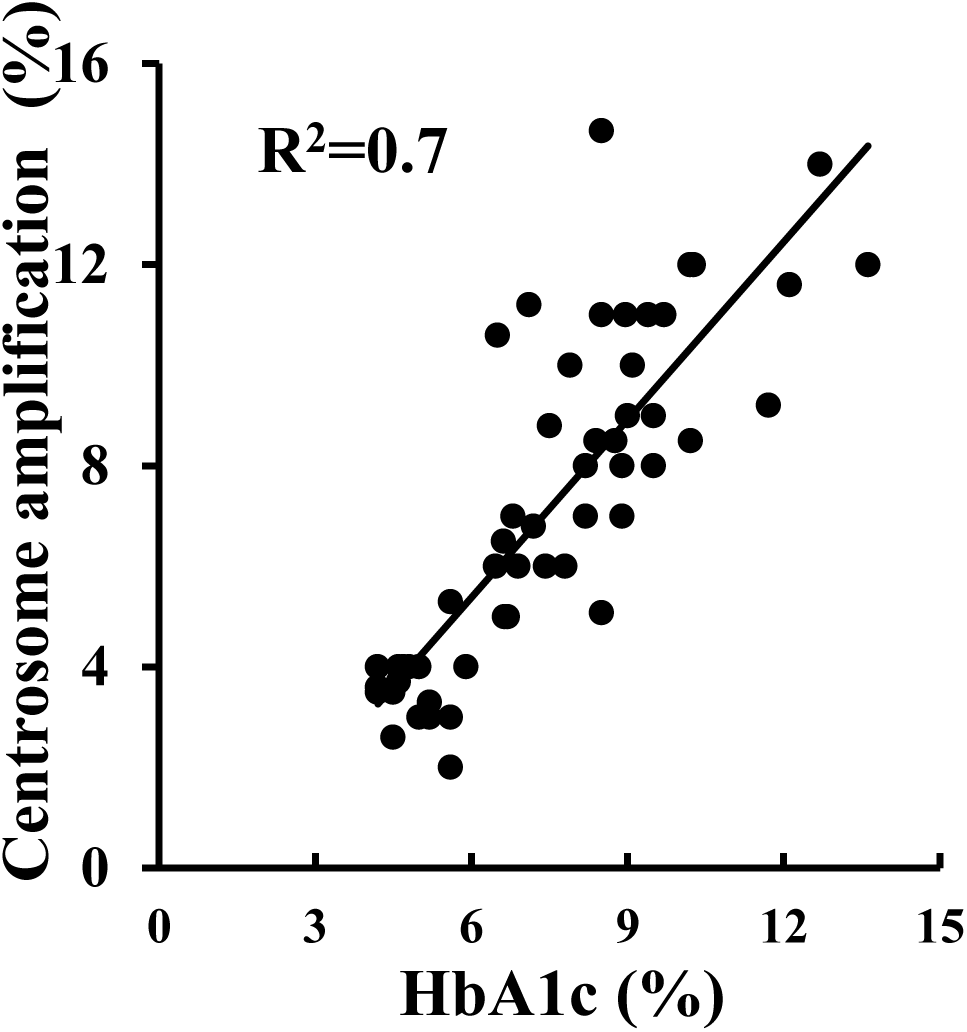
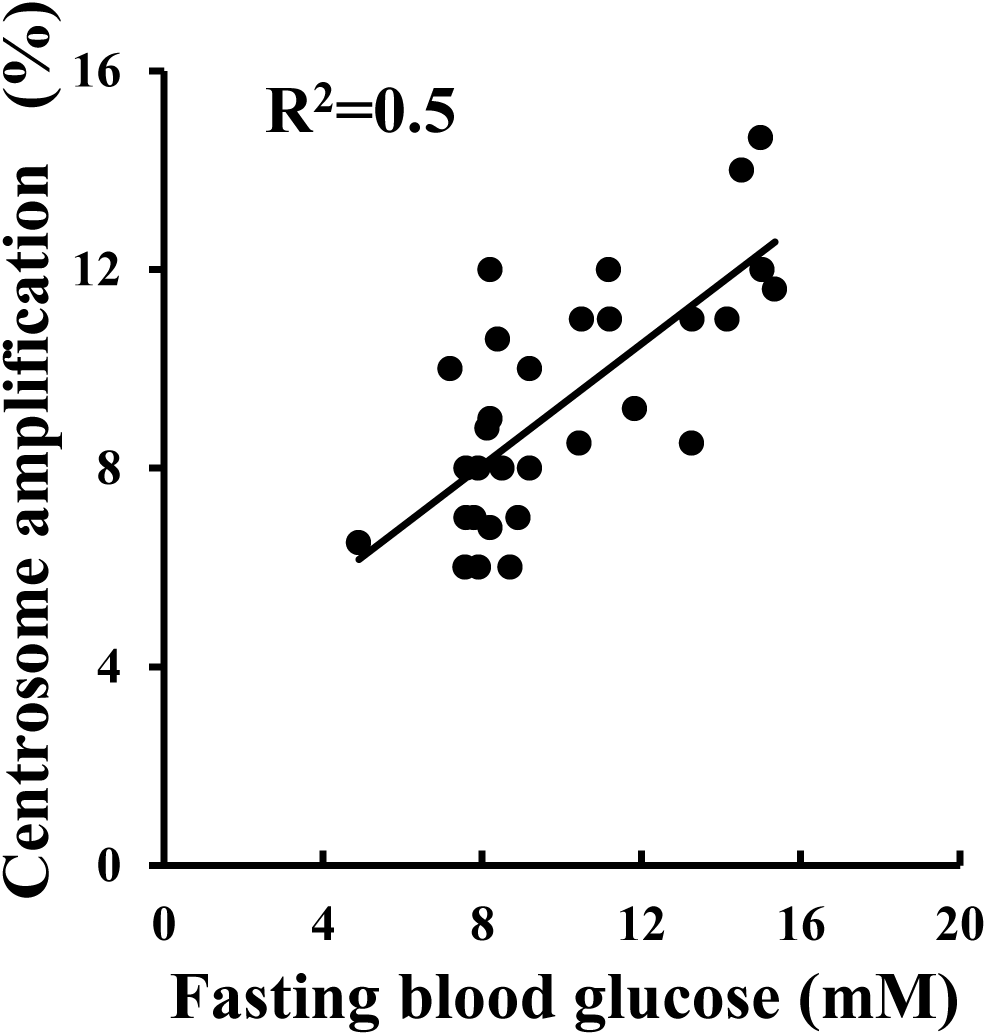
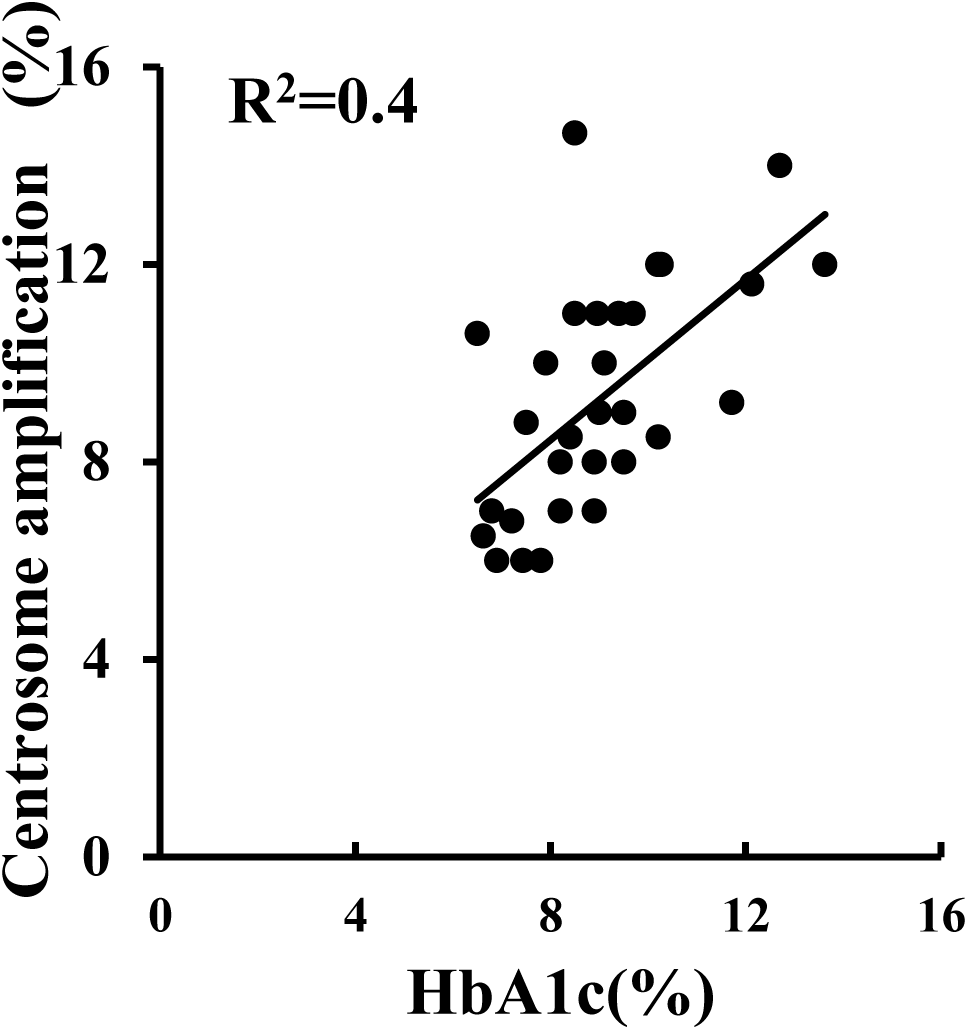
Cell centrosome amplification is increased in patients with type 2 diabetes, which is in correlation with fasting blood glucose and HbA1c. (**A**): image of centrosome amplification; (**B**): cell centrosome amplification is increased in patients with type 2 diabetes; (**C**): evidence that centrosome amplification correlates with the fasting blood glucose; (**D**) and (**E**): correlation analysis shows that the centrosome amplification correlates with the fasting blood glucose and HbA1c in all the volunteers; (**F**) and (**G**): in the diabetic patients alone, centrosome amplification also correlates with fasting blood glucose and HbA1c. Student t-test was used to compare the means between two groups. One way ANOVA analysis was used to compare multiple groups. Linear regression was performed for correlations. *: p < 0.05; **: p < 0.01, compared with that in the control group.

### High glucose, insulin and palmitic acid, alone or in combinations, induce centrosome amplification via ROS production

We then investigated whether the pathophysiological factors in type 2 diabetes could induce centrosome amplification via specific molecular pathway using colon cancer cells as an experimental model. Some results were verified in normal human breast epithelial cells wherever it was considered to be meaningful, which was to verify findings from experiments using cancer cells in non-cancerous cell model. Palmitic acid, the most common saturated free fatty acid, was used to represent free fatty acids. We tested ROS pathway to start with. Fluorescent spectrophotometry was used to quantify the level of ROS when experiments involved more than three samples. Otherwise, flow cytometry analysis was employed. The results showed that high glucose, insulin and palmitic acid, alone or in combinations, significantly induced ROS production (Fig. 2A) and centrosome amplification (Figs. 2C) in the cancer cells. High glucose, insulin and palmitic acid individually induced centrosome amplification in concentration-dependent manners (figures not shown). They, together, also triggered ROS production (Fig. 2B) and centrosome amplification (Fig. 2D) in the normal human breast epithelial cells. Antioxidant N-acetylcysteine (NAC) was able to inhibit the ROS production and the centrosome amplification (Figs. 2A-2D). The potential of high glucose, insulin and palmitic acid in inducing centrosome amplification was three factors > two factors > single factor (all at p<0.05; Fig. 2C).

**Fig. 2.**
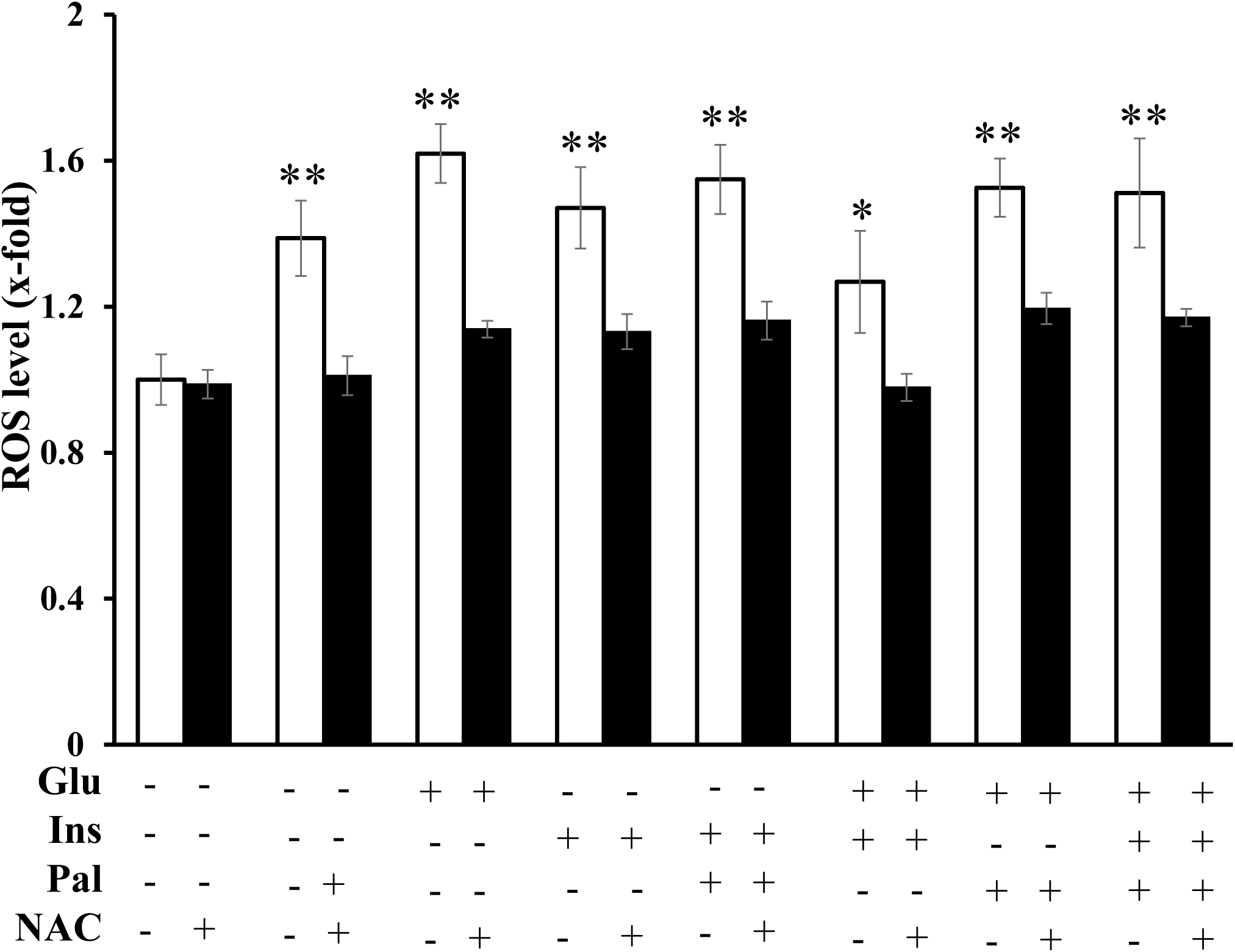
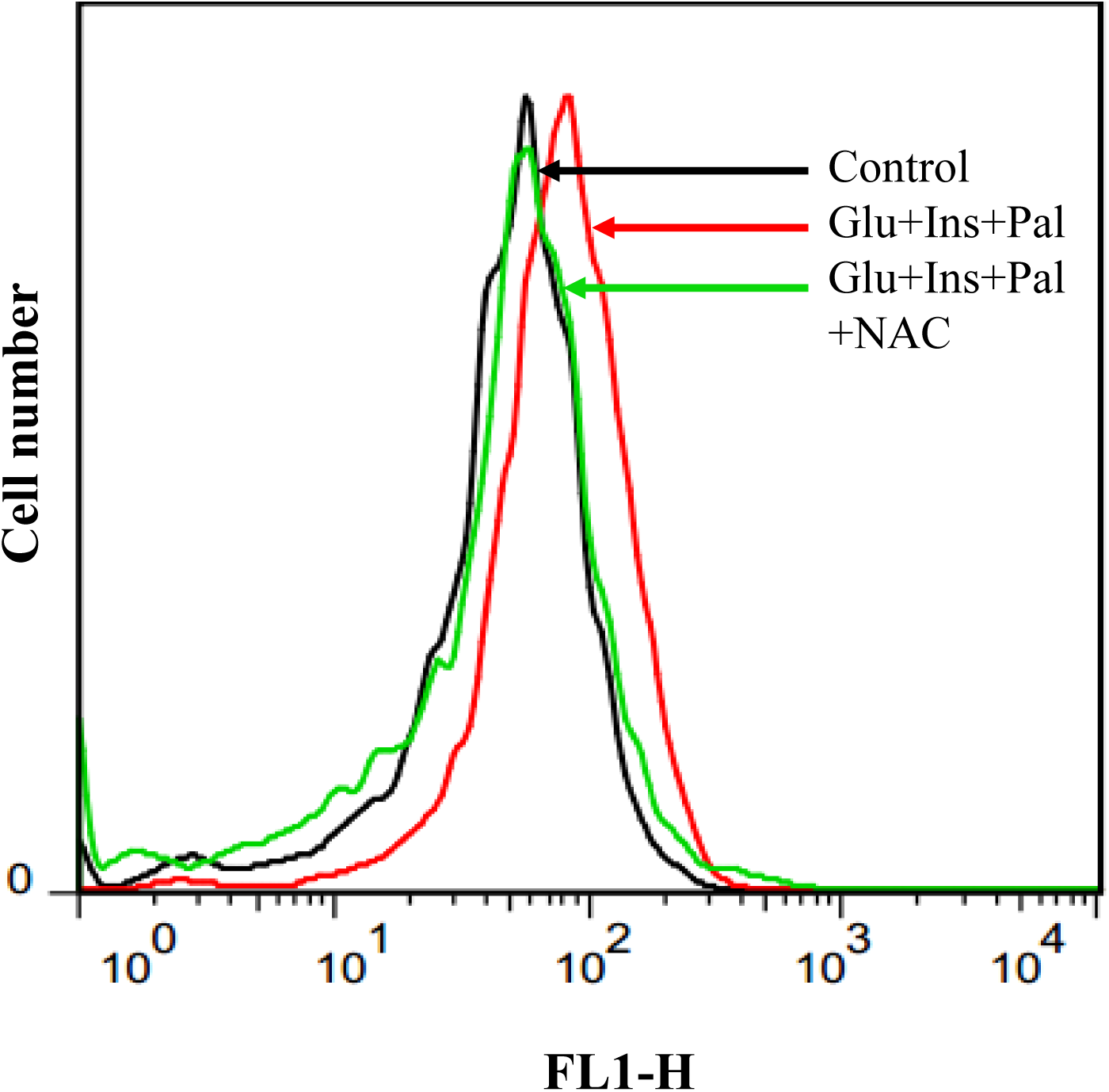
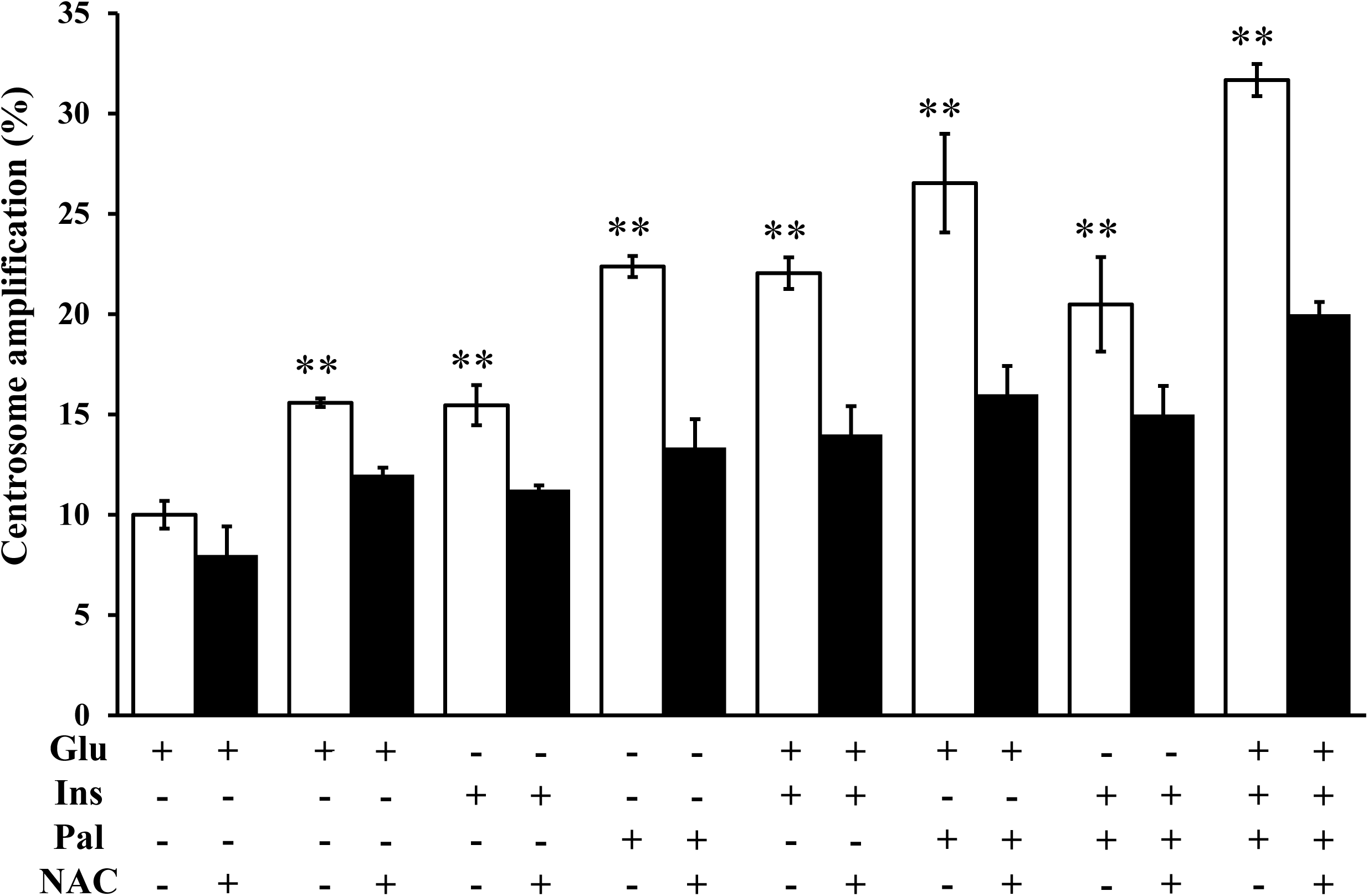
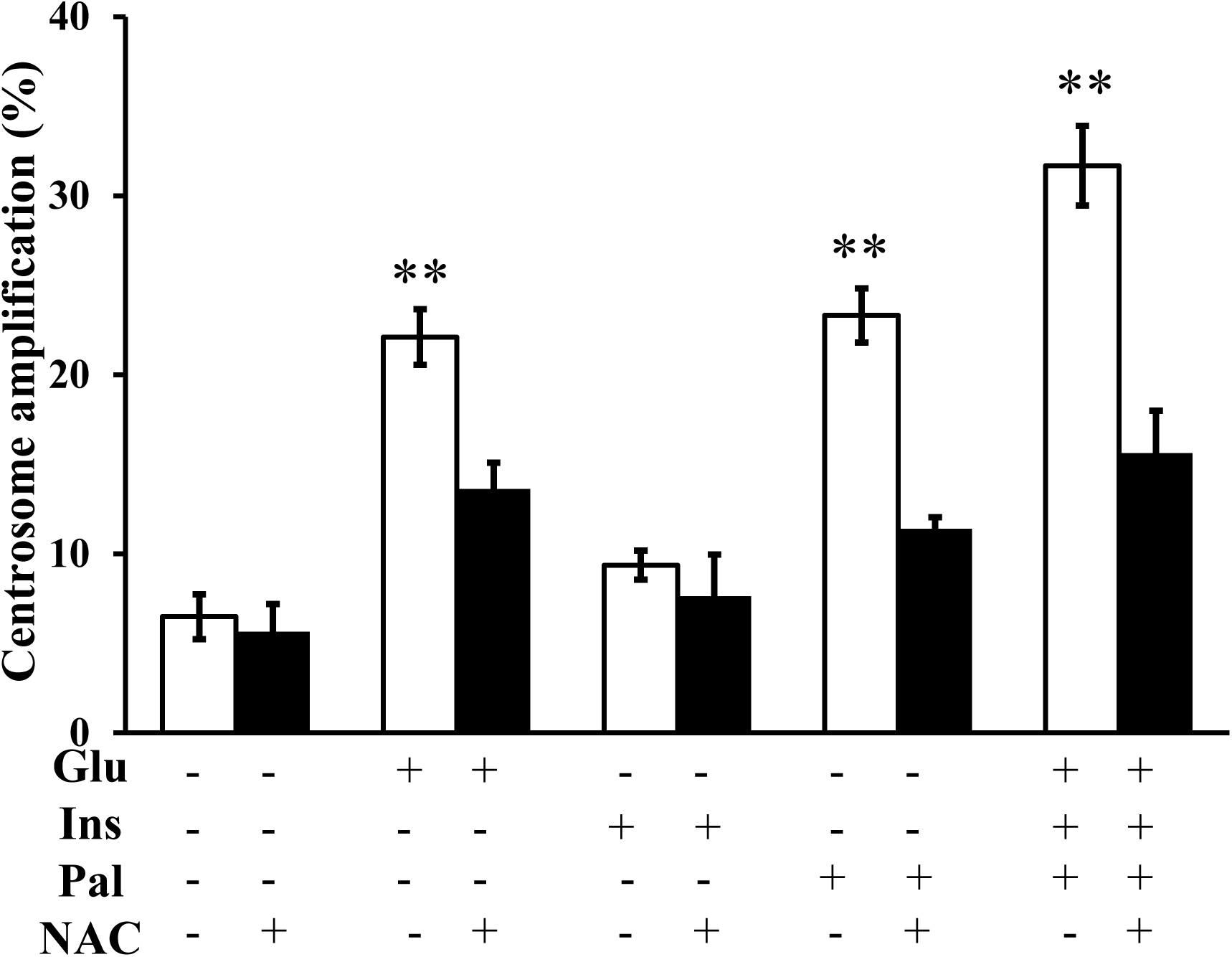
High glucose, insulin and palmitic acid, alone or in combinations, induce centrosome amplification via ROS production. (**A**): high glucose, insulin and palmitic acid, alone or in combinations, triggers ROS production in the cancer cells; (**B**): the pathophysiological factors, together, trigger ROS production in normal human breast epithelial cell; (**C**): high glucose, insulin and palmitic acid, alone or in combinations, can promote centrosome amplification in the cancer cells; (**D**): high glucose, insulin and palmitic acid can trigger centrosome amplification in the normal human breast epithelial cells. (**A**)-(**D**): NAC inhibits the ROS production and the centrosome amplification. Glu: glucose, 15 mM; Ins: insulin, 5 nM; Pal: palmitic acid, 150 µM; NAC: 3 mM. One way ANOVA analysis was used to compare multiple groups. *: p < 0.05; **: p < 0.01.

### Functional transcriptomic analysis identifies AKT, ROCK1 and 14-3-3σ as signal mediators for the centrosome amplification

To further identify the molecular signals for the centrosome amplification, we performed a functional transcriptomic analysis between the control samples and those treated with high glucose, insulin and palmitic acid. The differentially expressed genes were summarized in the supplementary data sheet (Table S1). In total, 729 genes were differentially expressed, with 508 upregulated and 221 downregulated. Results of bioinformatic annotation were included in supplementary figures S1-S5. Functional enrichment using GO analysis assigned the genes to 37 terms in three categories of molecular component, molecular function and biological process (Fig. S1). The top terms were molecular function, cellular component, biological process and cell, which had 9%, 9%, 8.4% and 8.3% of the genes, respectively. KEGG annotation identified 37 pathways which were related to different functional groups (Fig. S2). The top pathways were signal transduction, infectious diseases, cancer, immune system and endocrine system, which had 15.6%, 11.6%, 11.1%, 7.4% and 7.0% of the genes, respectively. Notably, AKT (Fig.S3), ROCK (Fig. S4) and 14-3-3σ (Fig.S5) pathways were activated, which were chosen for further functional analysis. Western blot analyses confirmed that high glucose, insulin and palmitic acid activated AKT, which was inhibited by AKT inhibitor Ly294002 or siRNA (Fig. 3A). Similarly, ROCK1 (Fig. 3B) and 14-3-3σ (Fig. 3C) protein levels were upregulated, which were inhibited their specific siRNA (Figs. 3B and 3C). We then performed functional analyses to examine whether AKT, ROCK1 and 14-3-3σ mediated the centrosome amplification. Indeed, inhibition of AKT using chemical inhibitor or siRNA could inhibit the centrosome amplification (Fig. 3D). Similarly, siRNA for ROCK1 (Fig. 3E) and siRNA for 14-3-3σ (Fig. 3F) were also able to inhibit the centrosome amplification. Moreover, AKR chemical inhibitor, AKT siRNA, ROCK1 siRNA and 14-3-3σ siRNA individually were all able to inhibit the centrosome in the normal human breast epithelial cells (Figs. 3G and 3H).

**Fig. 3.**
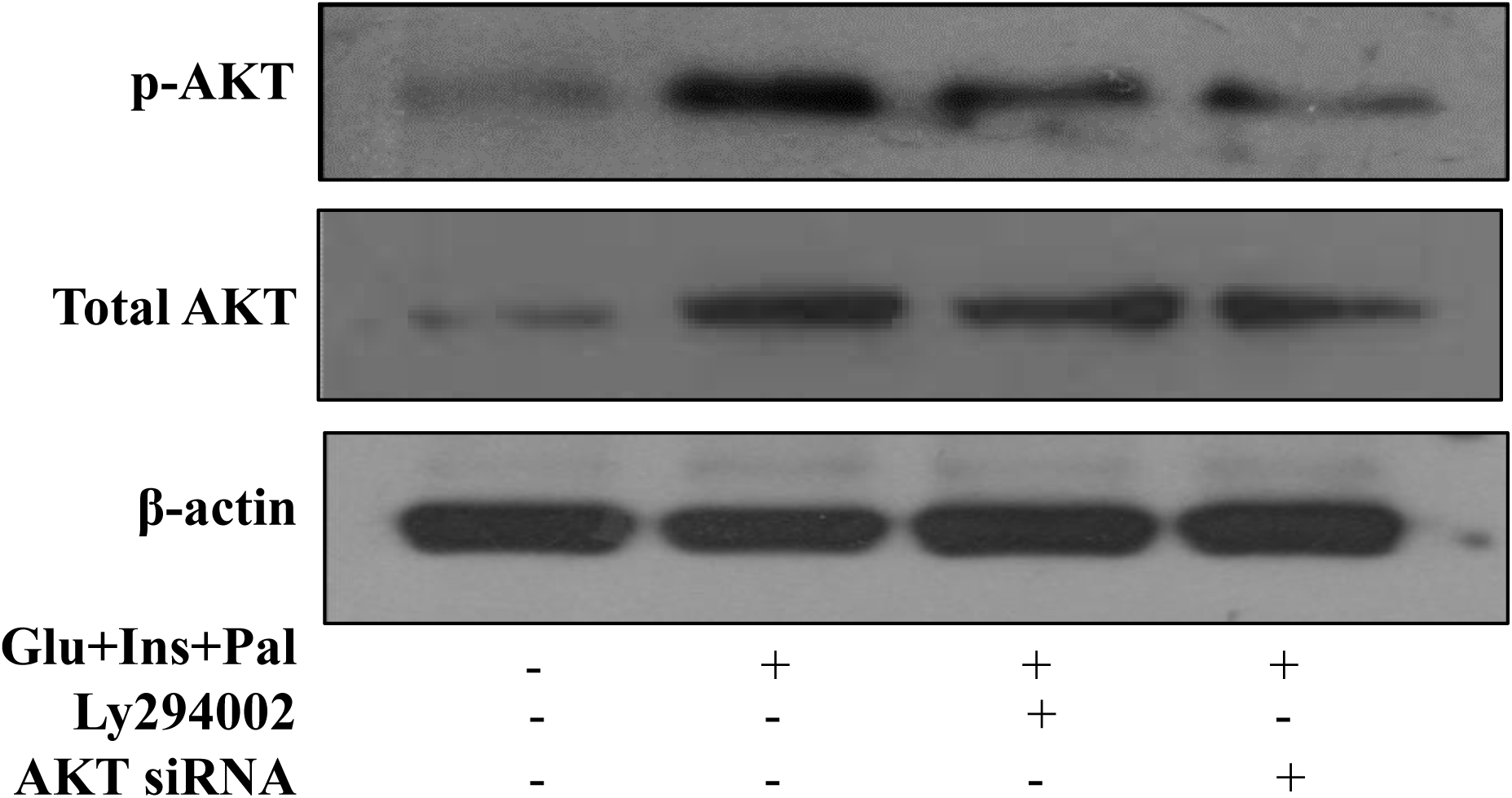
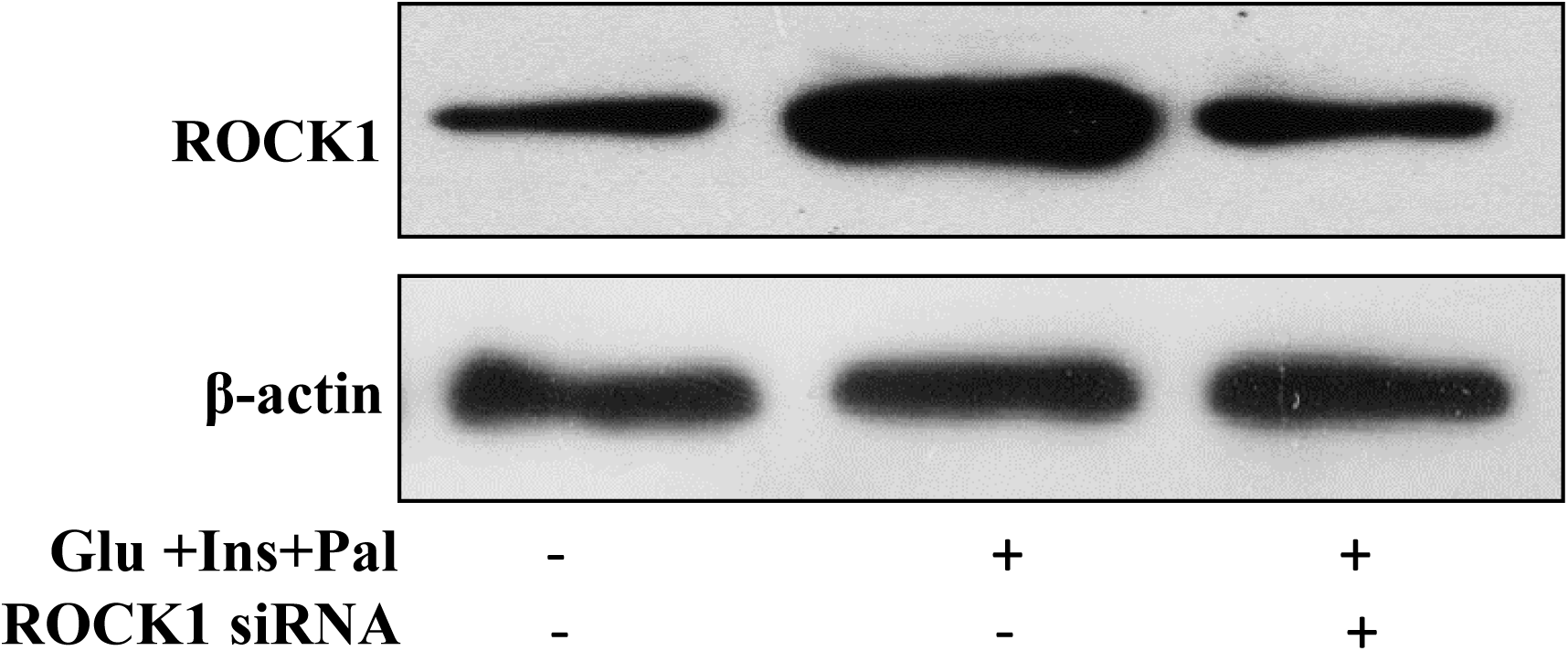
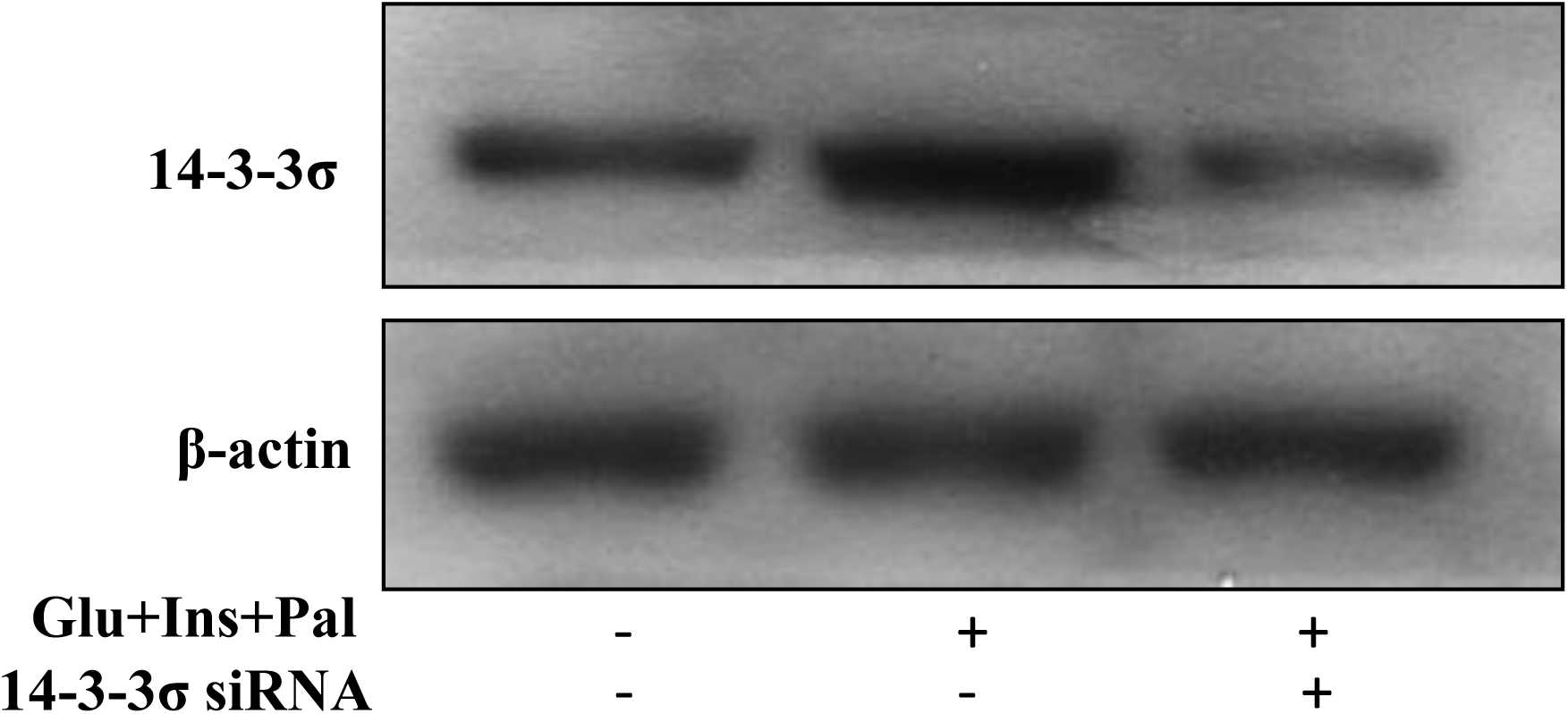
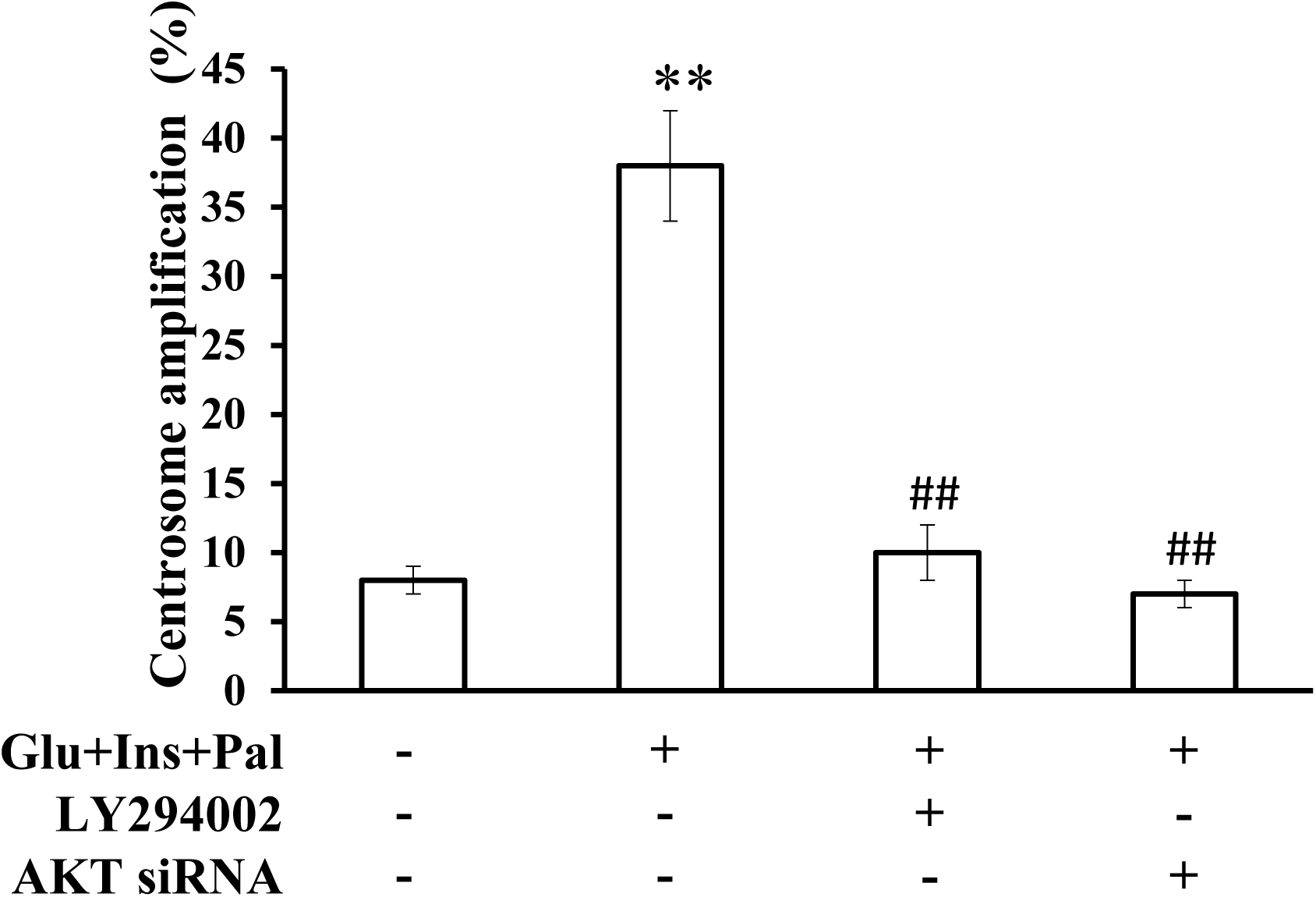
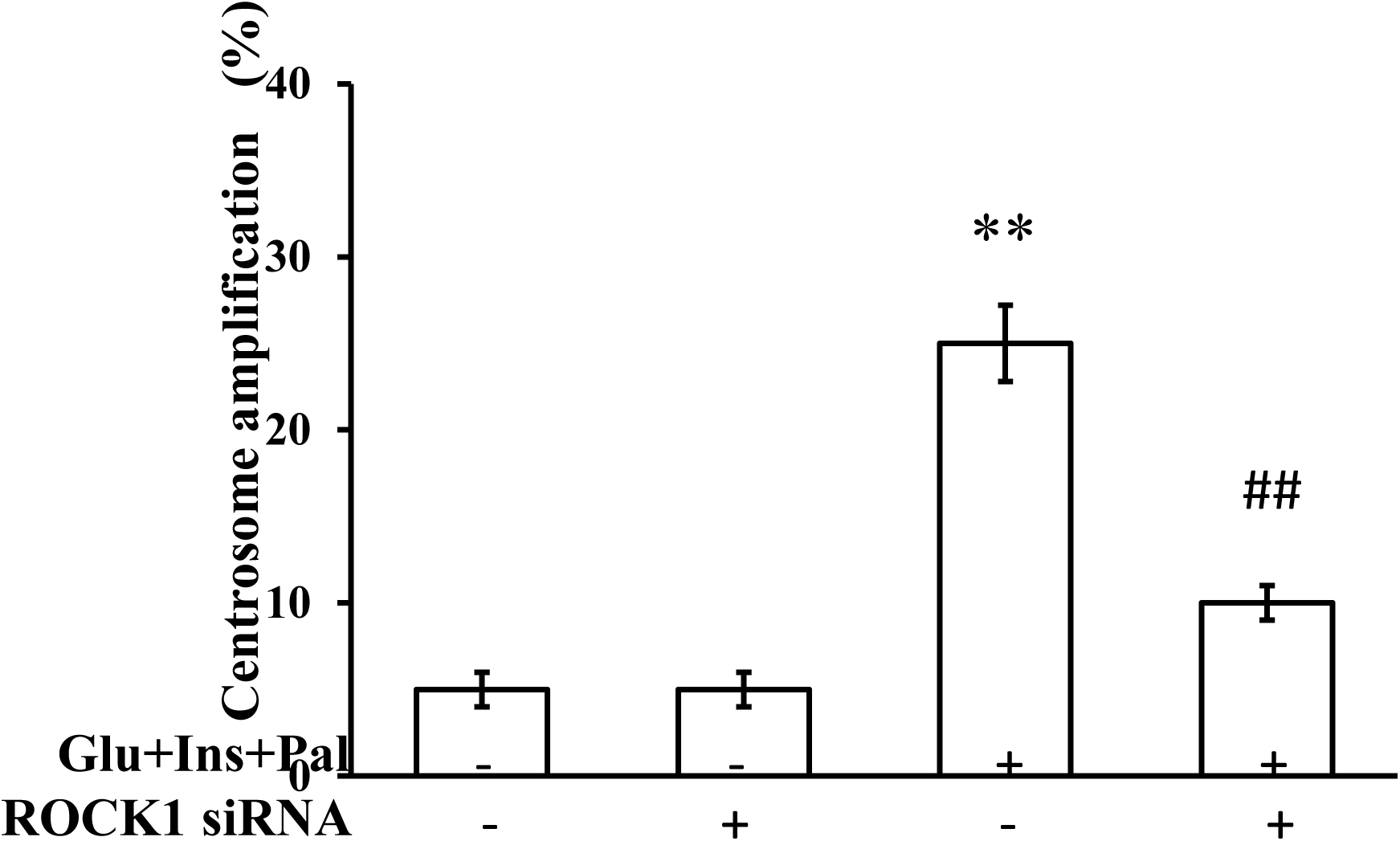
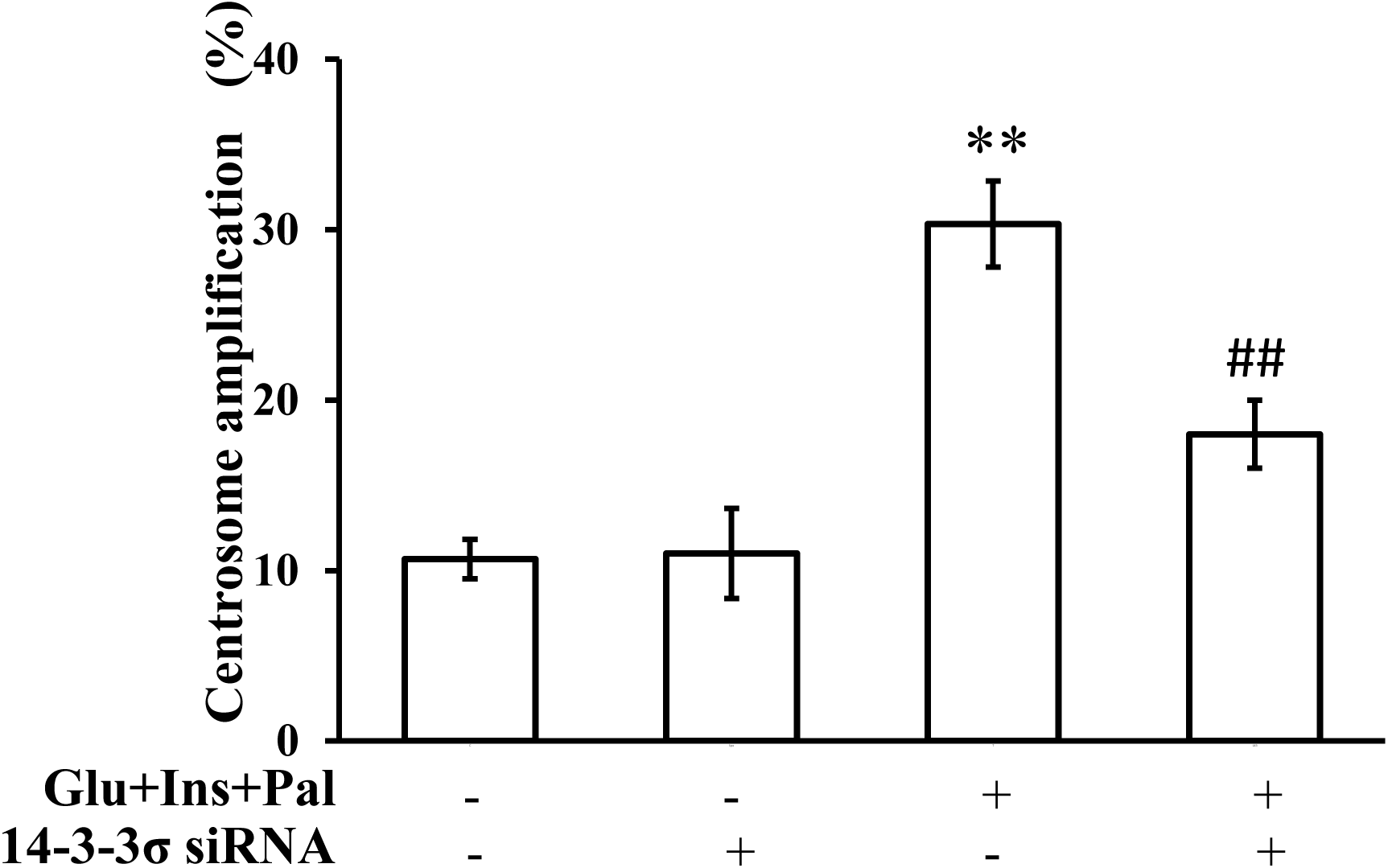
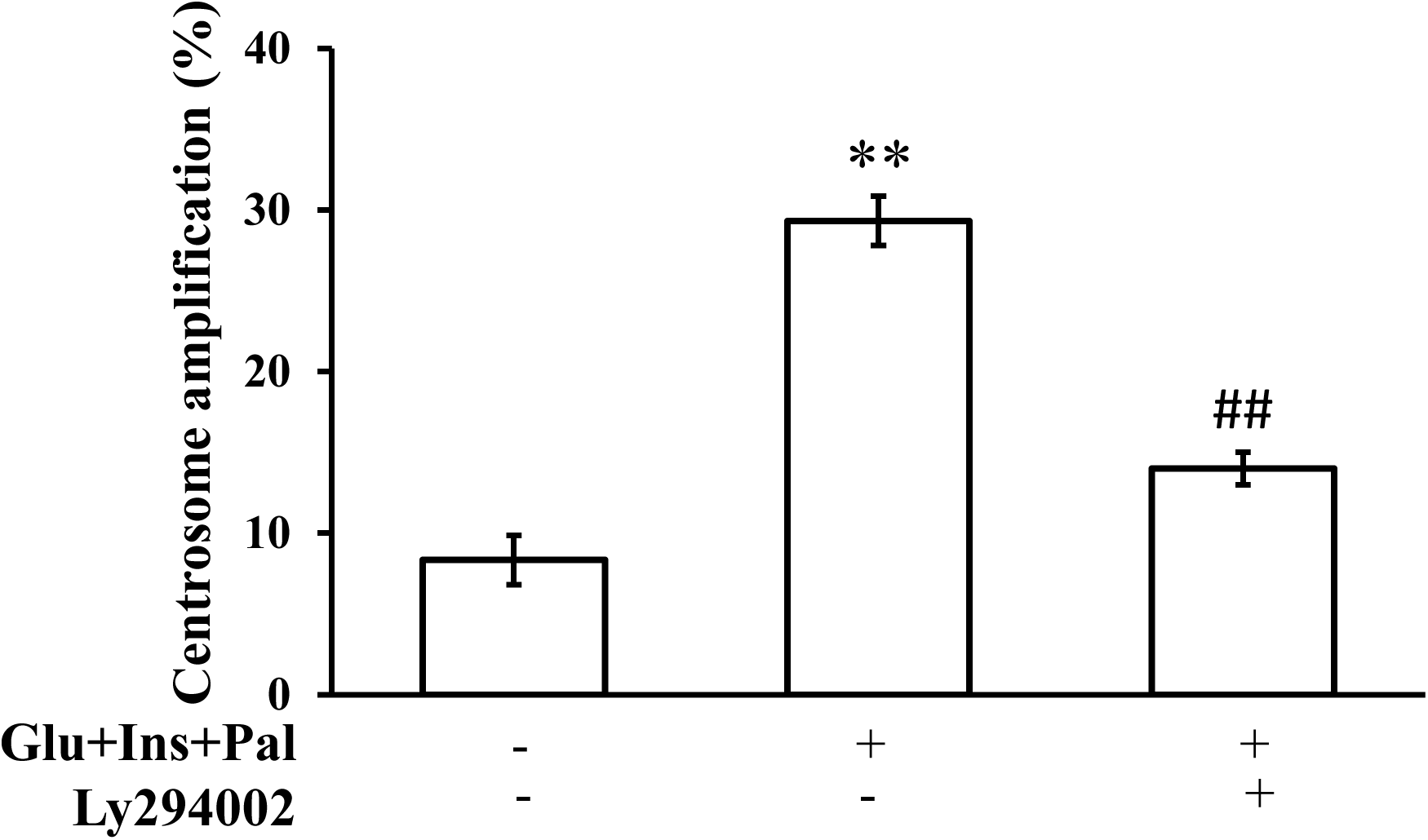
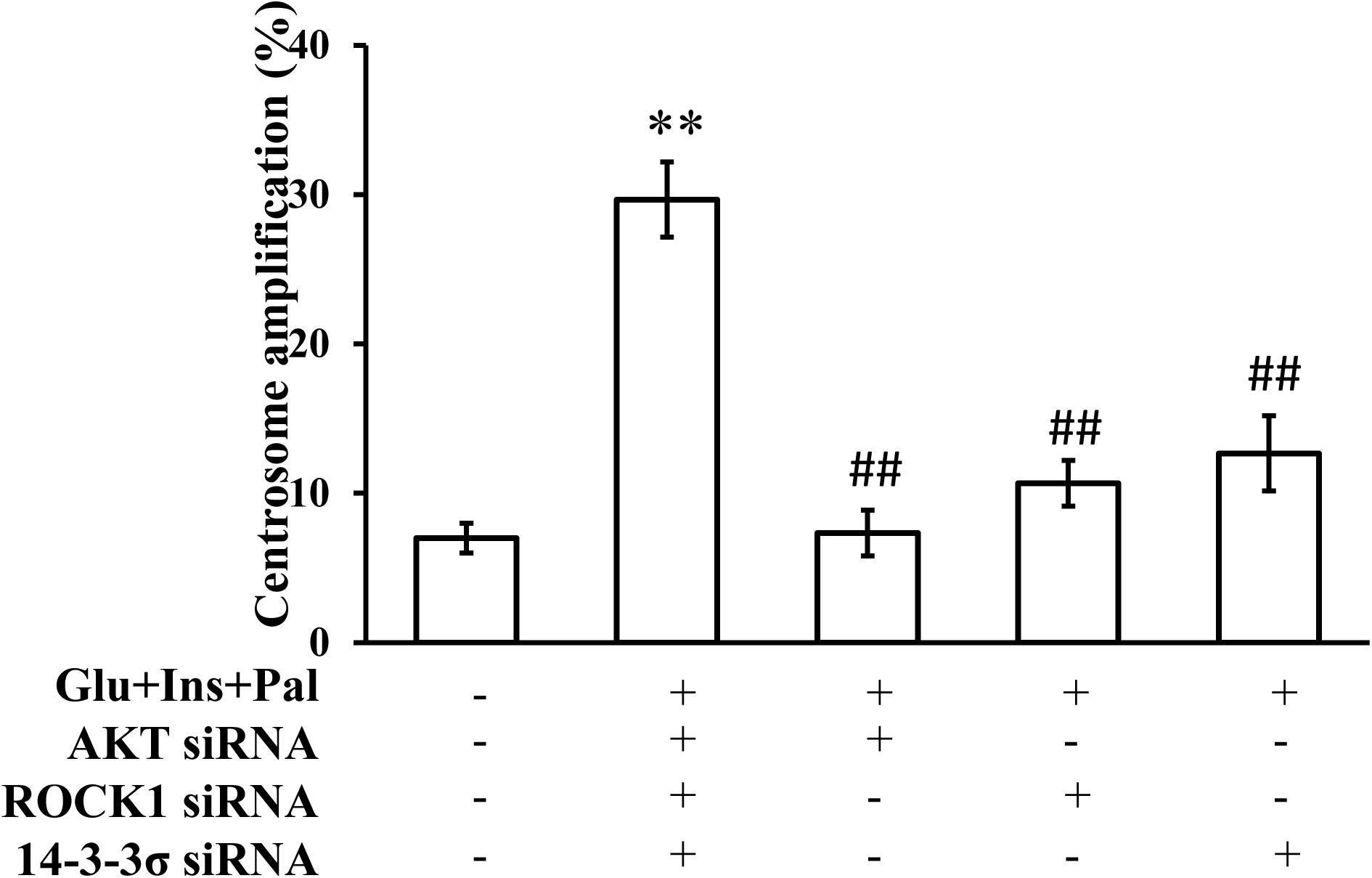
Functional transcriptomic analysis identifies AKT, ROCK1 and 14-3-3σ as signal mediators for the centrosome amplification. (**A**), (**B**) and (**C**):the diabetic pathophysiological factors activate of AKT, ROCK1 and 14-3-3σ, respectively; (**D**): inhibition of AKT using chemical inhibitor or siRNA blocks the centrosome amplification in the colon cancer cells; (**E**): ROCK1 siRNA inhibits the centrosome amplification; (**F**): 14-3-3σ siRNA inhibits the centrosome amplification in normal human breast epithelial cells. (G): AKT inhibitor inhibits centrosome amplification; (H): AKT siRNA, ROCK1 siRNA and 14-3-3σ siRNA inhibit centrosome amplification in normal human breast epithelial cells. One way ANOVA analysis was used to compare multiple groups.**: p < 0.01, compared with that in the control group; ##: p < 0.01, compared with that in the samples treated with Glu, Ins and Pal.Glu: glucose, 15 mM; Ins: insulin, 5 nM; Pal: palmitic acid, 150 μM; Ly294002: 30 μM.

### AKT-ROS-dependent signaling of ROCK1 and 14-3-3σ is the molecular pathway for the centrosome amplification

We next delineated the signal transduction pathway for the centrosome amplification in the colon cancer cells. The logic was that inhibition of an upstream signal would inhibit downstream one(s), while inhibition of a downstream signal would not affect the upstream one(s). We observed that inhibition of AKT using chemical inhibitor (Fig. 4A) orsiRNA (Fig. 4B) was able to inhibit ROS production as well as the upregulations of ROCK1 (Figs. 4C) and 14-3-3σ (Fig. 4D). NAC was able to inhibit the upregulation of ROCK1 (Fig. 4E) and 14-3-3σ (Fig. 4F), but did not affect the activation of AKT (figure not shown). ROCK1 or 14-3-3σ siRNA did not affect AKT activation or ROS production (figures not shown).

**Fig. 4.**
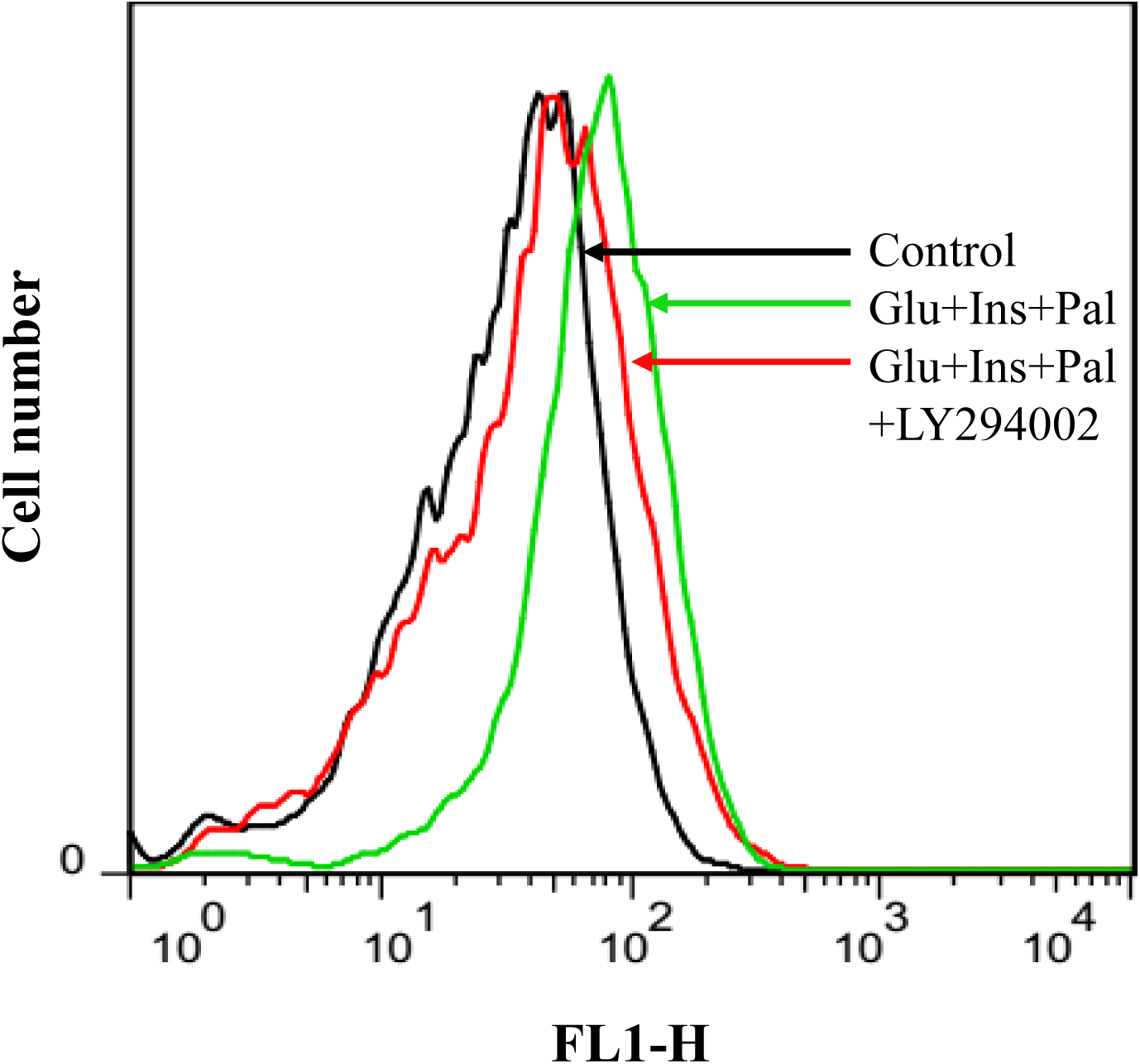
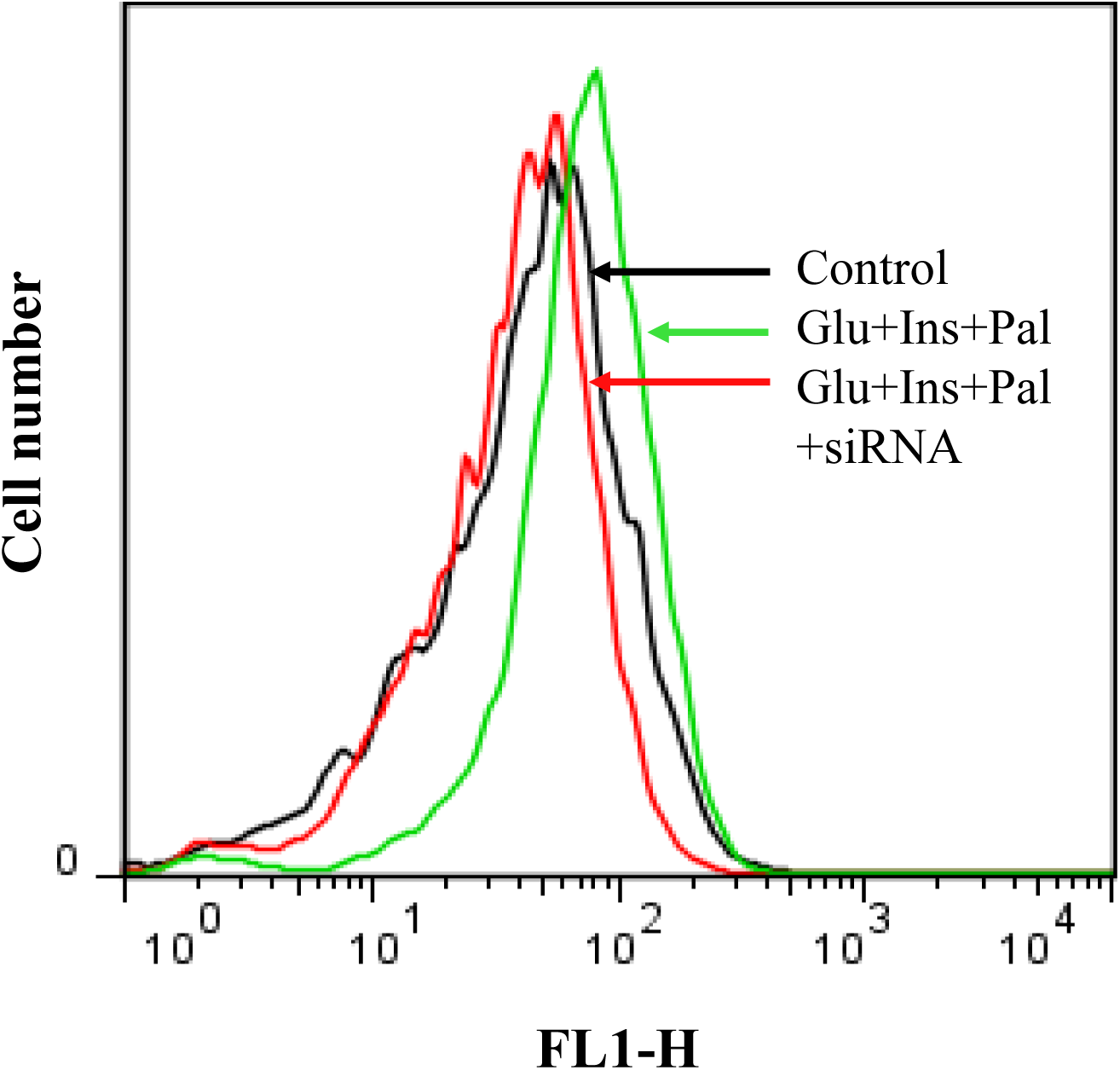
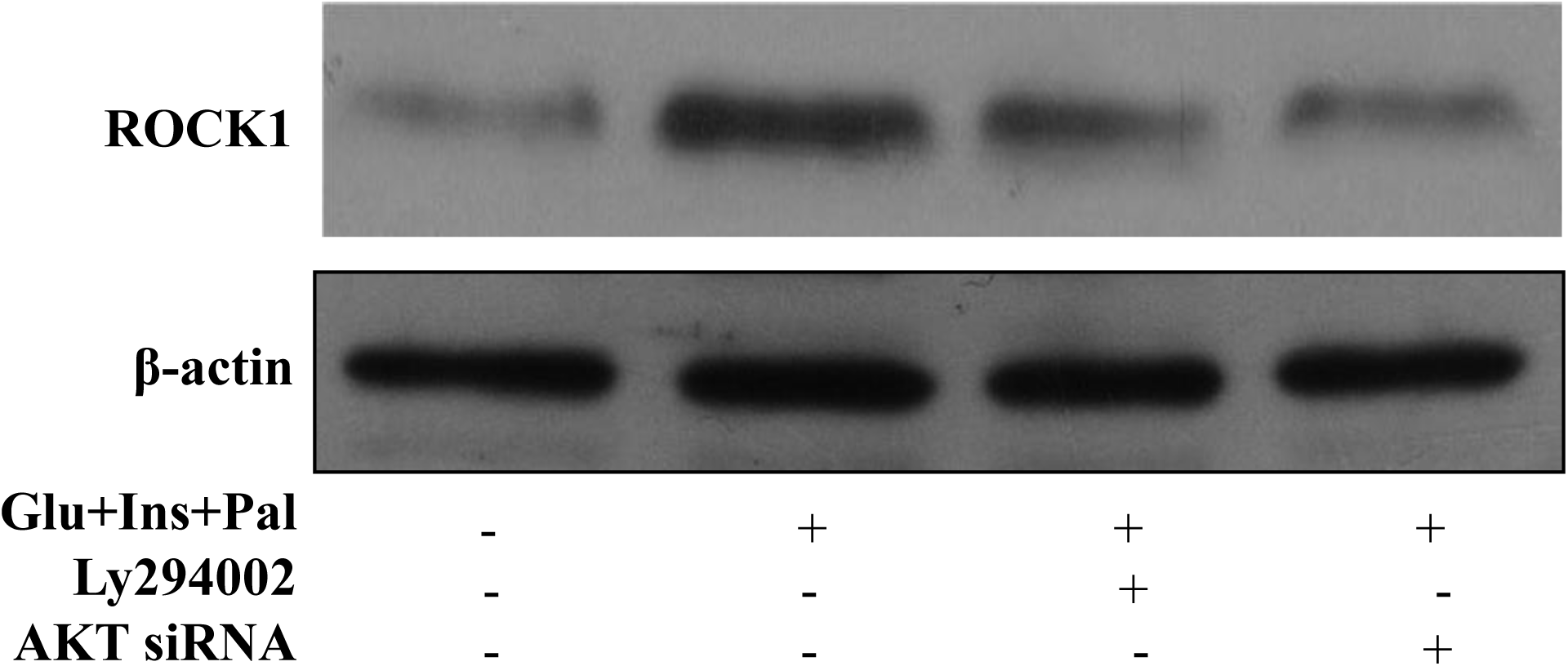
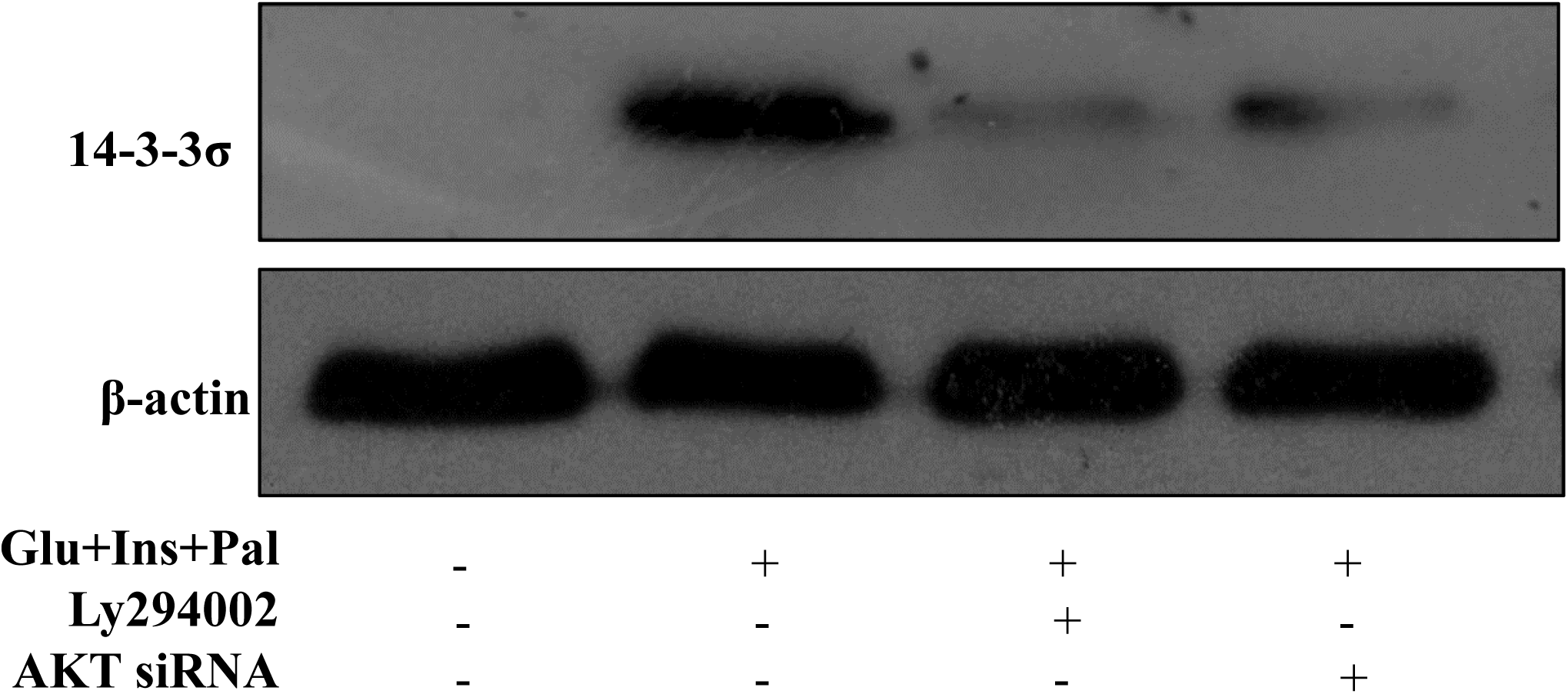
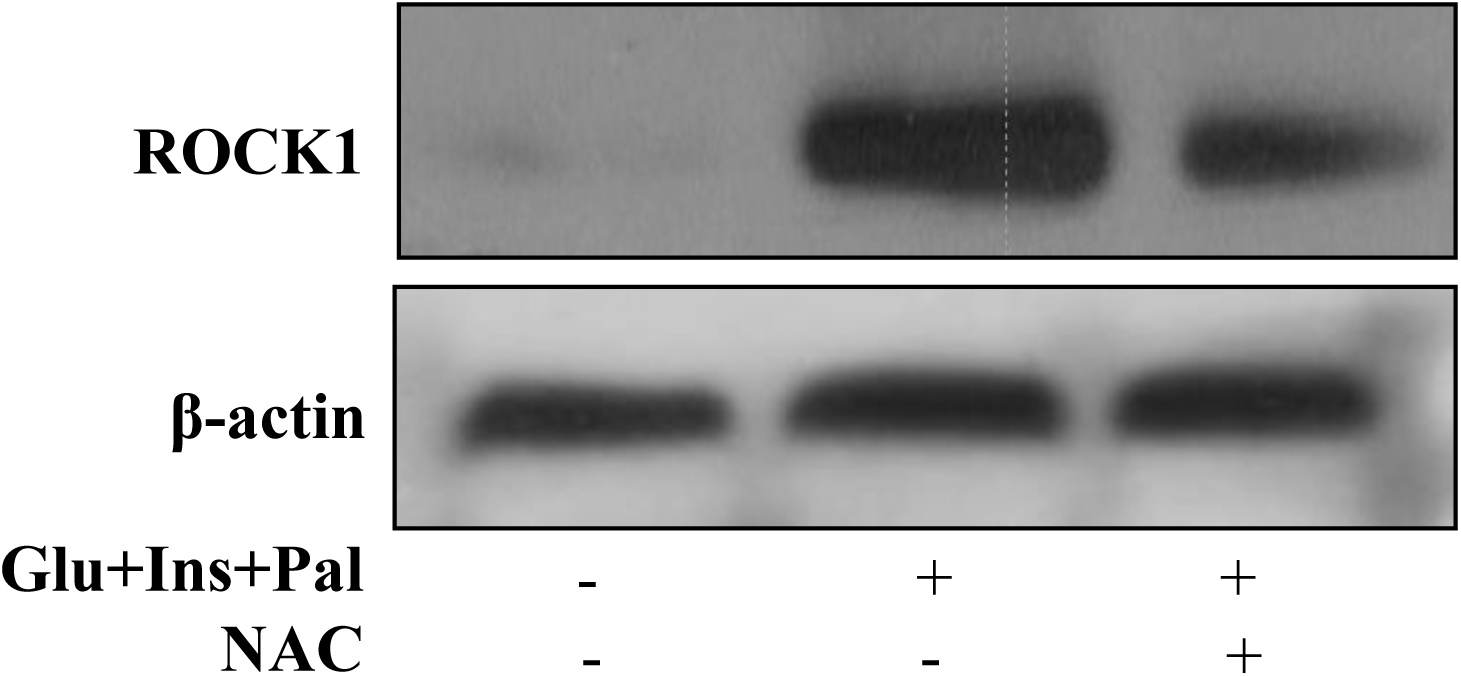
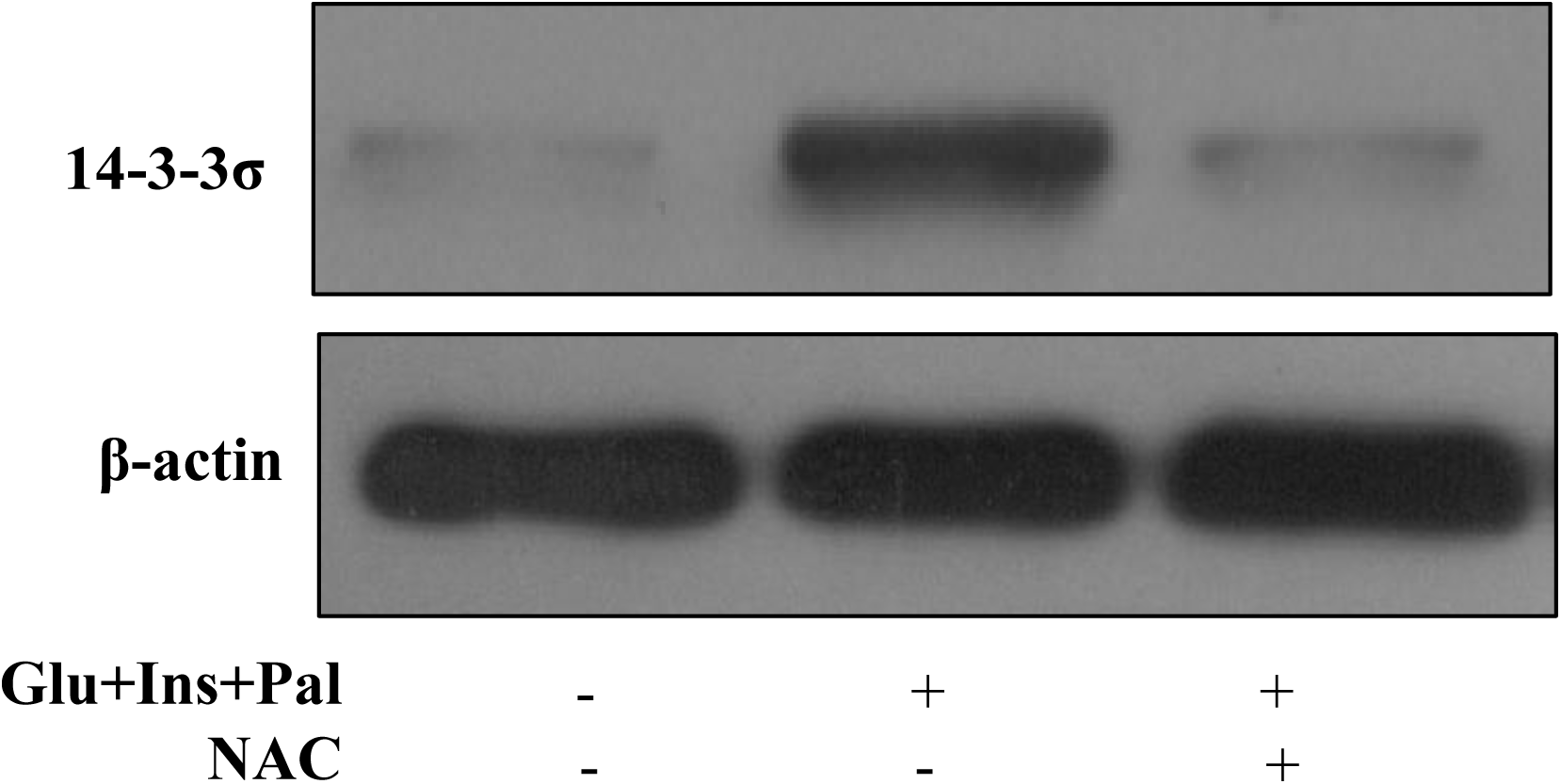
AKT-ROS-dependent signaling of ROCK1 and 14-3-3σ is the molecular pathway for the centrosome amplification. (**A**) and (**B**): AKT chemical inhibitor and siRNA inhibit ROS production, respectively; (**C**): AKT chemical inhibitor or siRNA inhibits ROCK1 upregulation; (**D**): AKT chemical inhibitor or SiRNA inhibits 14-3-3σ upregulation; (**E**) and (**F**): NAC inhibits the upregulation of ROCK1 and 14-3-3σ, respectively. Glu: glucose, 15 mM; Ins: insulin, 5 nM; Pal: palmitic acid, 150 µM; Ly294002, 30 μM; NAC: 3 mM.

### High glucose, insulin and palmitic acid increase the binding between ROCK1 and 14-3-3σ as well as their centrosome translocation

Next, we investigated the relationships amongst ROCK1, 14-3-3σ and centrosome in the cancer cells by using ROCK1 and 14-3-3σ antibodies to pull down their binding partners, as well as to analyse the centrosome translocation of the two proteins. In pull-down samples using ROCK1 antibody, 14-3-3σ protein was detected, which was increased in the cells treated with high glucose, insulin and palmitic acid (Fig. 5A). Similarly, high glucose, insulin and palmitic acid increased the pull down ofROCK1 using 14-3-3σ antibody (Fig. 5A). AKT inhibitor and NAC were able to inhibit the increase in co-precipitation of ROCK1 and14-3-3σ from the cells treated by high glucose, insulin and palmitic acid (Figs. 5A). These observations on the Western blot analysis images were confirmed when the images were quantified (Figs. 5B and 5C. As shown in confocal images in Figs. 5D and 5E respectively, ROCK1 and 14-3-3σ were localized in the centrosome in all the treated samples. In the cells with more than one centrosome, only half of the centrosomes had ROCK1, with another half of centrosomes free of ROCK1 (Fig. 5D). The fluorescent intensity of ROCK1 staining, which indicates the level of ROCK1 in the centrosomes, was higher in the treated cells compared to that in the control cells, which was inhibited by AKT inhibitor or NAC but not 14-3-3σ siRNA (Fig. 5F). Comparing with that only 1% of the control cells had 14-3-3σ in the centrosomes, approximately 20% of the cells treated with high glucose, insulin and palmitic acid had 14-3-3σ in the centrosomes, which was inhibited by AKT inhibitor, NAC or ROCK1 siRNA (all at p< 0.01; Figs. 5E and 5G).

**Fig. 5.**
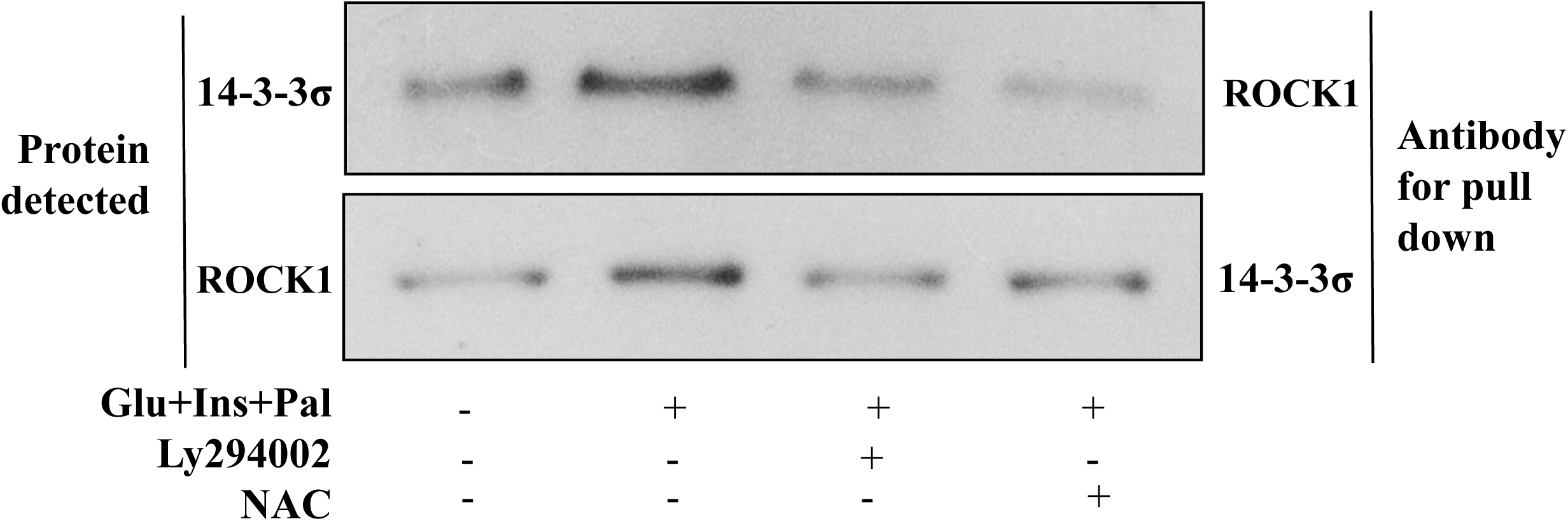
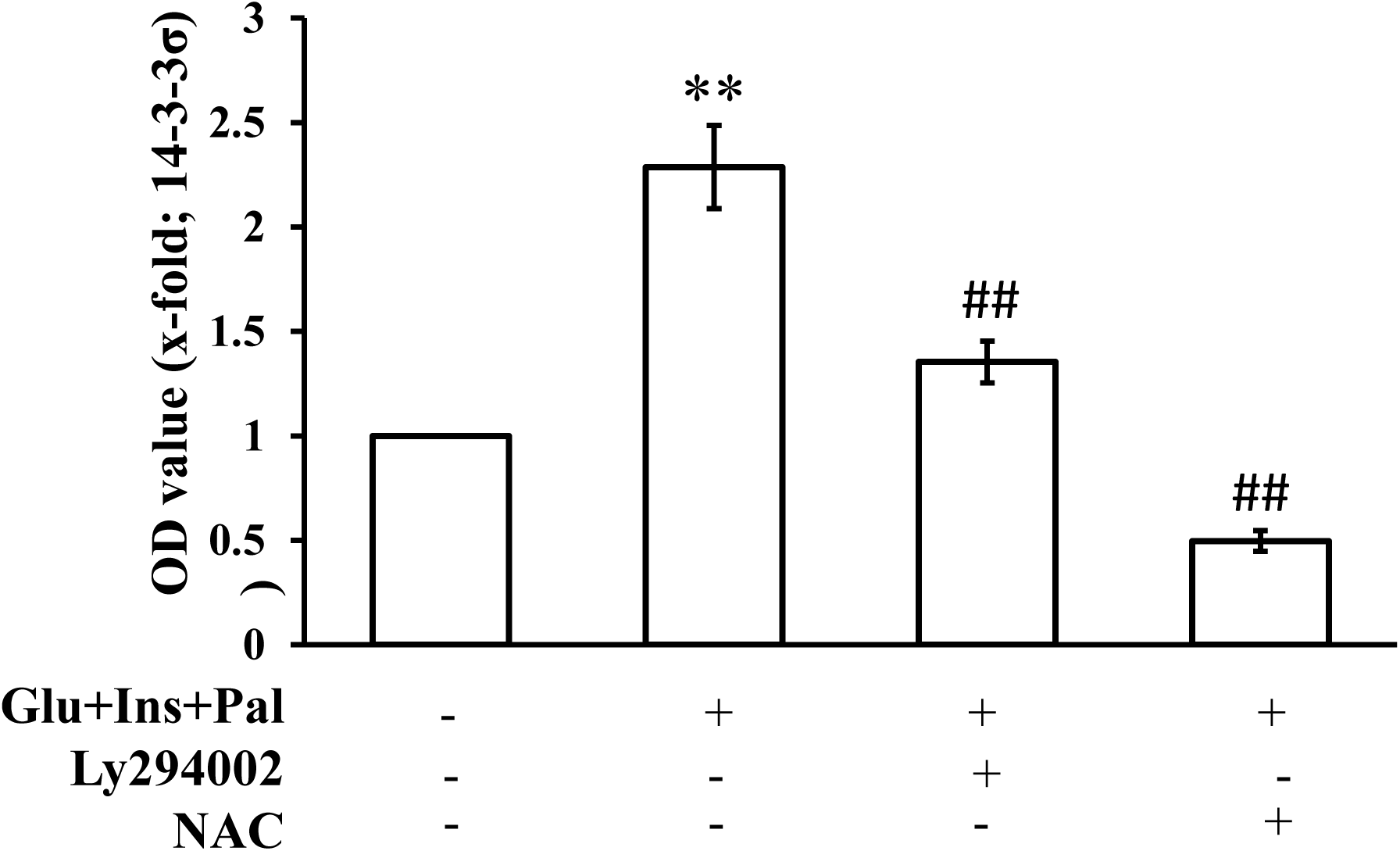
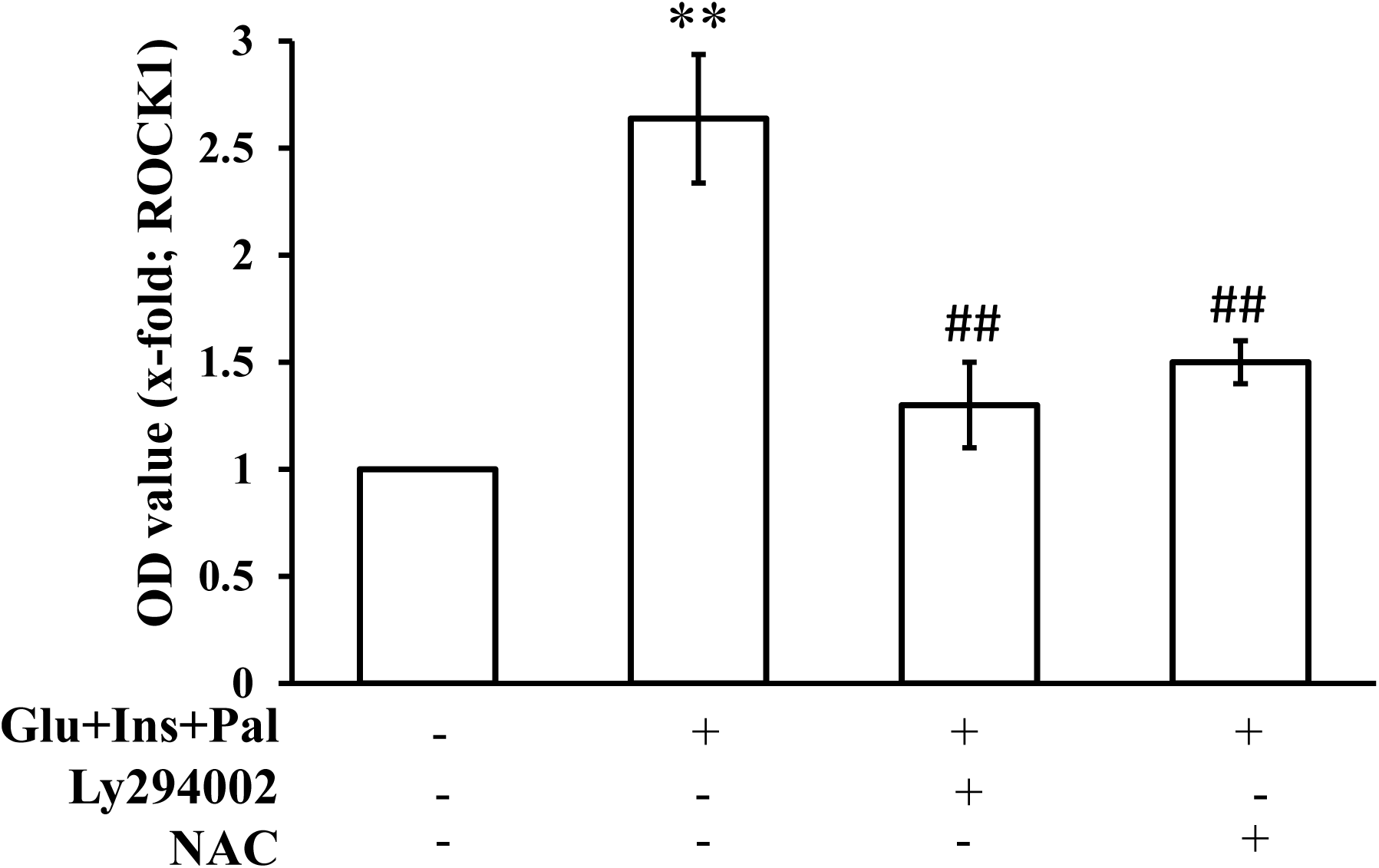
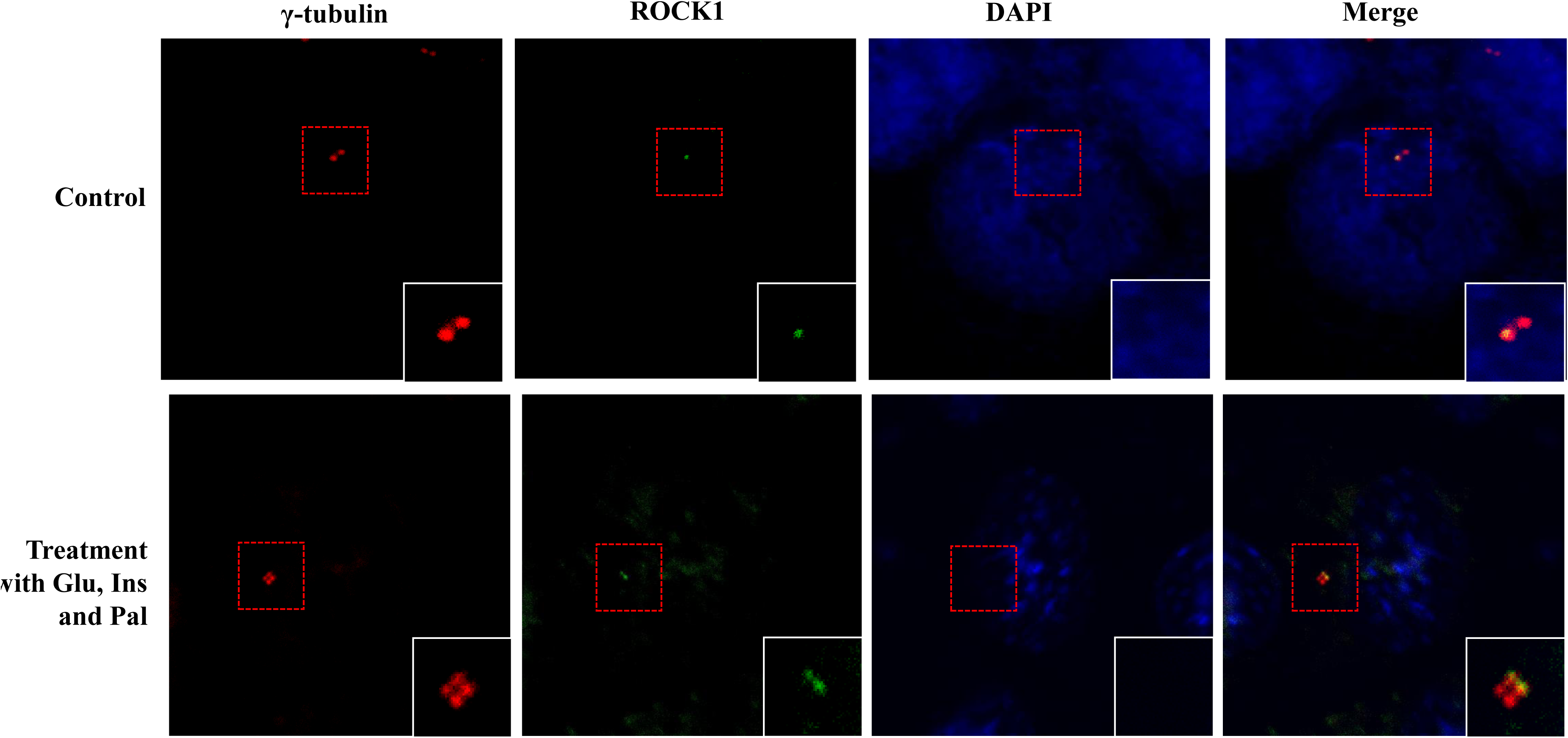
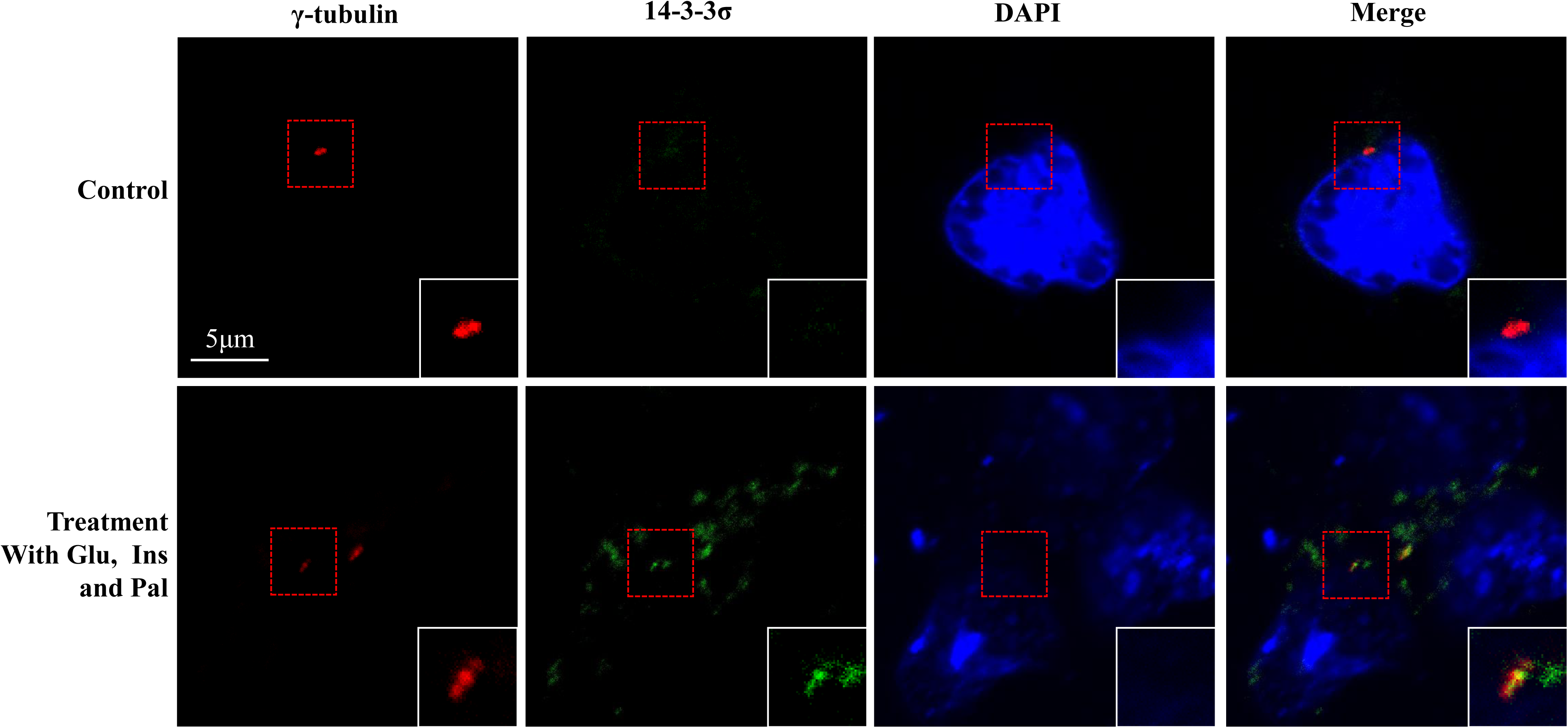
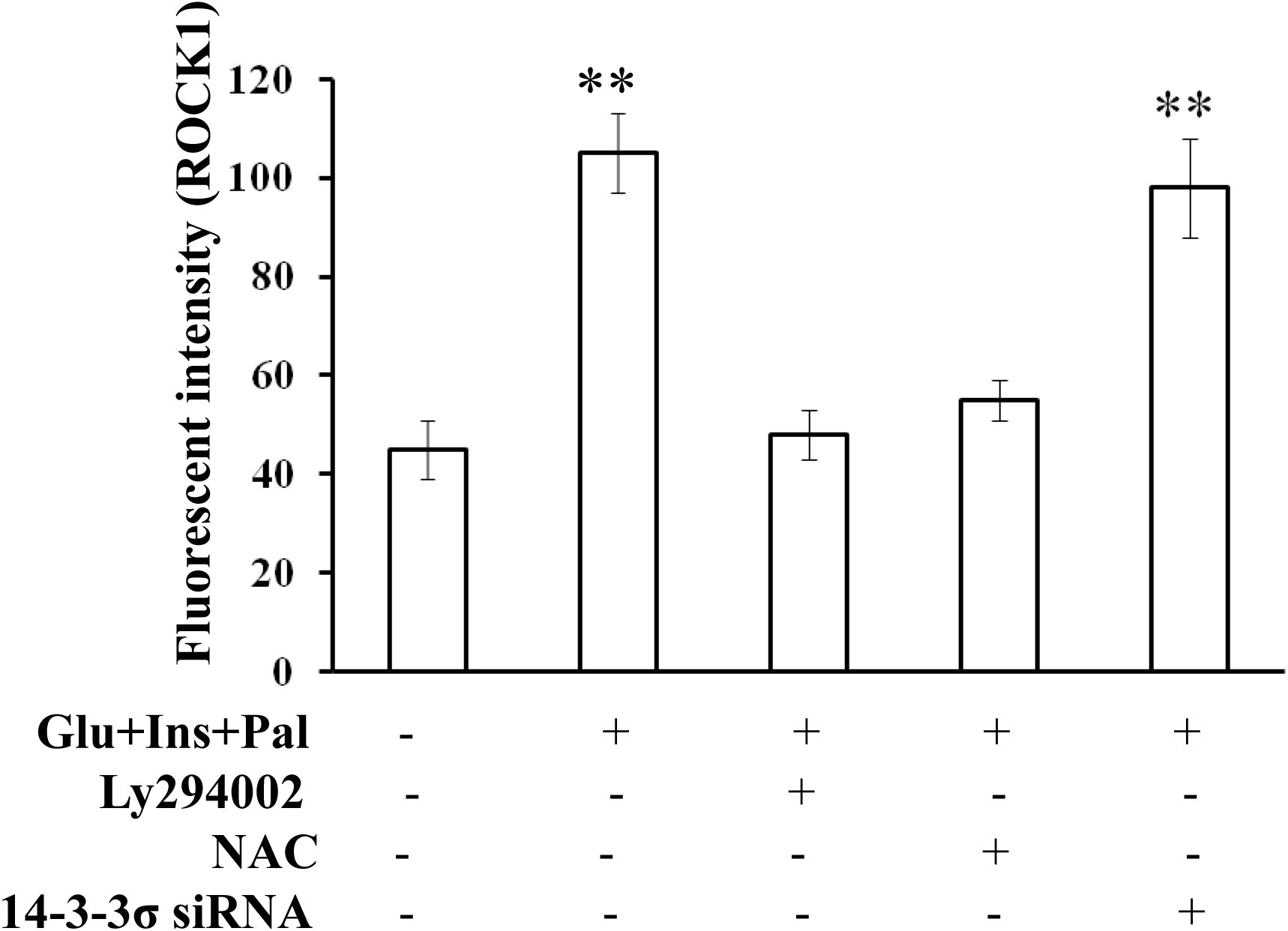
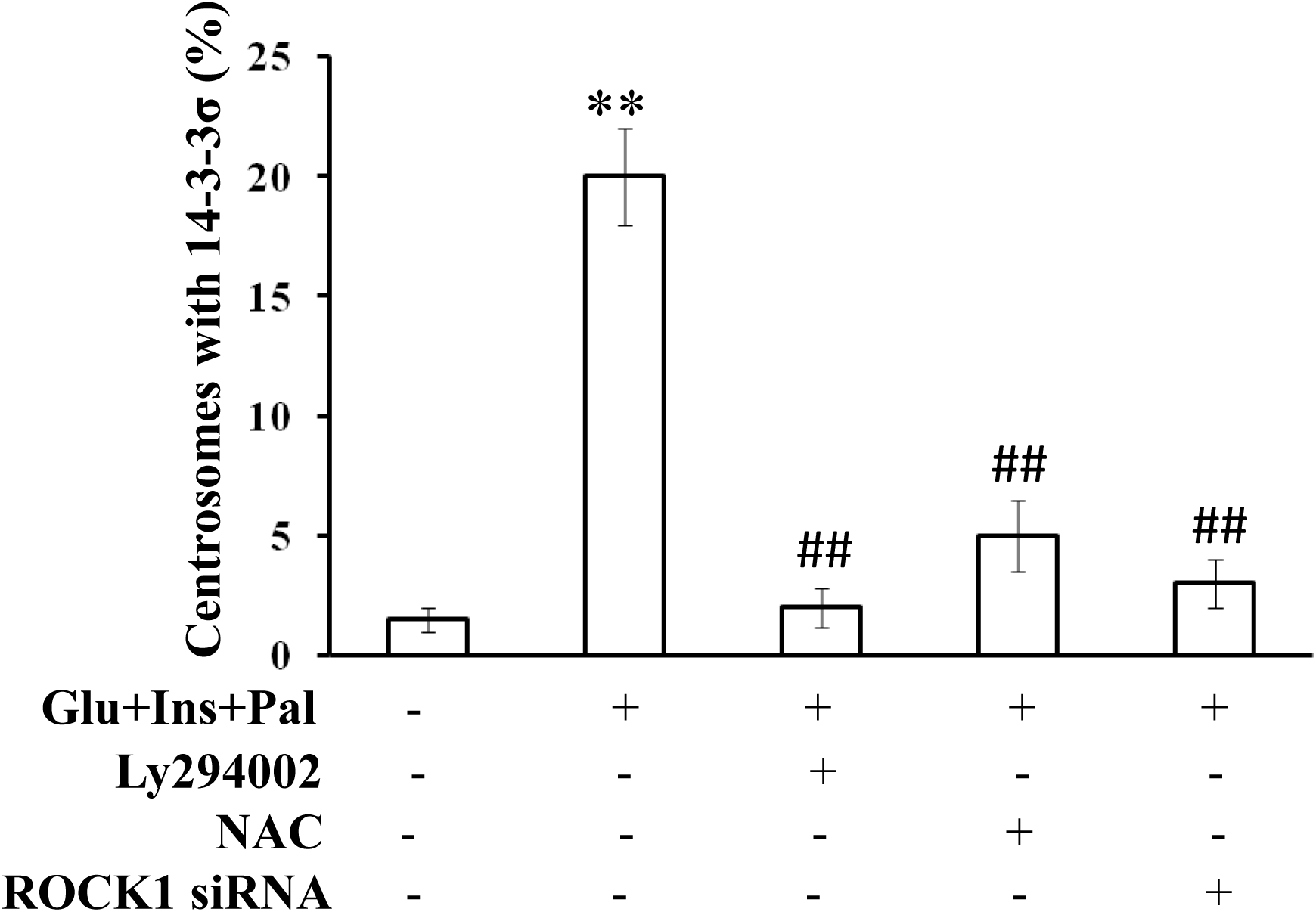
High glucose, insulin and palmitic acid increase the binding between ROCK1 and 14-3-3σ as well as their centrosome translocation. (**A**): use of ROCK1 or 14-3-3σ antibodies to pull down their binding partner, and the influences of Ly294002 and NAC; (**B**) and (**C**): quantification of the gel images of Western blot analysis shown in Fig. 5A; (**D**): confocal image of ROCK1 localization in the centrosomes; (**E**): confocal image of 14-3-3σ localization in the centrosomes; (**F**): high glucose, insulin and palmitic acid increases level of ROCK1 localization in the centrosomes, which is inhibited by AKT inhibitor and NAC but not 14-3-3σ siRNA; (**G**): experimental treatment increases the localization of 14-3-3σ to the centrosomes, which is inhibited by AKT inhibitor, NAC or ROCK1 siRNA. One way ANOVA analysis was used to compare multiple groups.**: p < 0.01, compared with that in the control group; ##: p < 0.01, compared with that in the samples treated with Glu, Ins and Pal.Glu: glucose, 15 mM; Ins: insulin, 5 nM; Pal: palmitic acid, 150 μM; Ly294002, 30 μM; NAC, 3 mM.

### The increases in AKT activation and ROS production as well as expression, binding and centrosome distribution of ROCK1 and 14-3-3σ *in vitro* are verified in the PBMC from the volunteers

Finally, we investigated whether the molecular signalling events found *in vitro* occurred *in vivo* in patients with type 2 diabetes. We compared these molecular events between the PBMC from healthy volunteers and those from type 2 diabetic patients. The results showed that PBMC from the diabetic patients had increased level of AKT activation (Fig. 6A) and ROS production (Fig. 6B). The PBMC from the patients also had increased expression (Fig. 6A), binding (Figs. 6C) and centrosome distribution of ROCK1 and 14-3-3σ (Figs. 6D and 6E).

**Fig. 6.**
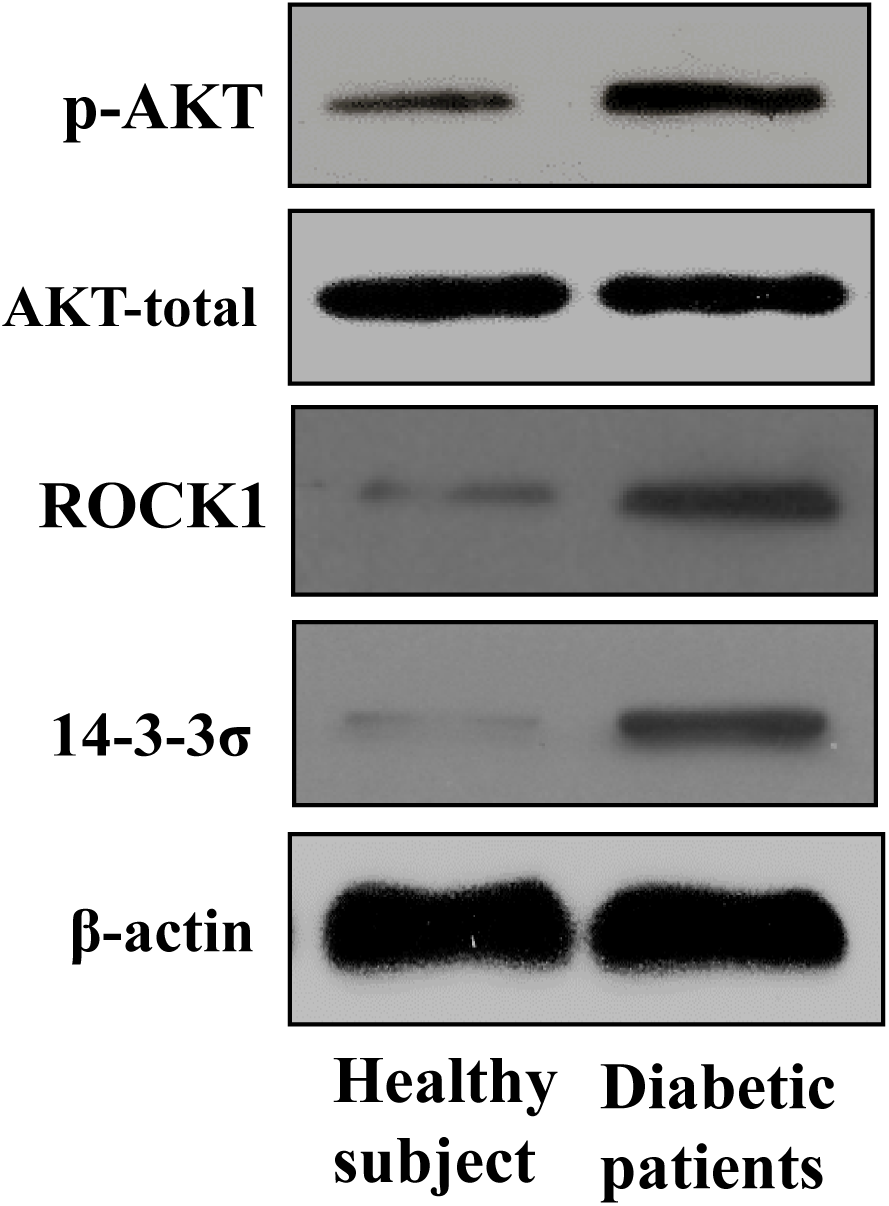
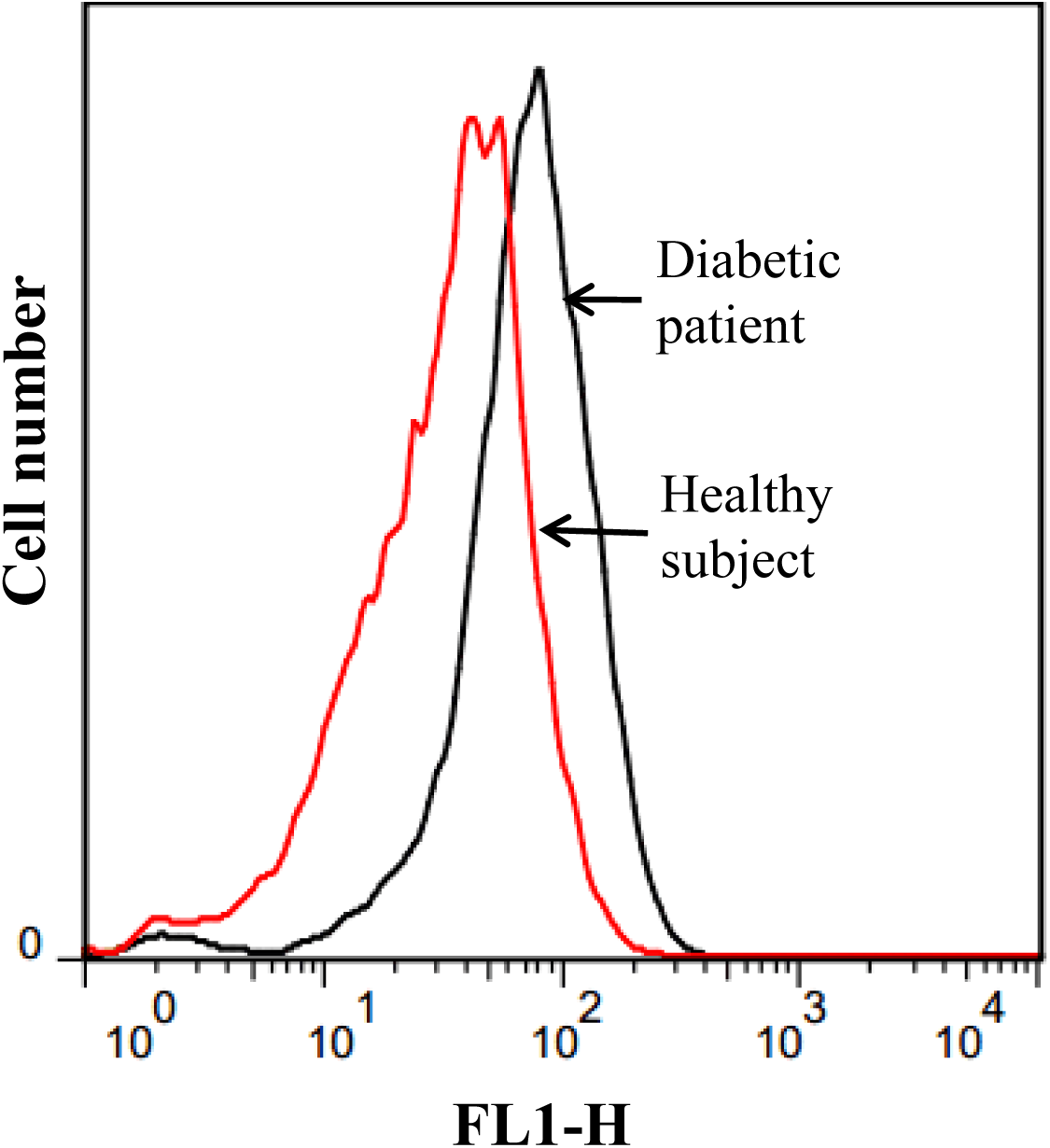
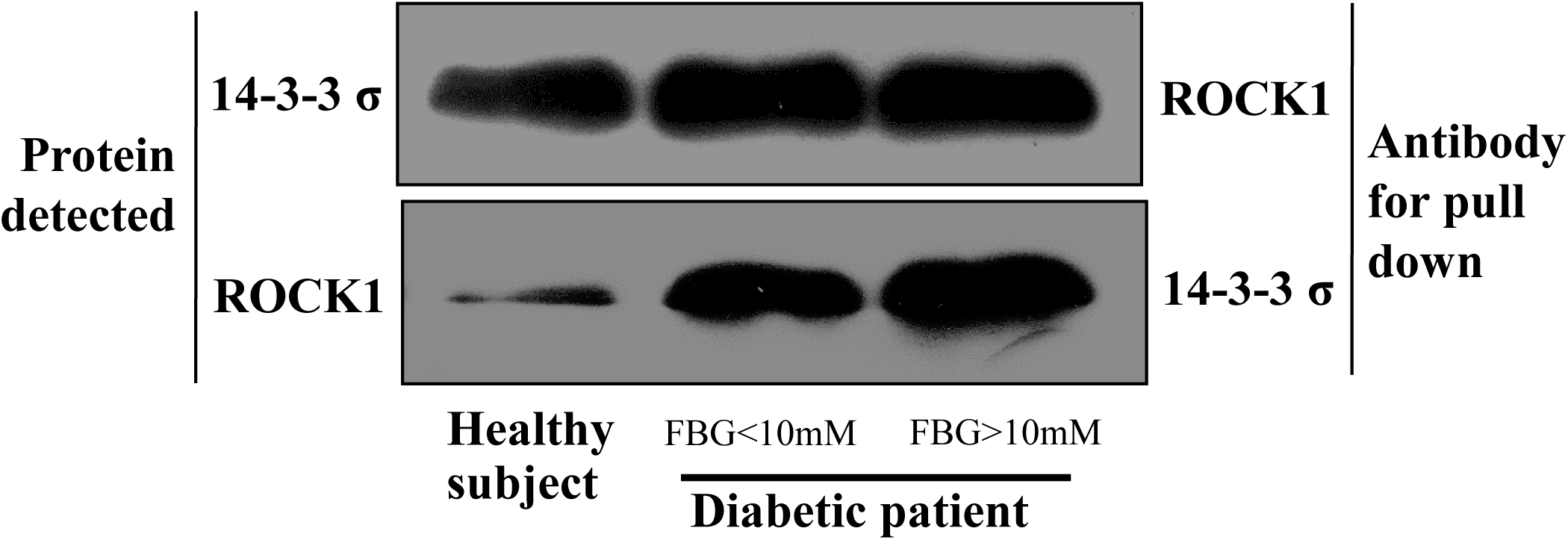
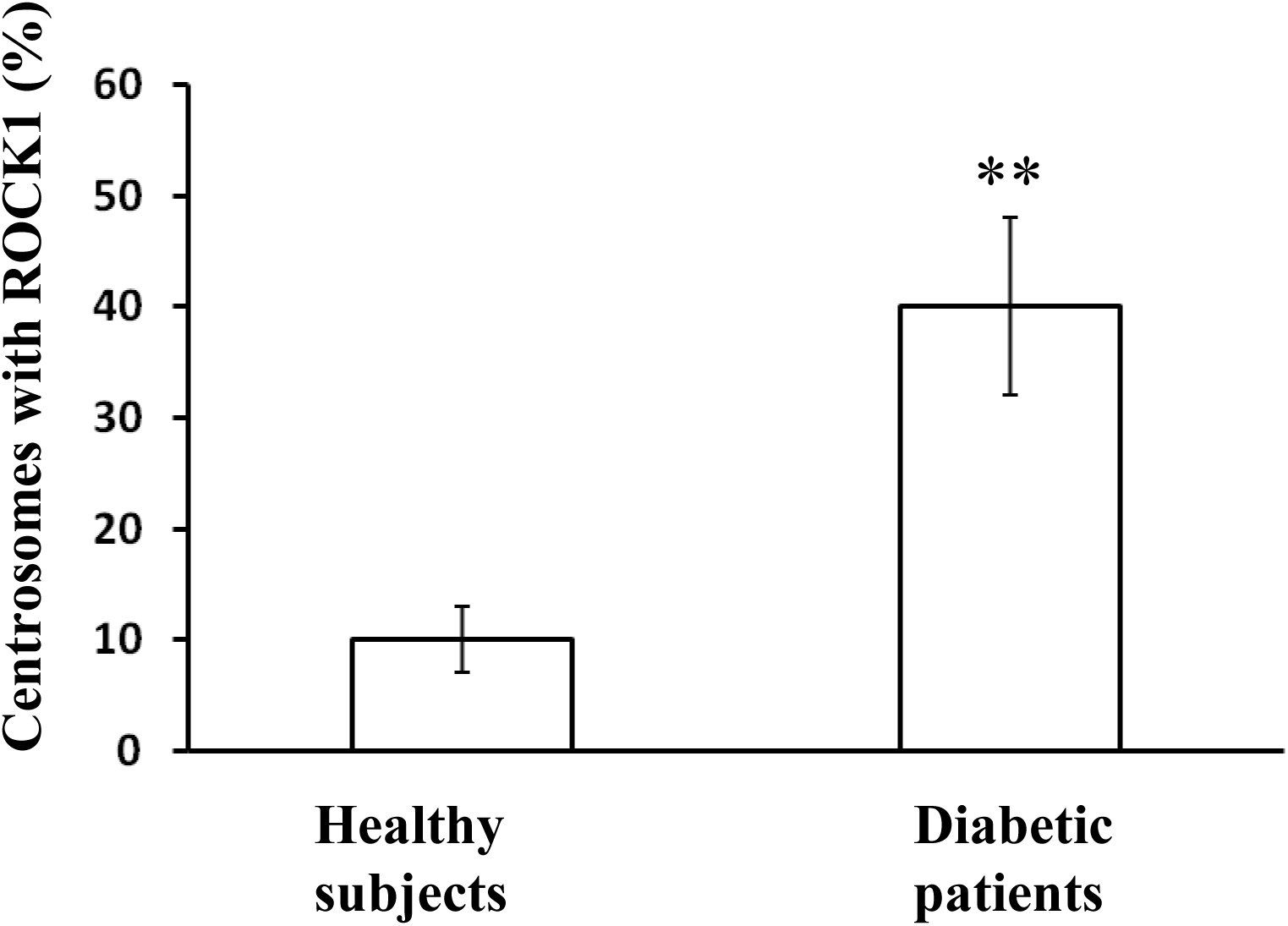
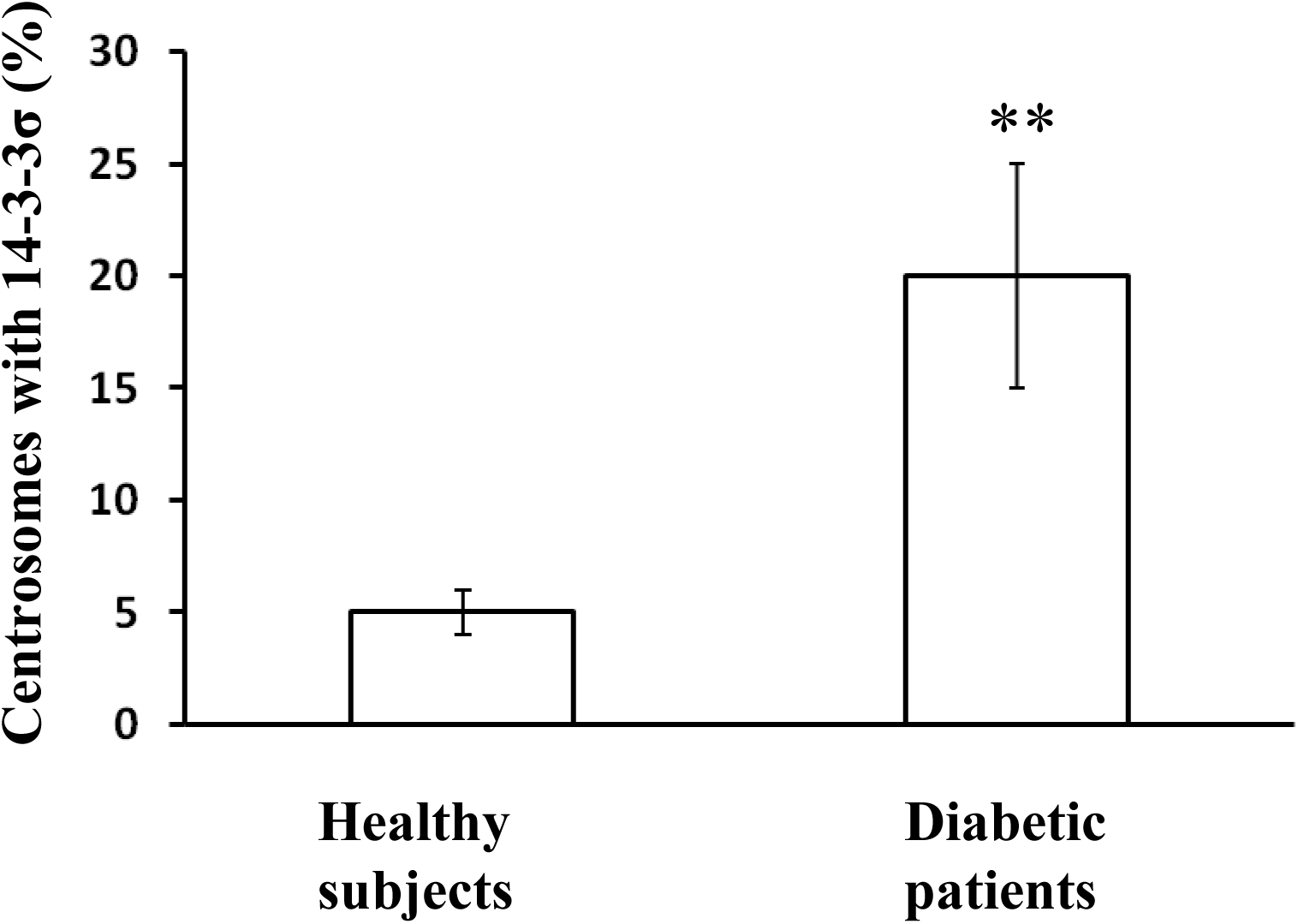
The increases in AKT activation and ROS production as well as expression, binding and centrosome distribution of ROCK1 and 14-3-3σ are confirmed in the PBMC from the patients with type 2 diabetes. (**A**): AKT activation as well as expressions of ROCK1 and 14-3-3σ are increased in PBMC from the patients; (**B**): ROS production is increased in the PBMC from the patients; (**C**): binding between ROCK1 and 14-3-3σ is increased in PBMC from the diabetic patients; (**D**) and (**E**): centrosome distributions of ROCK1 and 14-3-3σ are increased in the PBMC from the type 2 diabetic patients, respectively. For Western blot analyses and co-immunoprecipitation, five clinical samples were randomly selected and combined for the experiments. ROS quantification was performed in triplicate using different samples. One way ANOVA analysis was used to compare multiple groups. **: p < 0.01.

## DISCUSSION

Our results showed that patients with type 2 diabetes had 2.6-fold increase in cell centrosome amplification *in vivo* in PBMC, which correlated with poor glycemic control (Fig. 1). Pathophysiological factors in type 2 diabetes, i.e., high glucose, insulin and palmitic acid, could trigger centrosome amplification (Figs. 2C and 3D). AKT-ROS-dependent upregulation of expression, binding and centrosome translocation of ROCK1 and 14-3-3σ was the underlying molecular pathway (Figs. 3-6).

Type 2 diabetes increases the risk of developing all-site cancer, with the exception of prostate cancer (Giovannucci et al., 2010). Cancer patients with type 2 diabetes have poorer prognosis than those without diabetes (Mills et al., 2013). However, it remains unknown why and how type 2 diabetes favours cancer development. Centrosome amplification can initiate in animal model (Levine et al., 2017) and increase cancer cell invasion potential (Godinho et al., 2014). In the present study, we showed that patients with type 2 diabetes had increased level of cell centrosome amplification *in vivo* (Fig. 1). Thus, we speculate whether there is a link between the diabetes-associated centrosome amplification (Fig. 1) and the increased cancer risk in type 2 diabetes.

Our results (Figs. 3-6) support the findings that AKT (Na et al., 2015) and ROS (Chae et al., 2005) can mediate centrosome amplification, and further place AKT upstream of ROS. In an apoptosis model, activation of ROCK1 causes AKT inactivation (Zhang et al., 2016), which is different from our results that placed ROCK1 downstream of AKT activation (Figs. 4C). Our result (Fig. 4E) showed that ROS was upstream of ROCK1, which is in agreement with the observations by Shen and Wang (Shen and Wang, 2015). Oh and Jang showed that AKT can be upstream of 14-3-3σ (Oh and Jang, 2009), which is supported by our result (Figs. 4D) that AKT was upstream of 14-3-3σ.

As shown in Fig. 4, inhibition of AKR inhibits all other signals. Inhibition of ROS inhibits ROCK1 and 14-3-3σ but not AKT. Inhibition of ROCK1 or 14-3-3σ did not affect any other signals. This suggests that AKT-ROS-dependent signalling of ROCK1 and 14-3-3σ is the pathway underlying the centrosome amplification by the experimental treatment. Our results (Figs. 5D and 5E) showed that ROCK1, but not 14-3-3σ, was a centrosome protein. ROCK1 was present in half of the centrosomes when a cell had two or more centrosomes. Whether it is present only in the mother centrosomes remains to be clarified. In most cases, 14-3-3σ was present in the centrosomes only in the treated cells (Fig. 5E). Knockdown of ROCK1 inhibited 14-3-3σ translocation to centrosome, while knockdown of 14-3-3σ did not affect the translocation of ROCK1 to the centrosomes (Figs. 5F and 5G), which suggests that ROCK1 transports 14-3-3σ to centrosome or is involved in 14-3-3σ translocation to centrosome after treatment. Downregulation of either ROCK1 or 14-3-3σ inhibited the centrosome amplification (Figs. 3E and 3F), which suggests that the integrity of ROCK1 and 14-3-3σ complex is required for the centrosome amplification. These data suggest that the pathophysiological factors of type 2 diabetes activate AKT-ROS signalling which promotes the expression and binding of ROCK1 and 14-3-3σ as well as their translocation to centrosome to promote centrosome amplification.

High glucose, insulin and palmitic acid, alone or in combinations, did not affect the cell viability under the experimental conditions, with the exception that high glucose treatment increased cell viability (data not shown). In some cell line models, centrosome amplification is associated with cell cycle arrest (Yih et al., 2006). In our study, high glucose, insulin and palmitic acid, alone or in combination, did not disturb the cell cycle (data not shown), which suggests that cell cycle arrest is not a prerequisite for centrosome amplification under our experimental conditions. At the concentrations of 10, 20 and 30 μM, linoleic acid did not significantly affect the centrosome amplification (figure not shown), suggesting that unsaturated free fatty acids are unable to suppress the centrosome amplification.

In our study, 3.4% of the PBMC from the healthy subjects displayed centrosome amplification (Fig. 1B), which agrees with the finding by Dementyeva and do-workers that approximately 3% of the peripheral blood cells from healthy donors display centrosome amplification (Dementyeva et al., 2010).

It is known that obesity increases the risk for cancer (Mazzarella, 2005). There is also evidence that type 1 diabetes modestly increases the risk for cancer (Zendehdel et al., 2003). Obesity is associated with increased plasma levels of insulin and free fatty acids (Golay et al., 1986). Type 1diabetes is associated with hyperglycaemia and increased free fatty acid level. All these pathophysiological factors were able to cause centrosome amplification (Figs. 2C-2D). Thus, we speculate whether centrosome amplification could play a role the cancer development in obesity and type 1 diabetes. Moreover, our results (Figs. 3B and 3C) showed that the diabetic pathophysiological factors could upregulate ROCK1 and 14-3-3σ which are associated with neurodegeneration (Joo et al., 2015; Hu et al., 2016), upon which we further speculate whether ROCK1 and/or 14-3-3σ could be new clues for the development of Alzheimer's disease which shows an increased risk in type 2 diabetes (Zhang et al., 2016).

The study involved limited number of subjects. Further analysis with large cohorts is required to confirm the findings. In the study, we used PBMC to demonstrate the increase in centrosome amplification in type 2 diabetes. This raises a concern of whether centrosome amplification occurs in other tissues prone to diabetes-related cancer.

In conclusion, our results show that type 2 diabetes promotes cell centrosome amplification, and suggest that activation of AKT-ROS-dependent upregulations in expression, binding and centrosome translocation of ROCK1 and 14-3-3σ by the pathophysiological factors in type 2 diabetes is the underlying molecular mechanism.

## MATERIALS AND METHODS

### Chemicals, antibodies and cells

All chemicals were purchased from Sigma (St. Louis, MO, USA). Gama-tubulin antibody (No. ab27074; mouse antibody) was purchased from Abcam (Cambridge, UK). Rock1 antibody (No. 4035; rabbit antibody) was provided by cell signaling technology (Boston, MA, USA). 14-3-3σ antibody (No. PLA0201; rabbit antibody) was purchased from Sigma (St. Louis, MO, USA). Other antibodies were provided by Cell Signalling Technology (Boston, MA, USA). HCT116 colon cancer cells were kindly provided by Dr. B. Vogelstein of the Johns Hopkins University School of Medicine, who produced the cell line. Normal human breast epithelial cells (PCS-600-010) and culture medium were purchased from the American Type Culture Collection (ATCC, Manassas, VA, USA). The culture medium and reagents for the colon cancer cells were purchased from Gibco (Beijing, China). The palmitic acid stock was conjugated to fatty acid-free bovine albumin in a 3:1 molar ratio at 37 °C for 1 hour before use. Anti-gamma tubulin antibody was used to detect centrosome by innumofluorescent staining. ROCK1 siRNA treatment largely reduced the staining of ROCK1. 14-3-3 was seen in the centrosome only after experimental treatment. These observations showed that non-specific staining by the antibodies would not affect the experimental observations.

### Clinical study

Institutional approval and written informed consent were obtained from the medical ethics committee and all participating subjects, respectively. All the volunteers were consecutively recruited during 2014 and 2016 at the Shanxi Hospital of Integrative Western and Chinese Medicine and Shanxi Medical University without any selection bias. Type 2 diabetes was diagnosed according to the 1999 WHO criteria. All healthy subjects were free hypertension. Each volunteer donated 5-ml blood sample. All the clinical data were collected by clinical doctors responsible for the clinical study.

### Cell culture

HCT116 cells were maintained in DMEM (low glucose, 5mM) supplemented with 50 U/ml penicillin, 50 g/ml streptomycin and 10% (v/v) foetal calf serum. Human primary breast epithelial cells were cultured in basal medium (PCS-600-030) with a growth kit (PCS-600-040, ATCC, Manassas, USA) according to ATCC instructions. Epithelial cells from the second passage in our lab were used in the study. In cell model studies, cells treated for 12 hours were used for ROS quantification. Cells treated for 24 hours were harvested for co-immunoprecipitation assay. Those treated for 30 hours were used for transcriptomic analysis. Cells treated for 48 hours were used for quantification of centrosome number and protein distribution in centrosomes. We performed time course assays and the time points were chosen, since these time points produced the highest level of differences for the measurements between the control and the treated samples.

### Quantification of ROS

Changes in the intracellular ROS levels were determined using the 2,7’-dichlorofluorescein (DCFH-DA, Beyotime, Shanghai, China) probe on a spectrofluorometer (SpectraMax M5, Molecular Devices, Silicon Valley USA). Cells grown in a 24-well plate were treated, washed twice with PBS, and incubated with DCFH-DA for 20 min at 37 oC. Then, the cells were harvested, washed once in PBS, and transferred to a 96-well plate. Optical density values were obtained at an excitation wavelength of 488 nm and an emission wavelength of 525 nm. Alternatively, the ROS levels were quantified using a flow cytometer (FACSCalibur, BD, New Jersey, USA) according to the manufacturer’s instructions.

### Confocal microscopy

A cover slip was placed in a well of a 6-well plate. Cells were plated at a density of 50,000 cells per well. Cells grown on the cover slips were fixed in cold methanol and acetone (1:1; v:v) for 6 min at -20°C, followed by three washes with PBS (10 min each time). Then, the cells were incubated with 0.1% Triton X-100 for 15 minutes and 3% BSA for 1 hour. The cells were incubated with a primary antibody in 3% BSA in PBS overnight at 4 °C, washed twice with PBS, and incubated with a FITC-conjugated secondary antibody in 3% BSA in 1×PBS for 1 hour at room temperature in the dark. Finally, the cells were mounted with mounting medium. Confocal microscopy was performed using the Zeiss LSM880 microscope (Oberkochen, Germany) with a 1.4 NA oil-immersion lens, and image processing was performed with Zen software (Oberkochen, Germany). One hundred cells were counted for their centrosome numbers for the percentage of centrosome amplification.

To separate peripheral blood mononuclear cells (PBMC), we loaded a 5-ml blood sample on the top of separation medium (LTS1077; TBD, Tianjin, China) in a centrifuge tube and centrifuged at 250 g for 20 min. The cells on the top were collected, washed twice in PBS, smeared onto a cover slip, and air dried for 24 hours at room temperature.

### Transcriptomic profiling and bioinformatic annotation

The cells were harvested after treatment and total RNA was extracted. Two cDNA libraries (control or treatment with high glucose, insulin, and palmitic acid) were constructed using the Illumina TruSeq RNA Sample Preparation Kit (Illumina, USA) according to the manufacturer’s instructions. After several steps of purification, adapter addition, and cDNA length selection, the two libraries were sequenced independently using an Illumina HiSeq^(tm)^500 platform (Shanghai Personal Biotechnology Co., Shanghai, China). Pathway assignments were generated using GO (geneontology.org) and KEGG databases (www.kegg.jp).

### Western blot analysis

The cells were lysed in RIPA buffer. Proteins were separated by polyacrylamide gel electrophoresis and transferred onto PVDF membrane. After blocking for 1 hour at room temperature with TBST containing 0.05% (v/v), Tween-20 and 5% (w/v) non-fat milk, the membranes were incubated with primary antibodies overnight at 4 °C, followed by washes with TBST containing 0.05% Tween-20. The membranes were then incubated with a horseradish peroxidase-conjugated secondary antibody for 1 hour at room temperature. ECL reagents (Thermo Biosciences, Massachusetts, USA) were used to visualize the protein bands which were captured on X-ray film.

### Co-immunoprecipitation

Cells were harvested under non-denaturing conditions, washed by ice-cold PBS for 3 times, lysed in 0.5 ml ice-cold cell lysis buffer and centrifuged. Supernatant was collected to a new tube and incubated with 20 ul Protein G Plus /Protein A agarose (Miliproe, IP05 USA) with gentle shaking for 2 hours at 4 oC. Protein G Plus /Protein A agarose was then removed by centrifuge for 10 min at 4 oC, and supernatant was incubated with primary antibody overnight at 4 oC with gentle shaking. After that, 30ul Protein G Plus /Protein A agarose were added and incubated under gentle shaking for 4 hours at 4 oC. Finally Protein G Plus /Protein A agarose was collected by centrifuge.

### Knockdown of protein level

The pre-designed siRNA oligonucleotides (Songon Technology, Shanghai, China) for Akt1, ROCK1 and 14-3-3σ were:

GAGUUUGAGUACCUGAAGCUGUU (sense) and

AACAGCUUCAGGUACUCAAACUC (antisense);

UGAUCUUGUAGCUCCCGCAUGUGUC (sense) and ACUAGAACAUCGAGGGCGUACACAG (antisense); and ACCTGCTCTCAGTAGCCTA (sense) and TAGGCTACTGAGAGCAGGT (antisense), respectively. HCT116 cells (5×104 cells per well) were seeded in 6-well plates and cultured for 24 hours, and then were transfected with 200 pM siRNA oligonucleotides using Lipofectamine 2000 transfection reagent (Invitrogen, California, USA), according to the manufacturer’s instructions. The total AKT protein level was evaluated by Western blot analysis 24 hours after transfection.

### Statistical analysis

All the experiments including the transcriptomic profiling were performed in triplicate. The data are expressed as the mean ± SD. Student’s t-test was performed to compare the difference between two groups. Multi-group comparisons were performed using one-way ANOVA analysis. Linear regression analysis was performed for correlations. The statistical analysis software package SPSS21 was employed for the statistical comparisons. A p value < 0.05 was considered statistically significant.

## Acknowledgement

The authors thank Dr. ZY Li for her help in establishing our cell culture facility.

## Author contribution

P Wang and YC Lu performed most of the experiments. J Wang and YF LI contributed the clinical studies. L Wang provided the technologies. H Yu was involved in grant applications and the study design. A Kong and J Chan contributed the ROS data set and were involved in designing the study. SC Lee was the principal investigator, who was in charge the whole project, from grant application and study design to manuscript preparation.

---

**Table S1.**
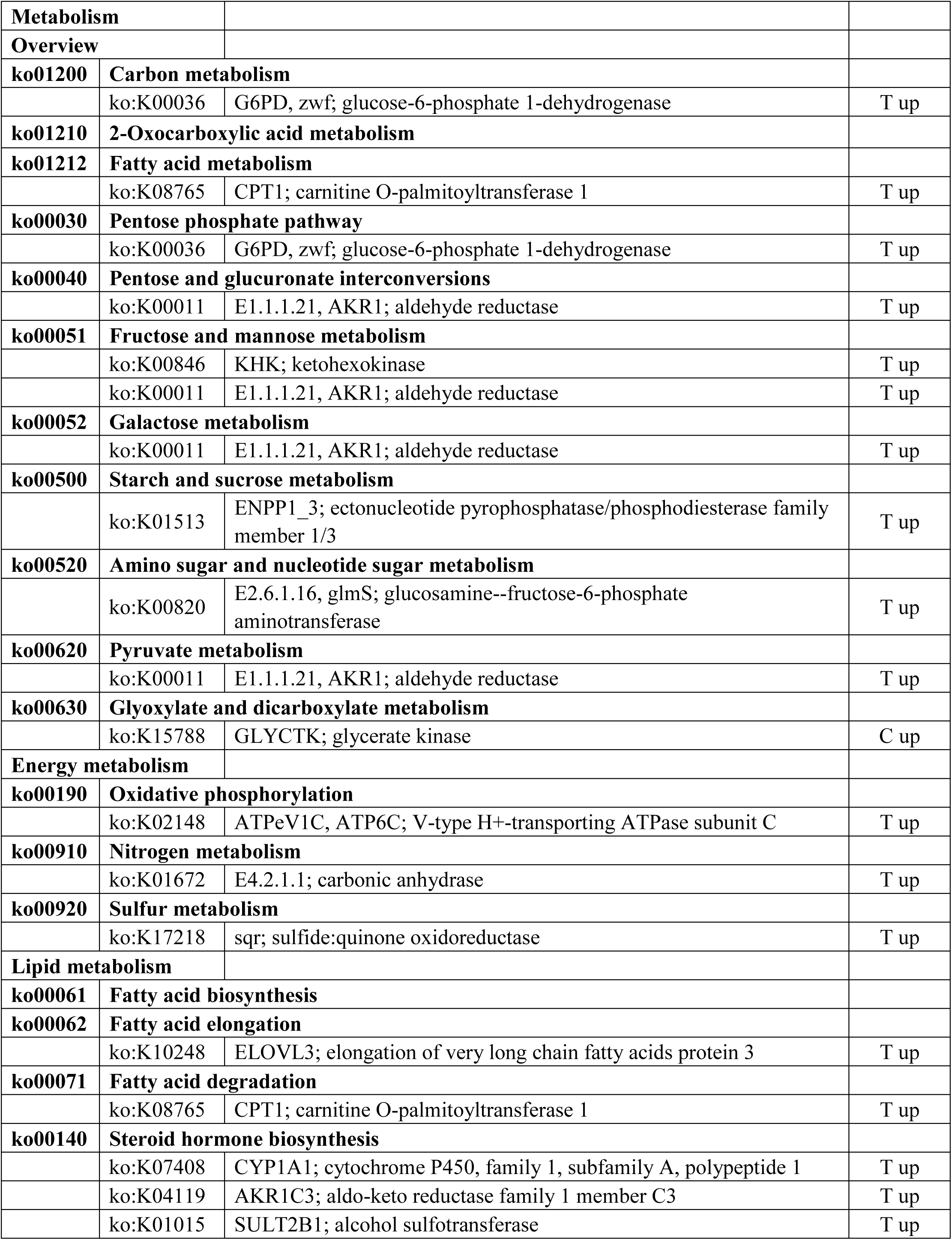
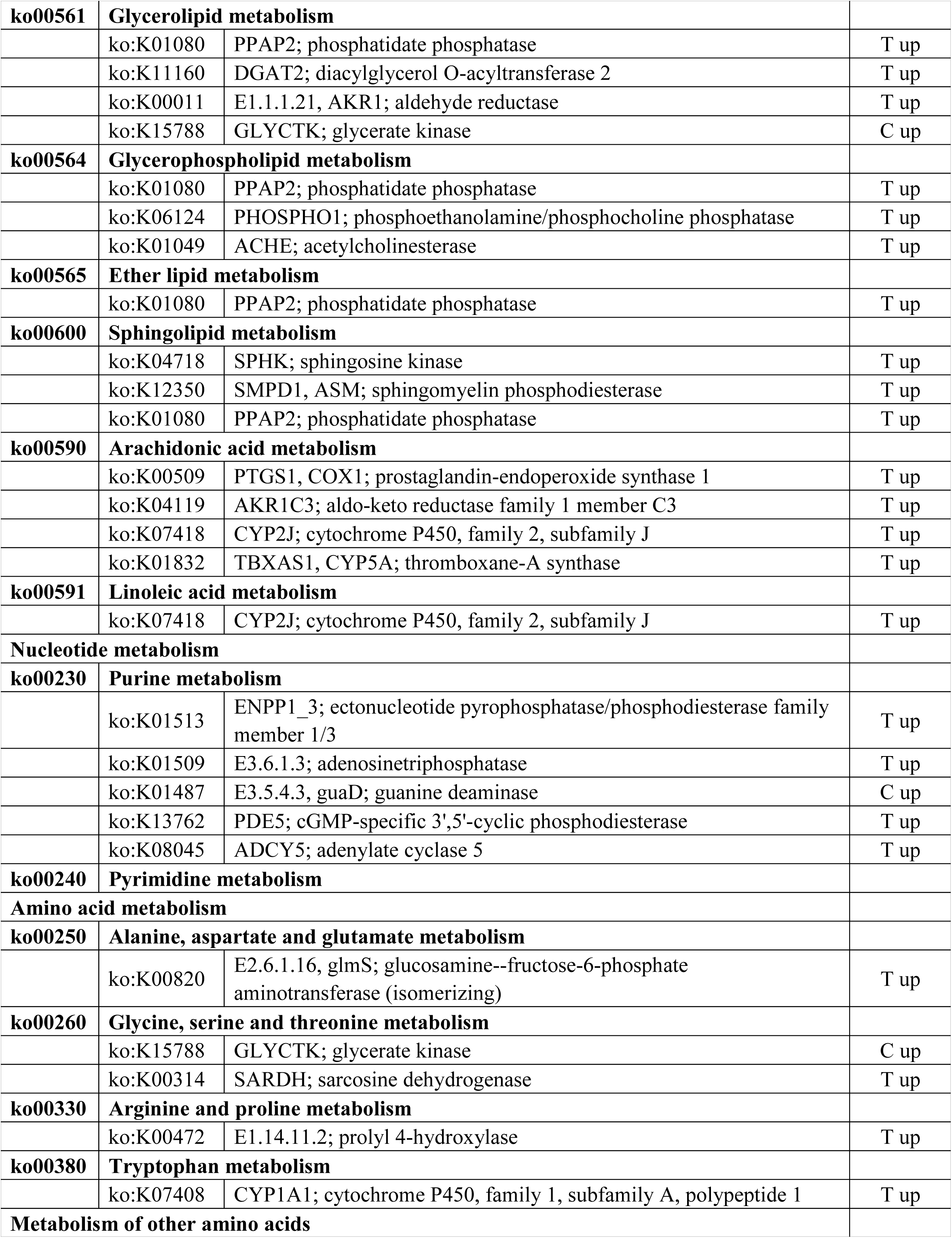
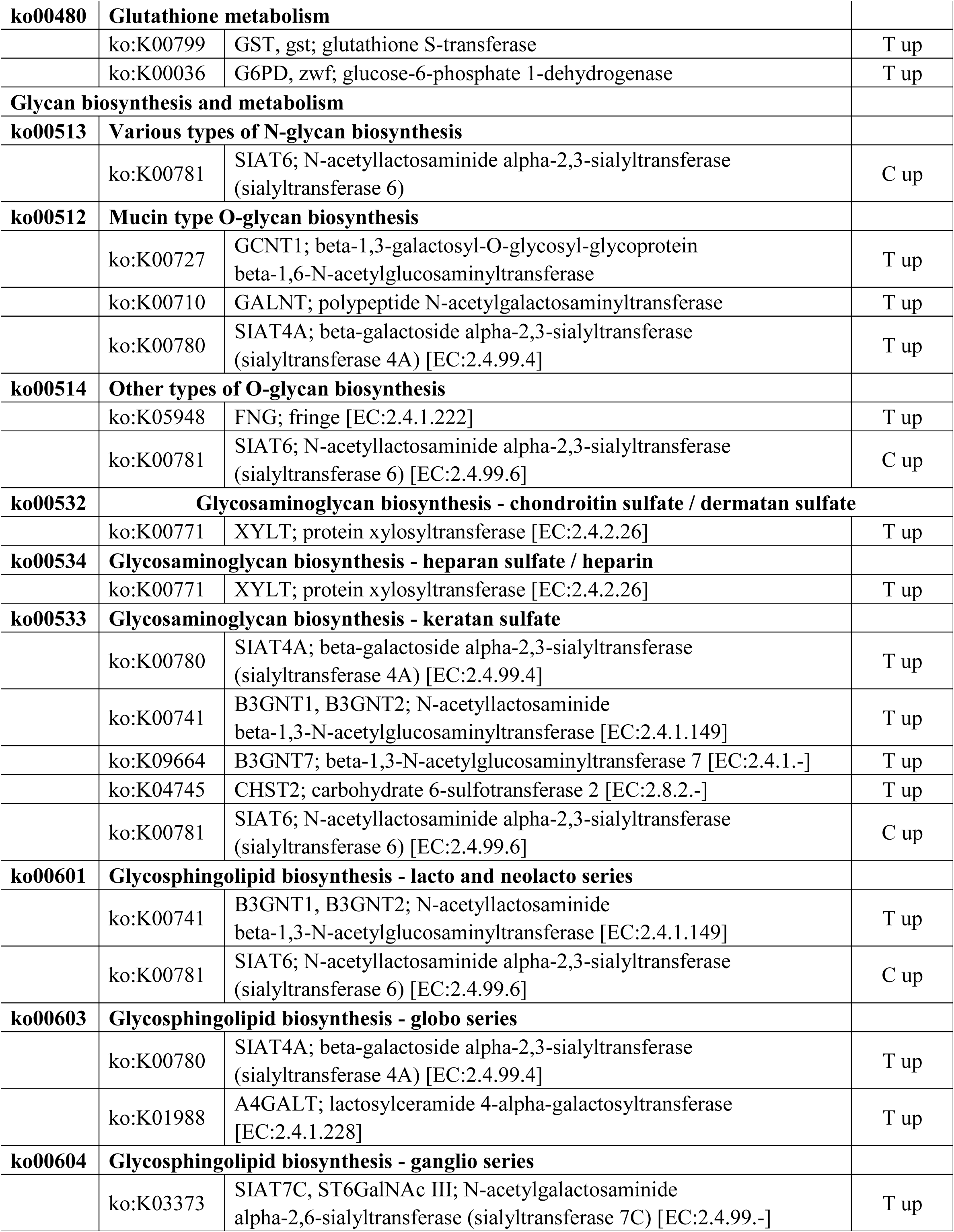
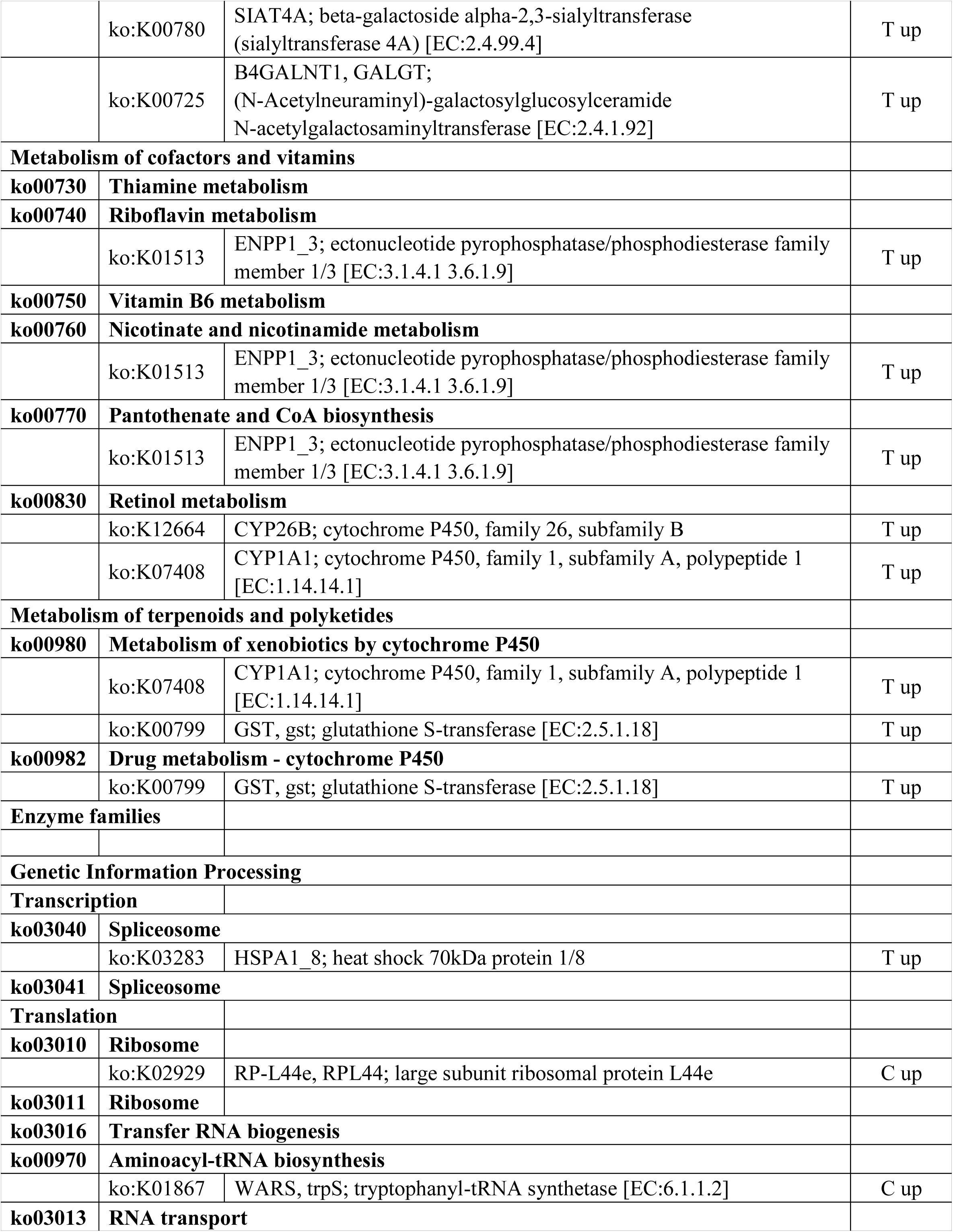
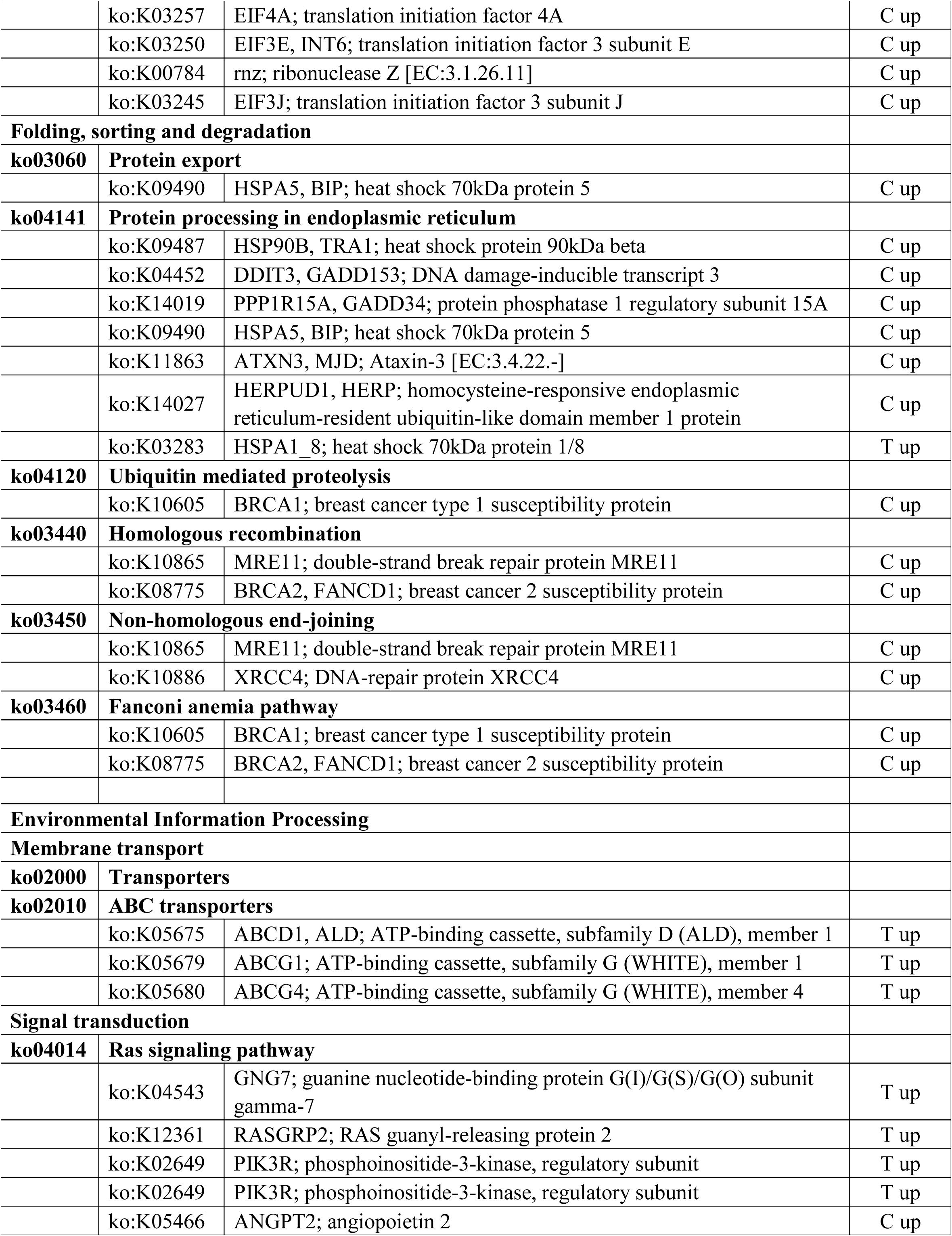
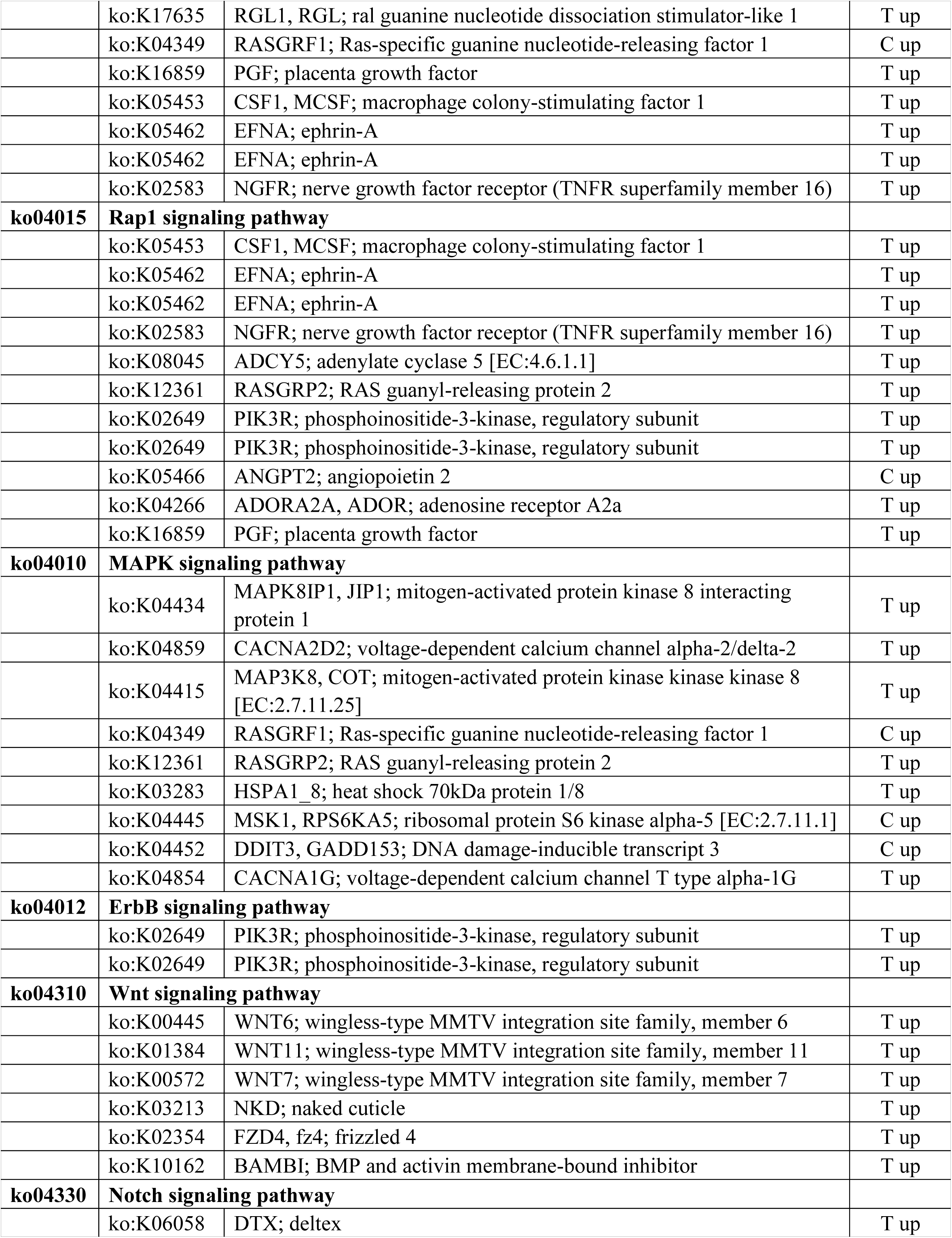
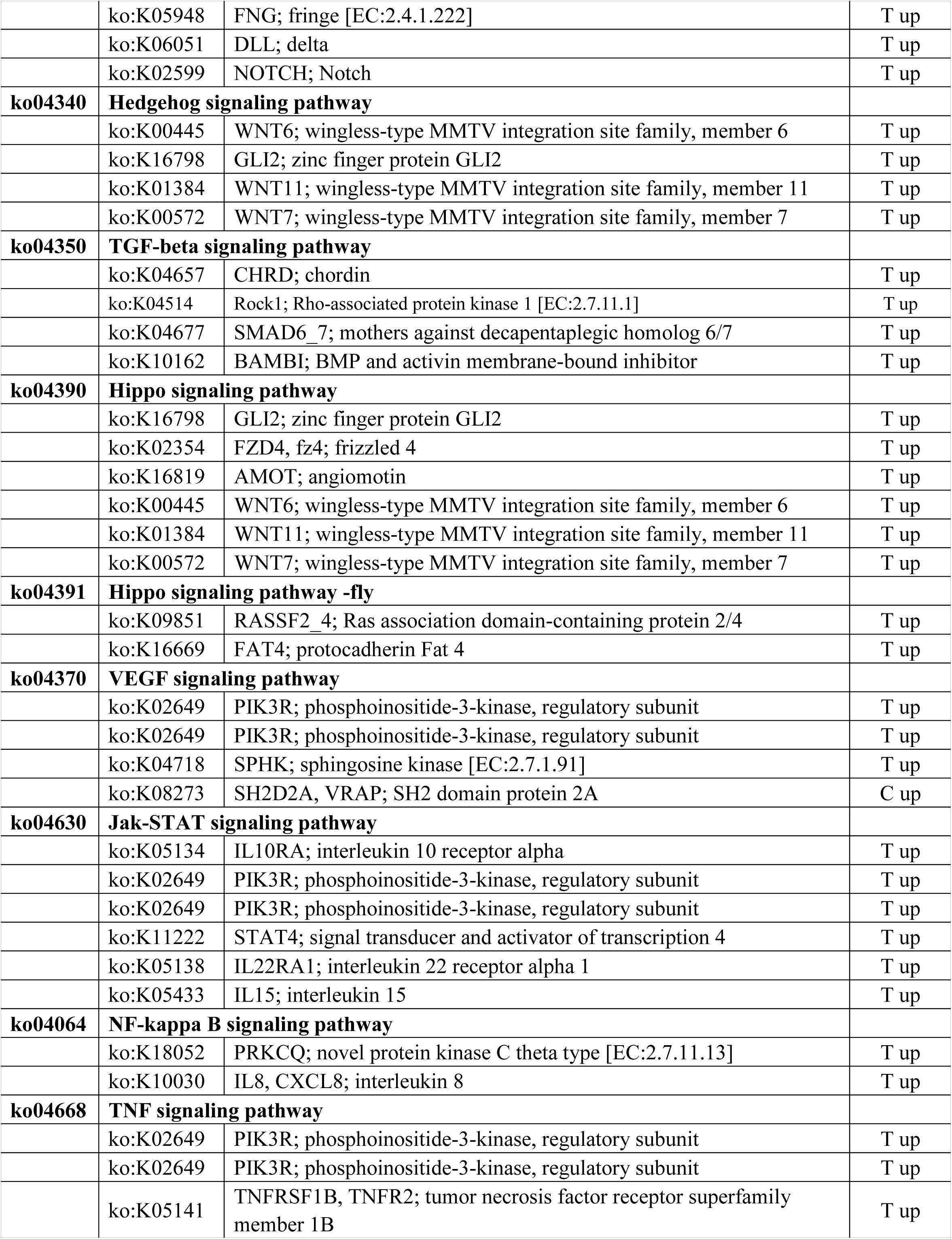
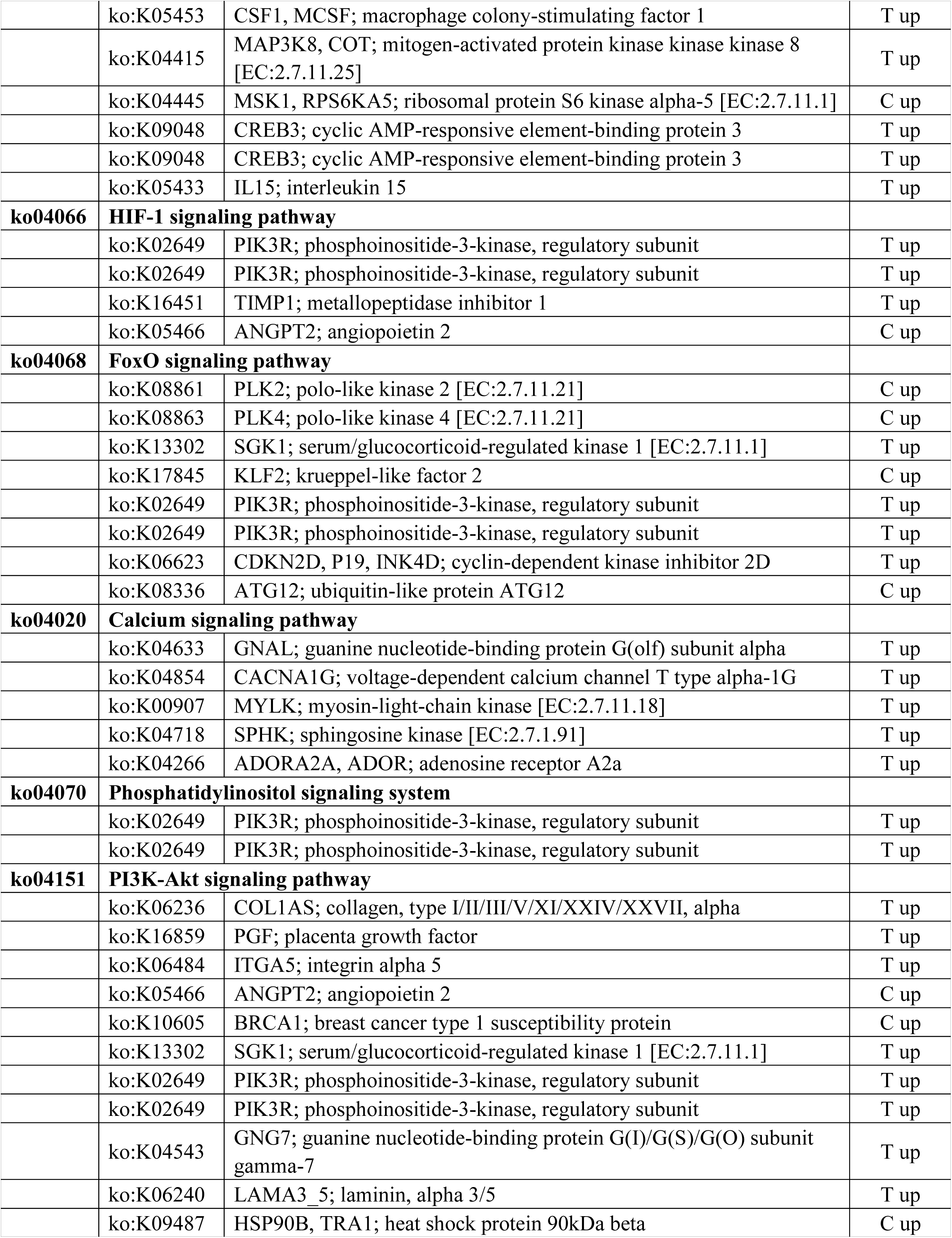
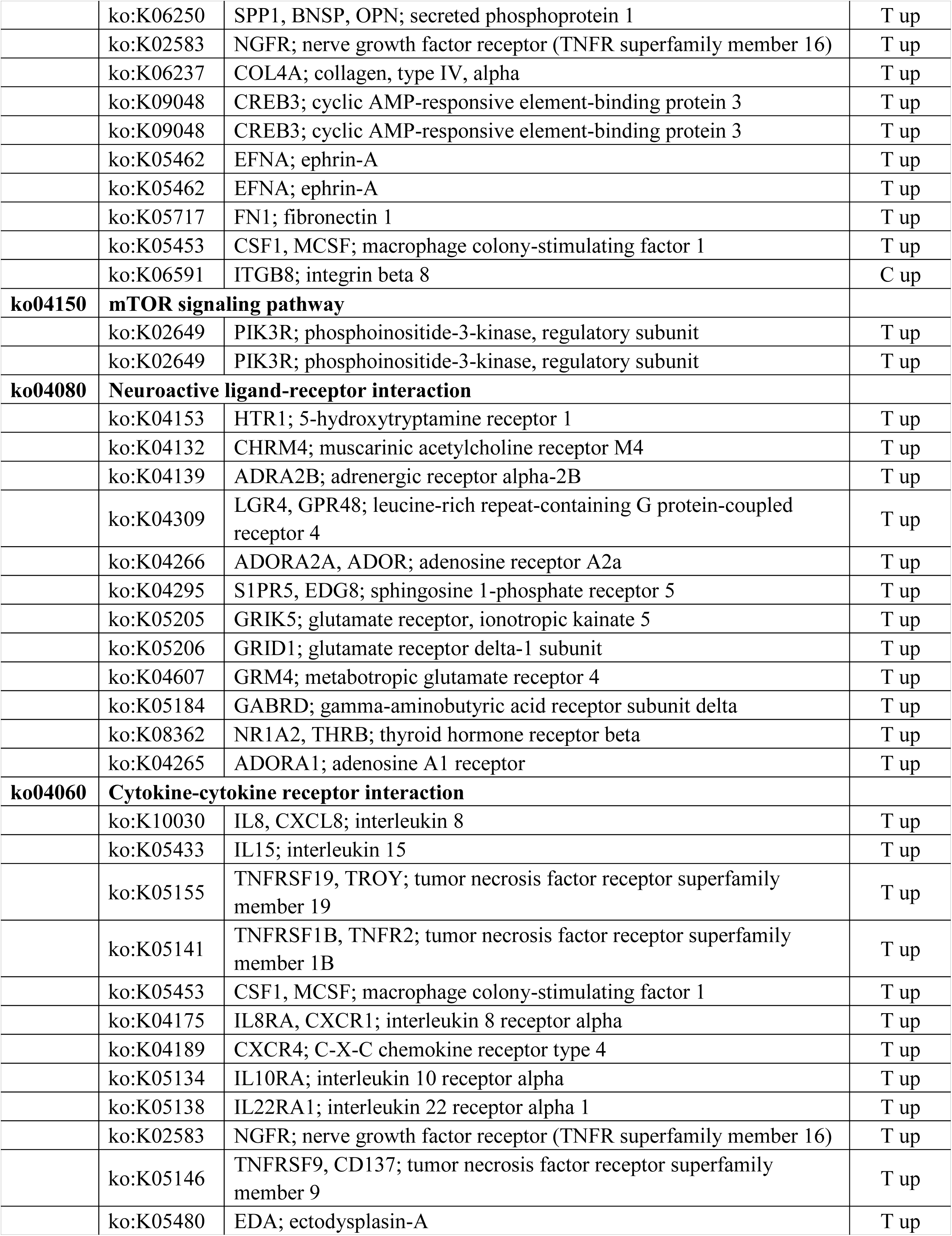
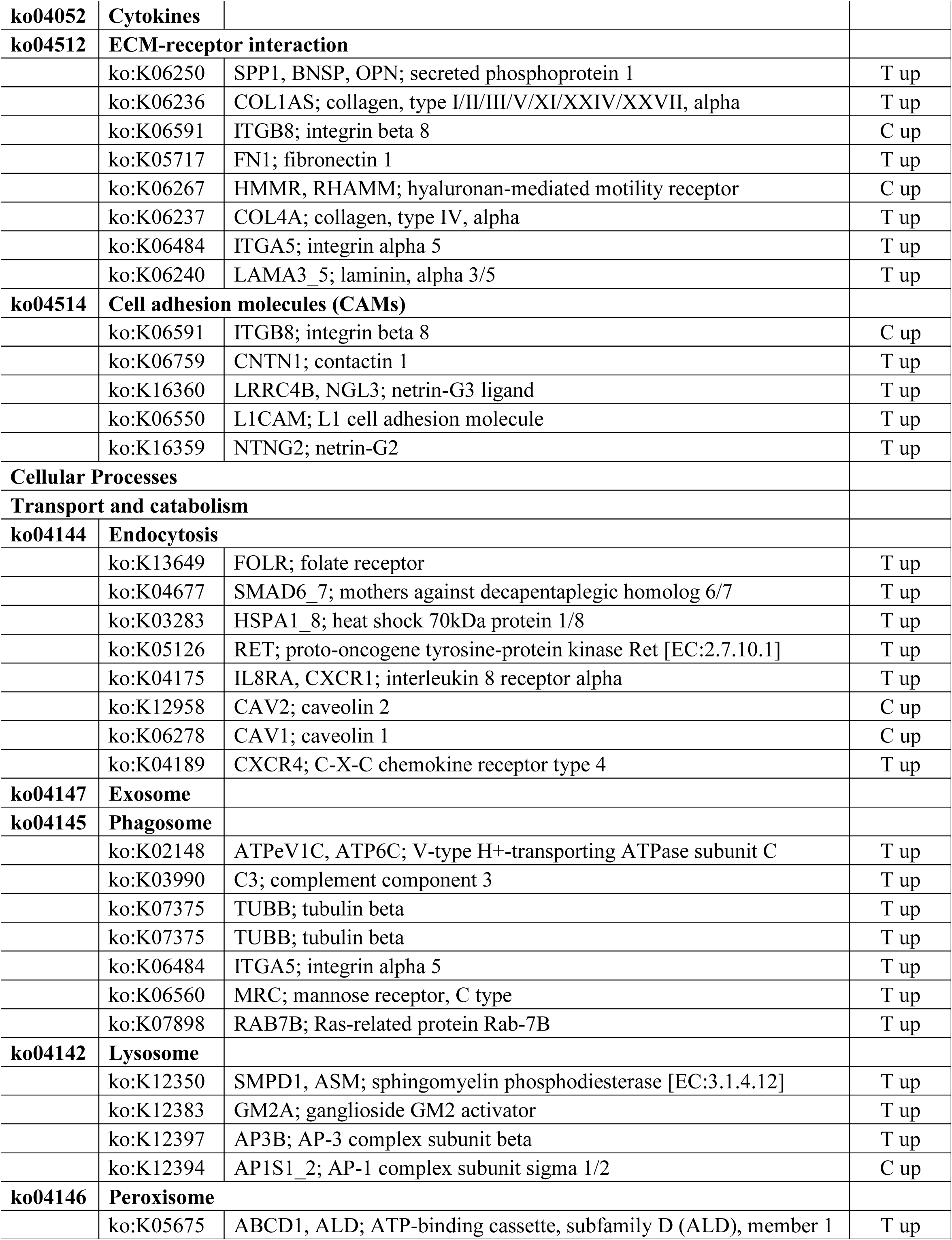
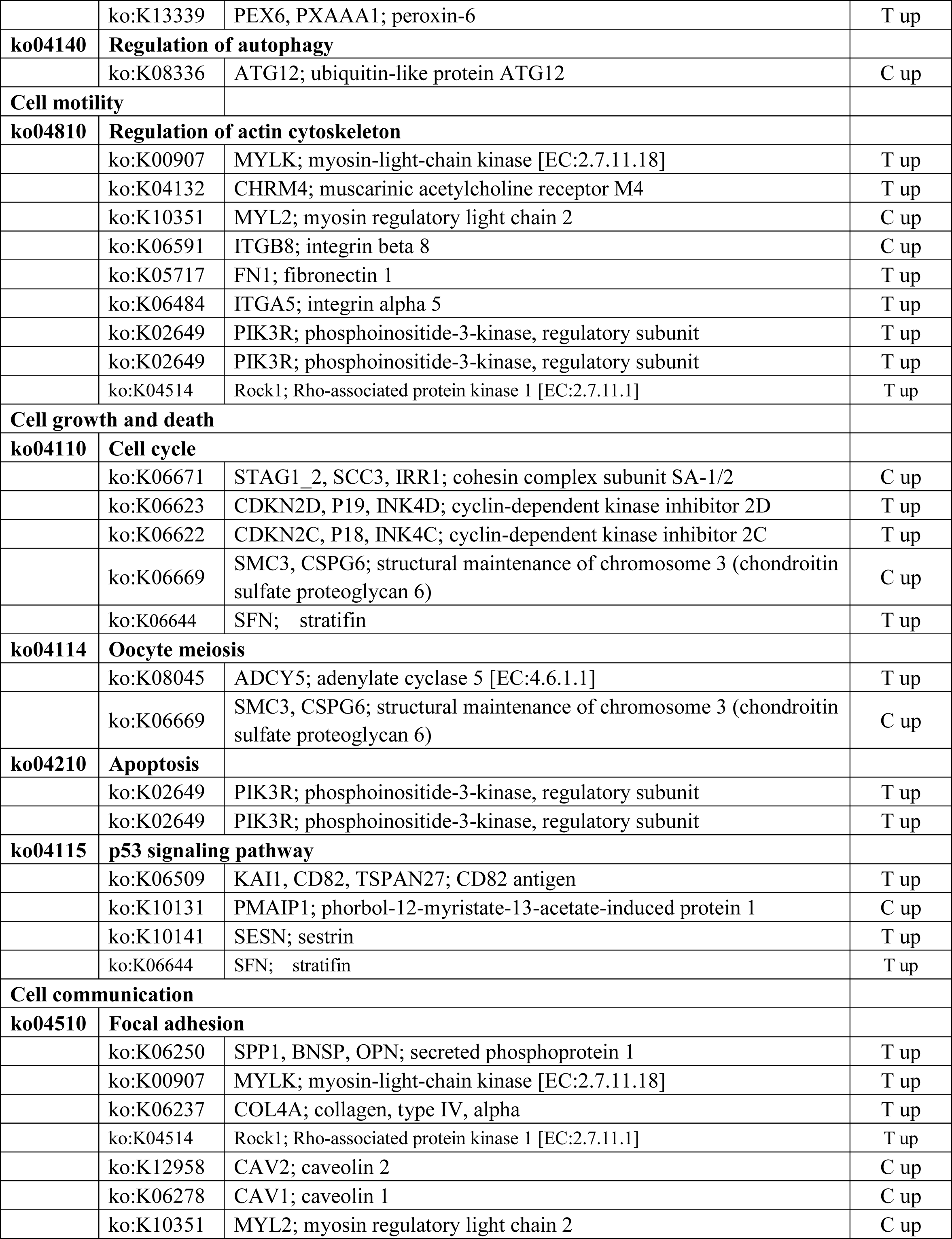
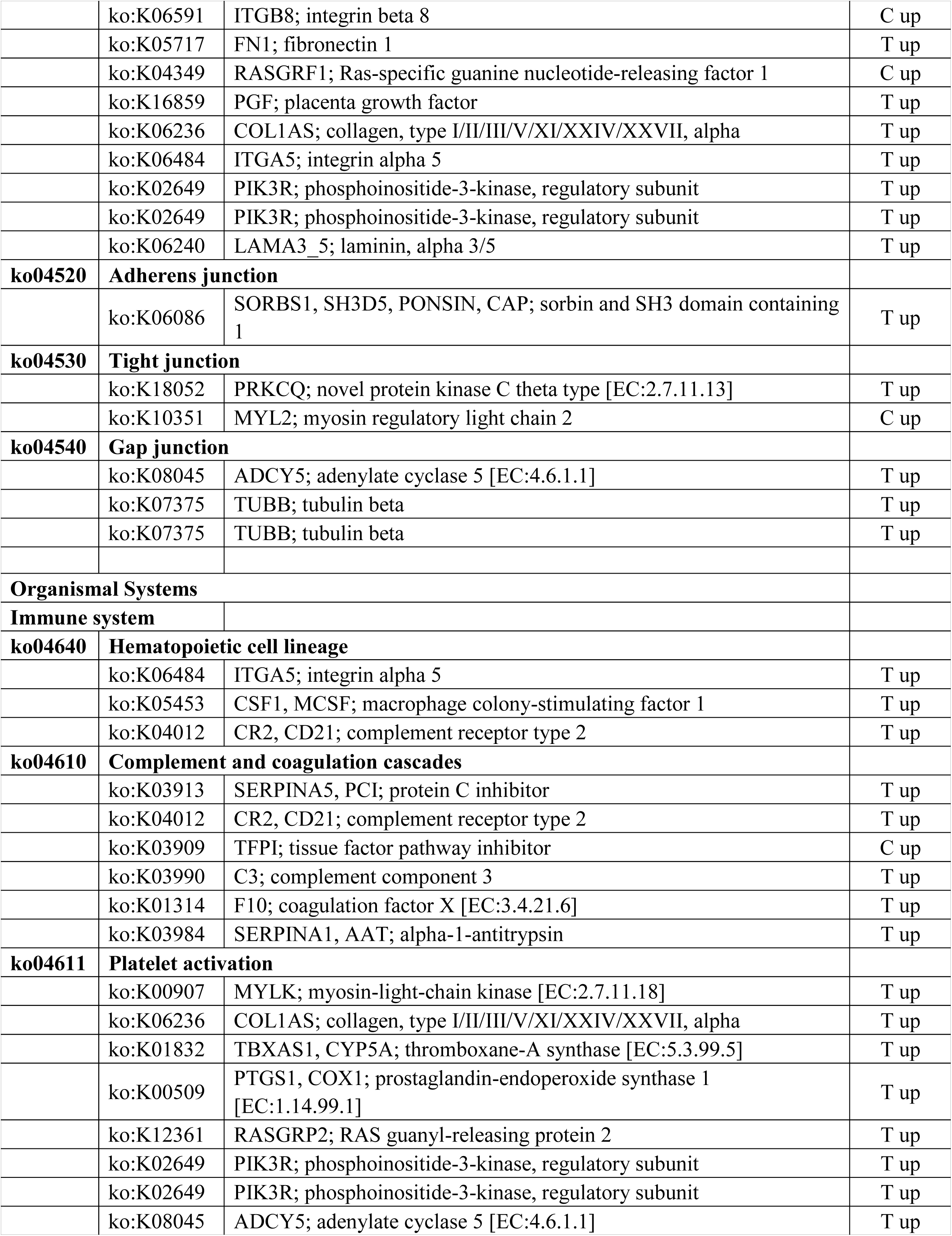
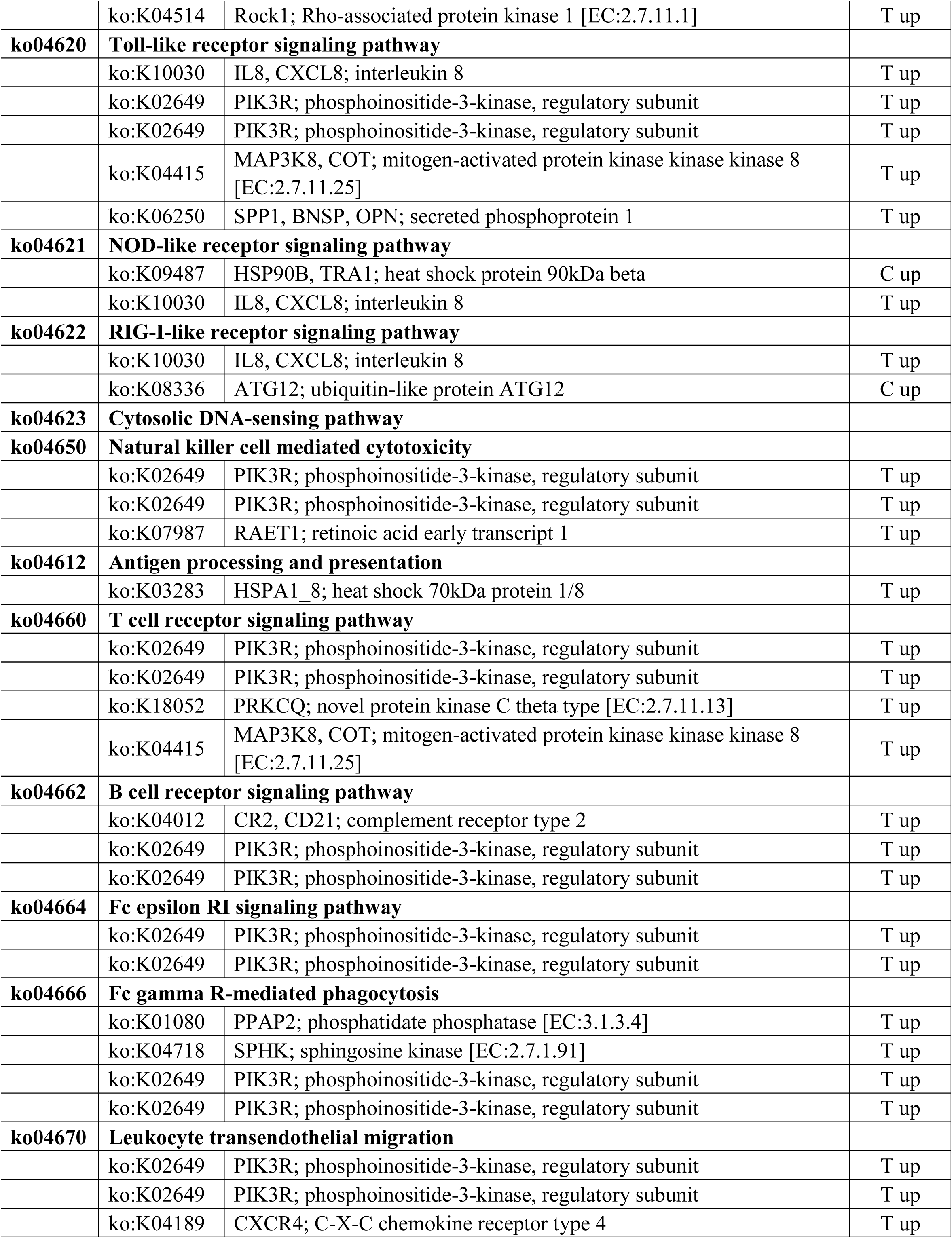
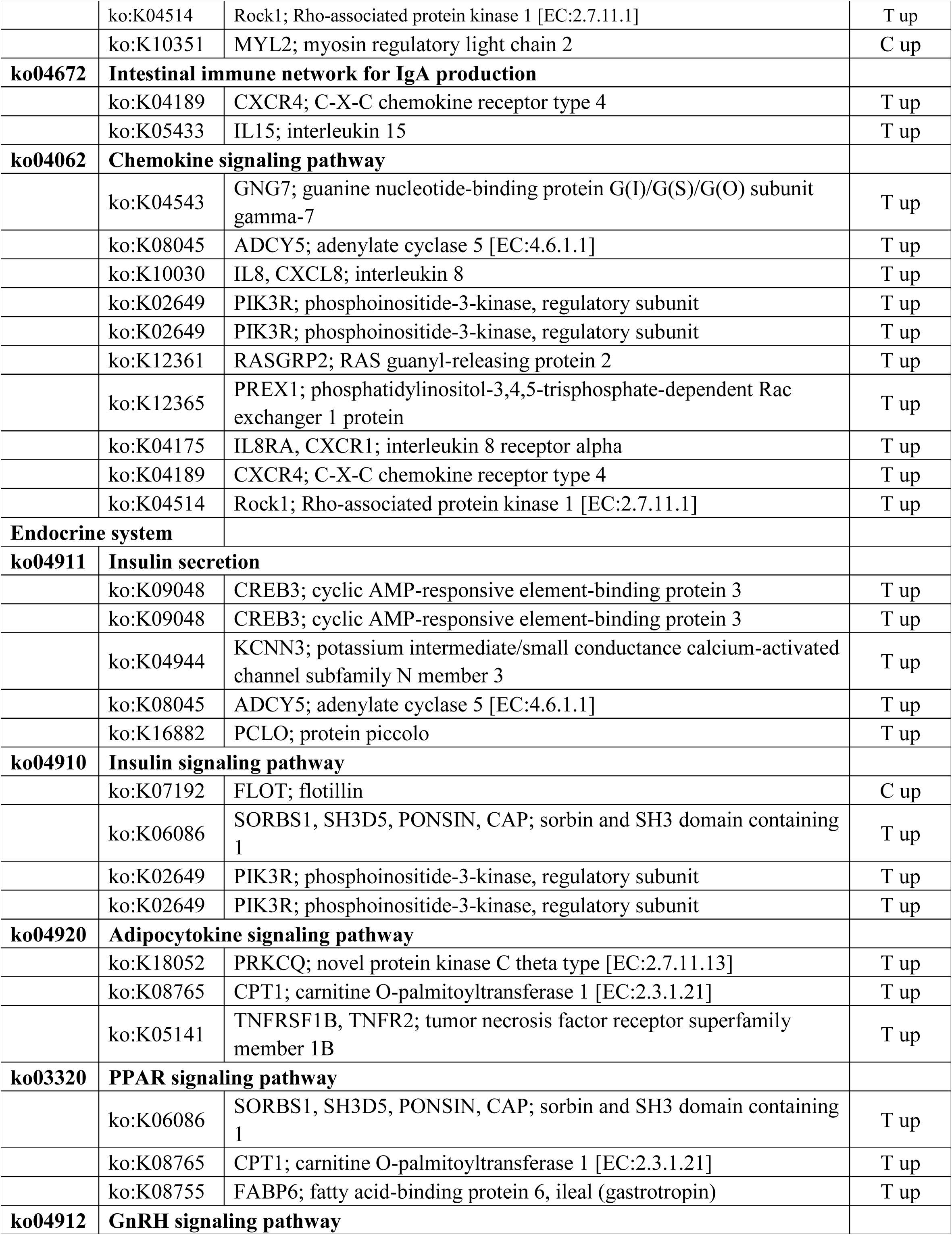
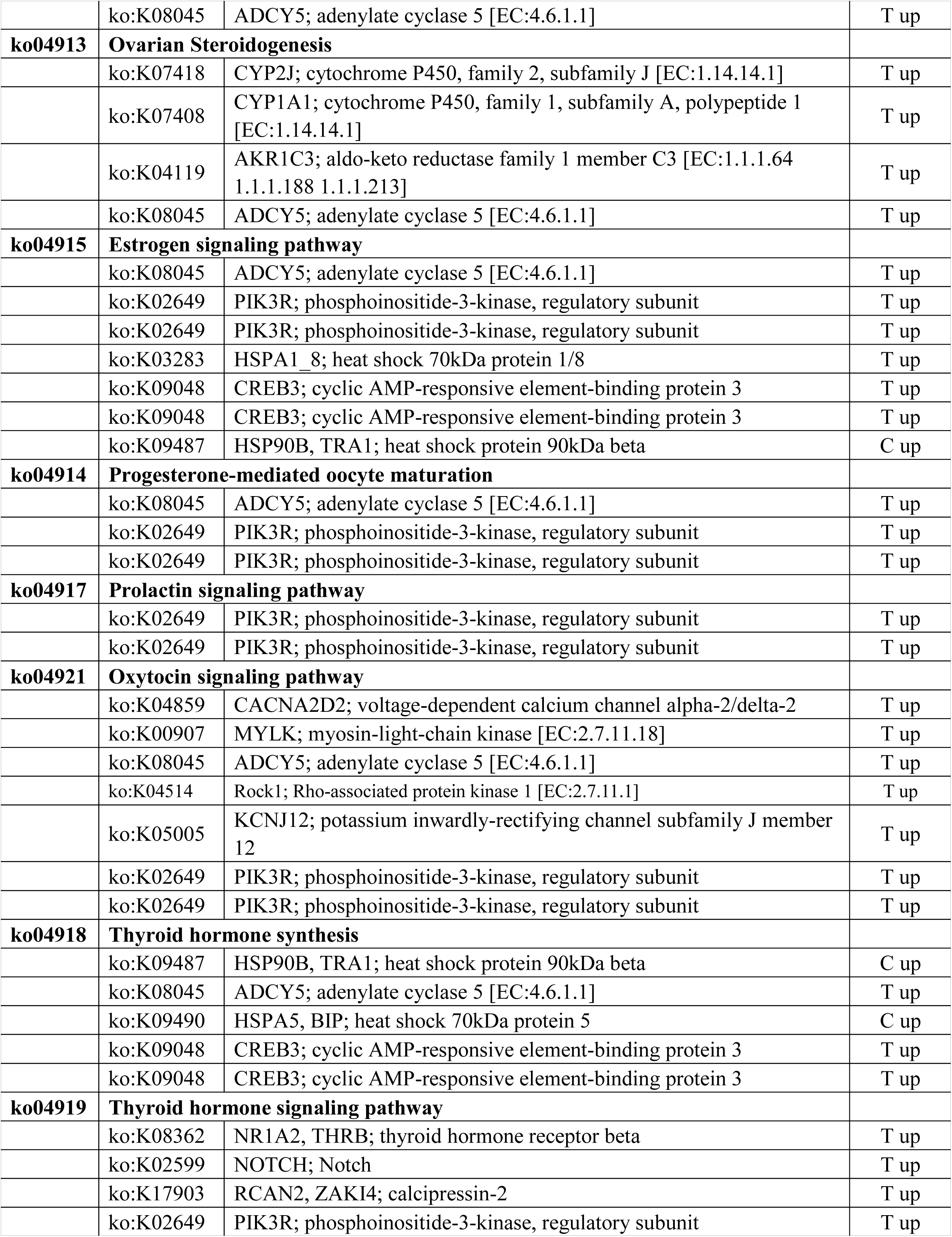
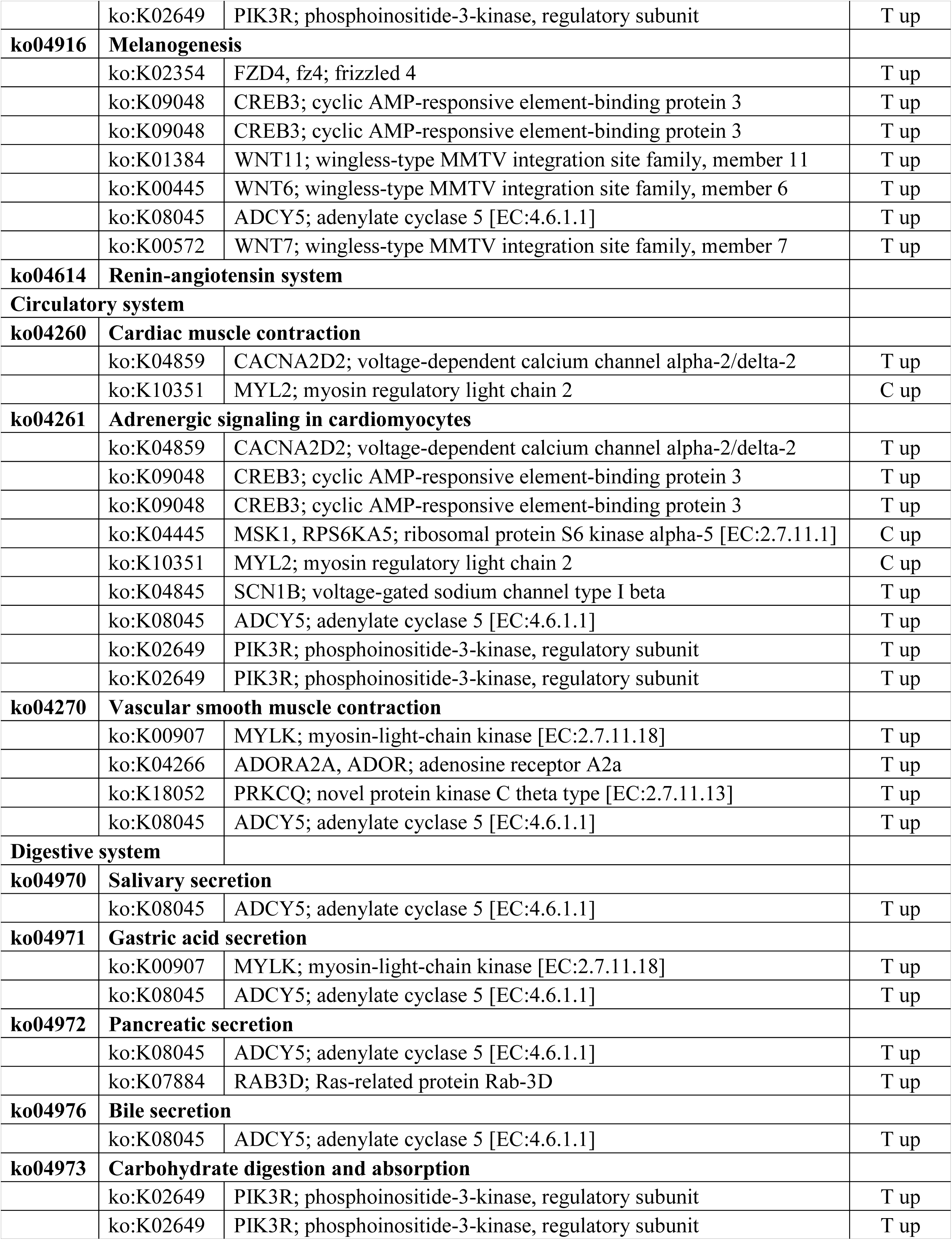
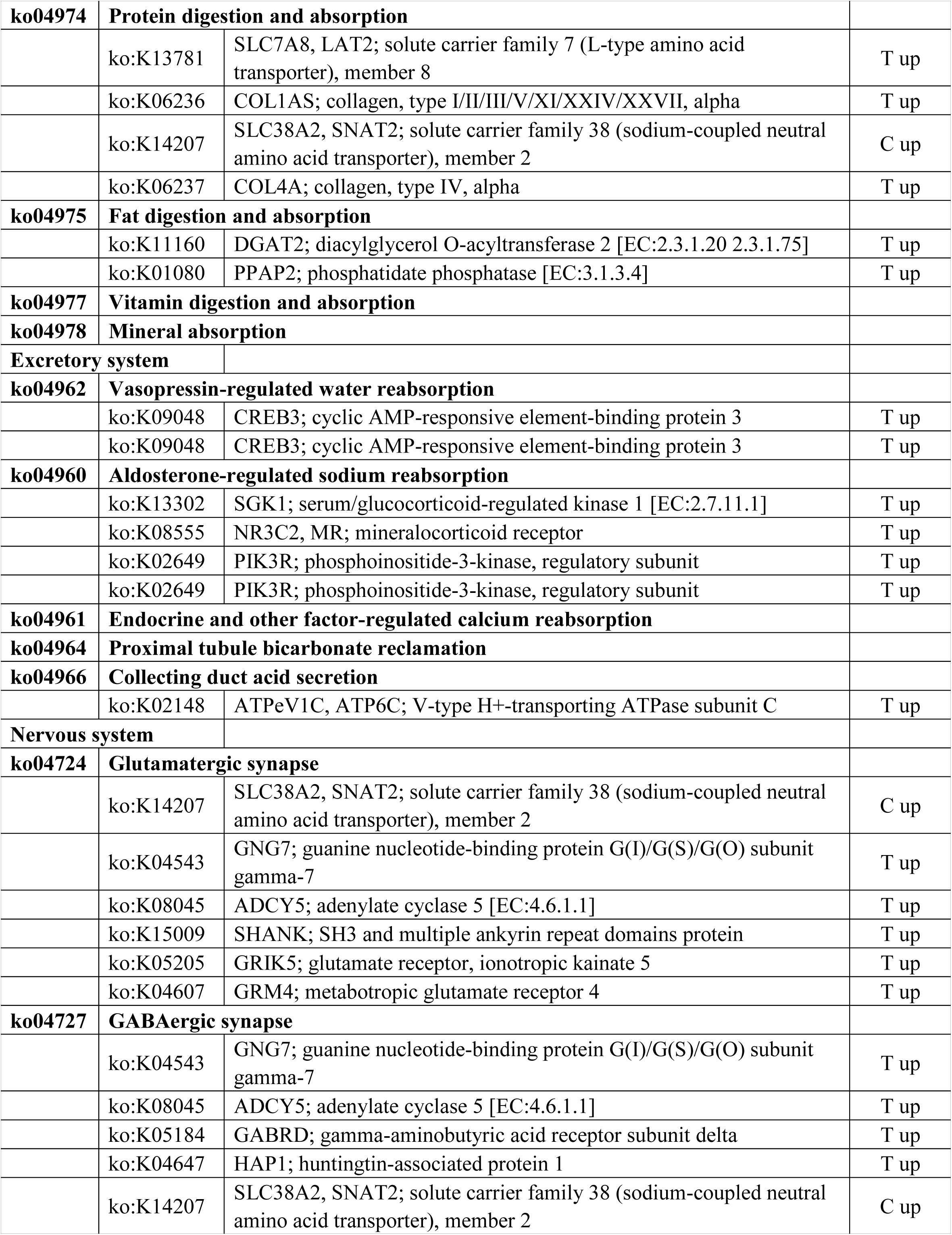
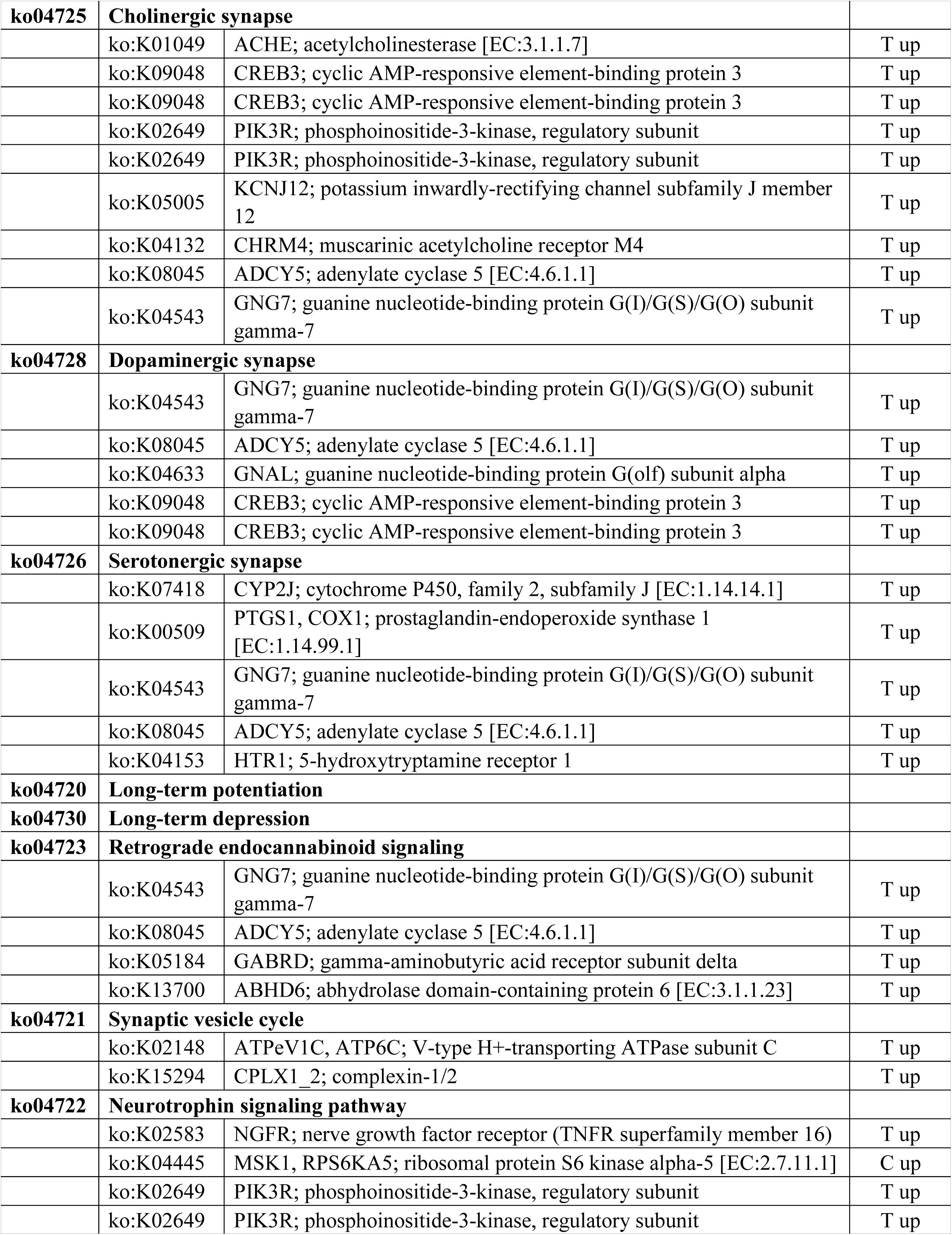
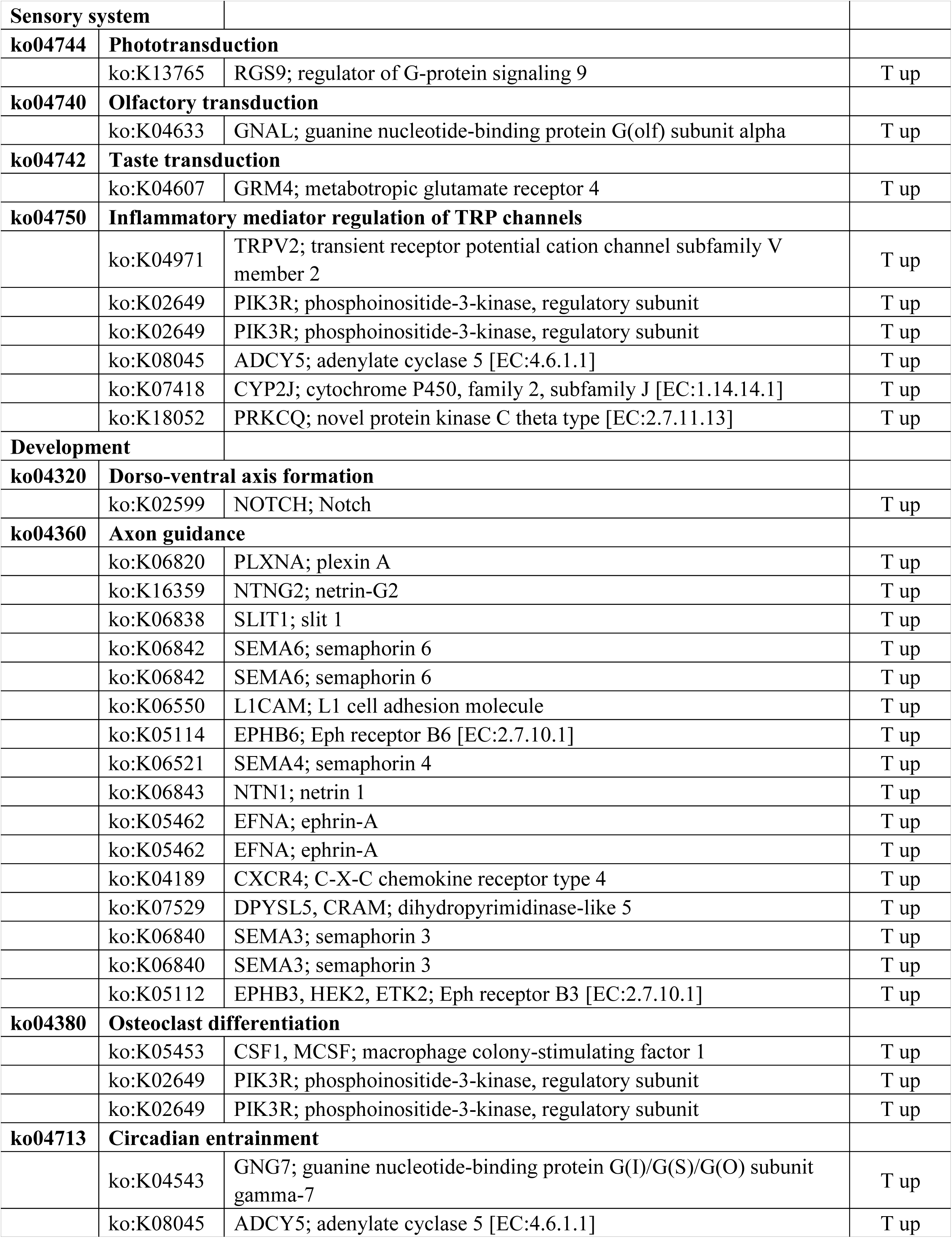
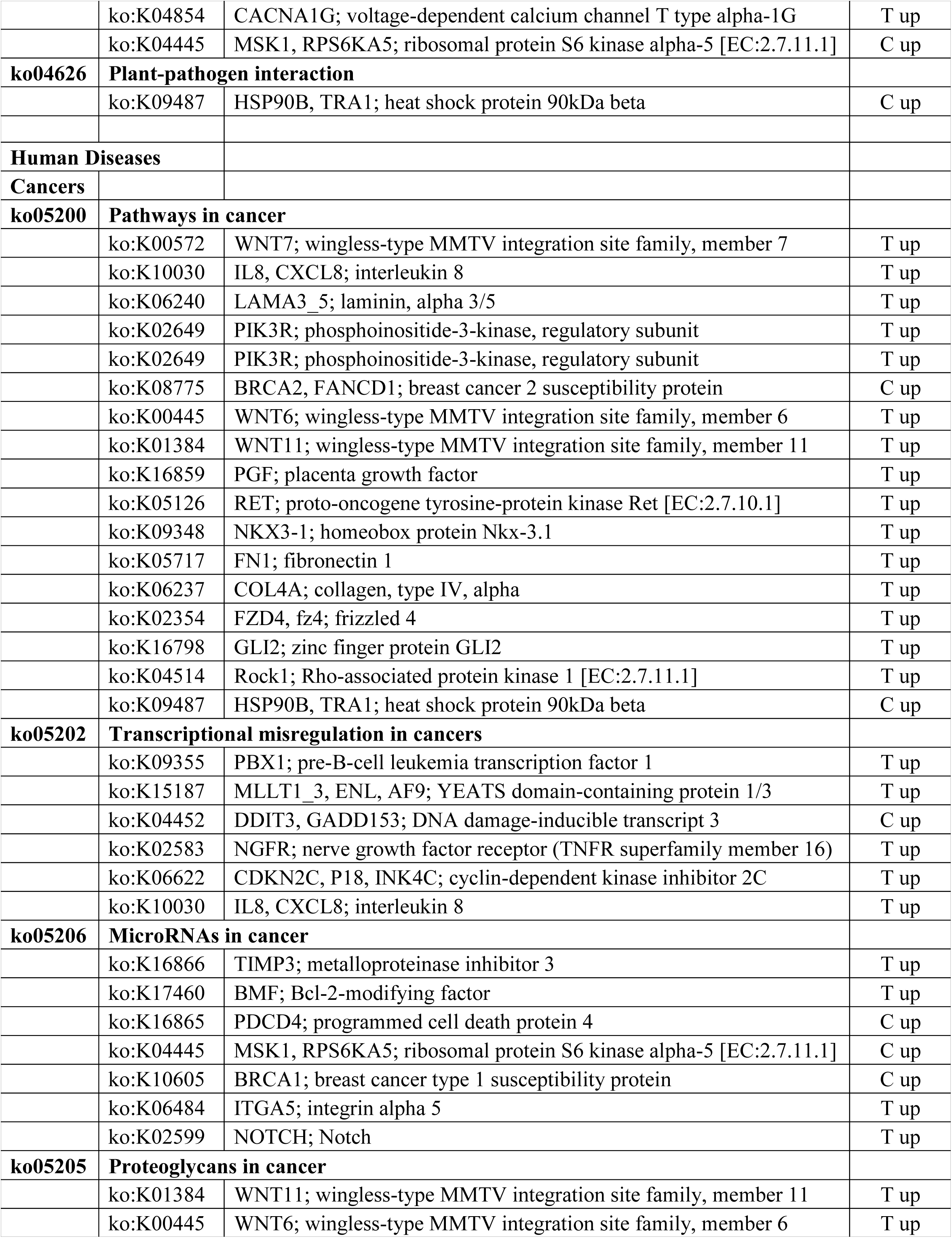
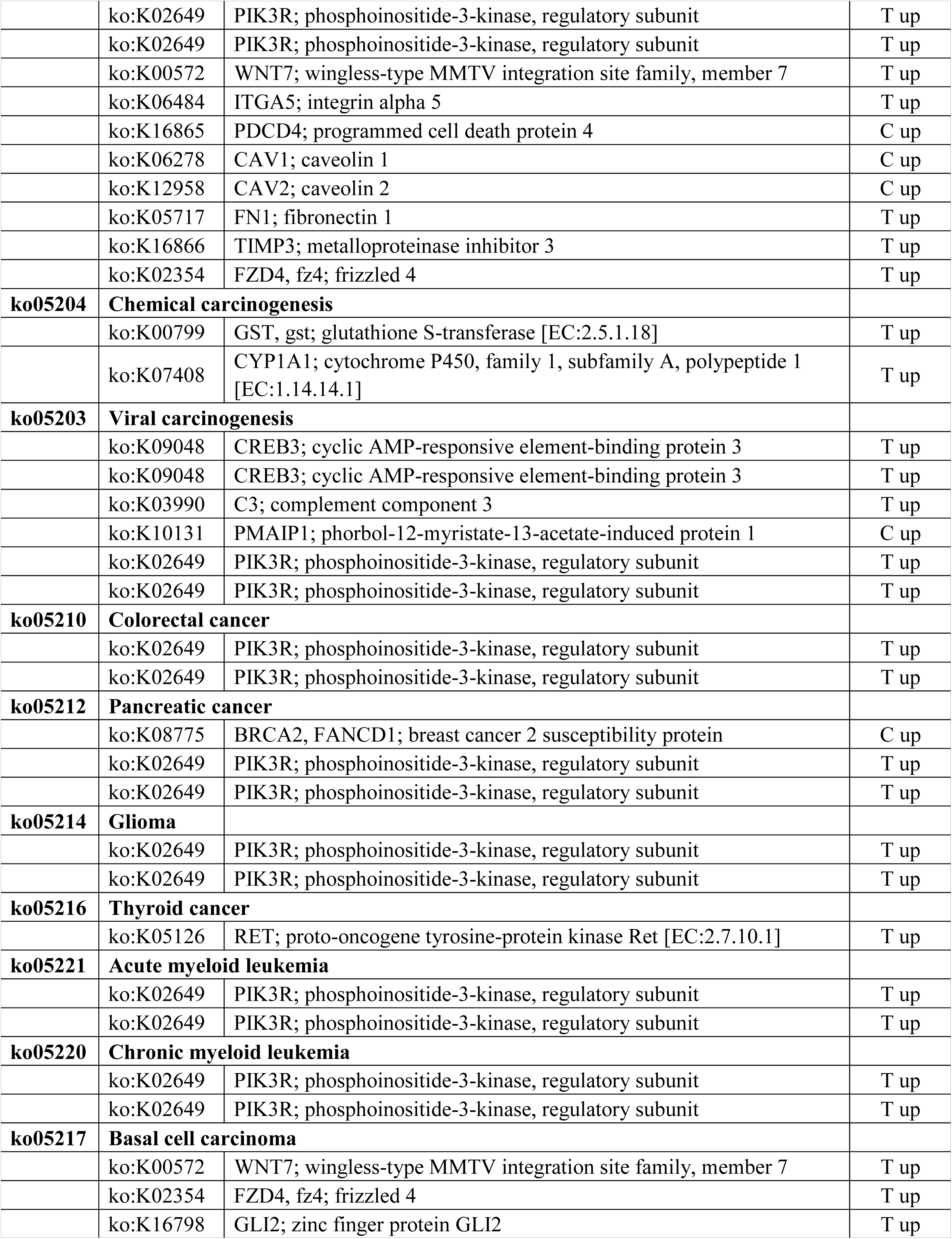
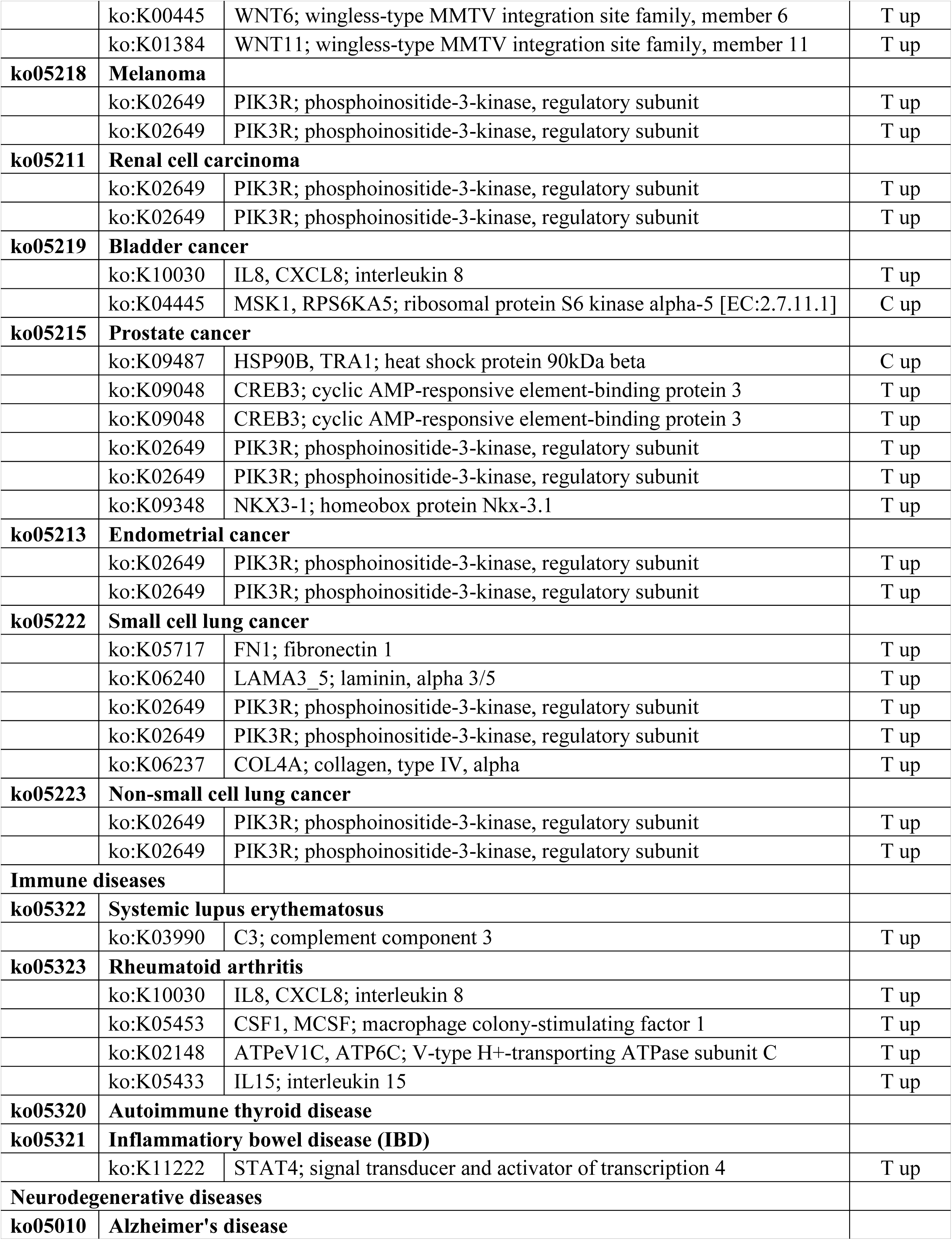
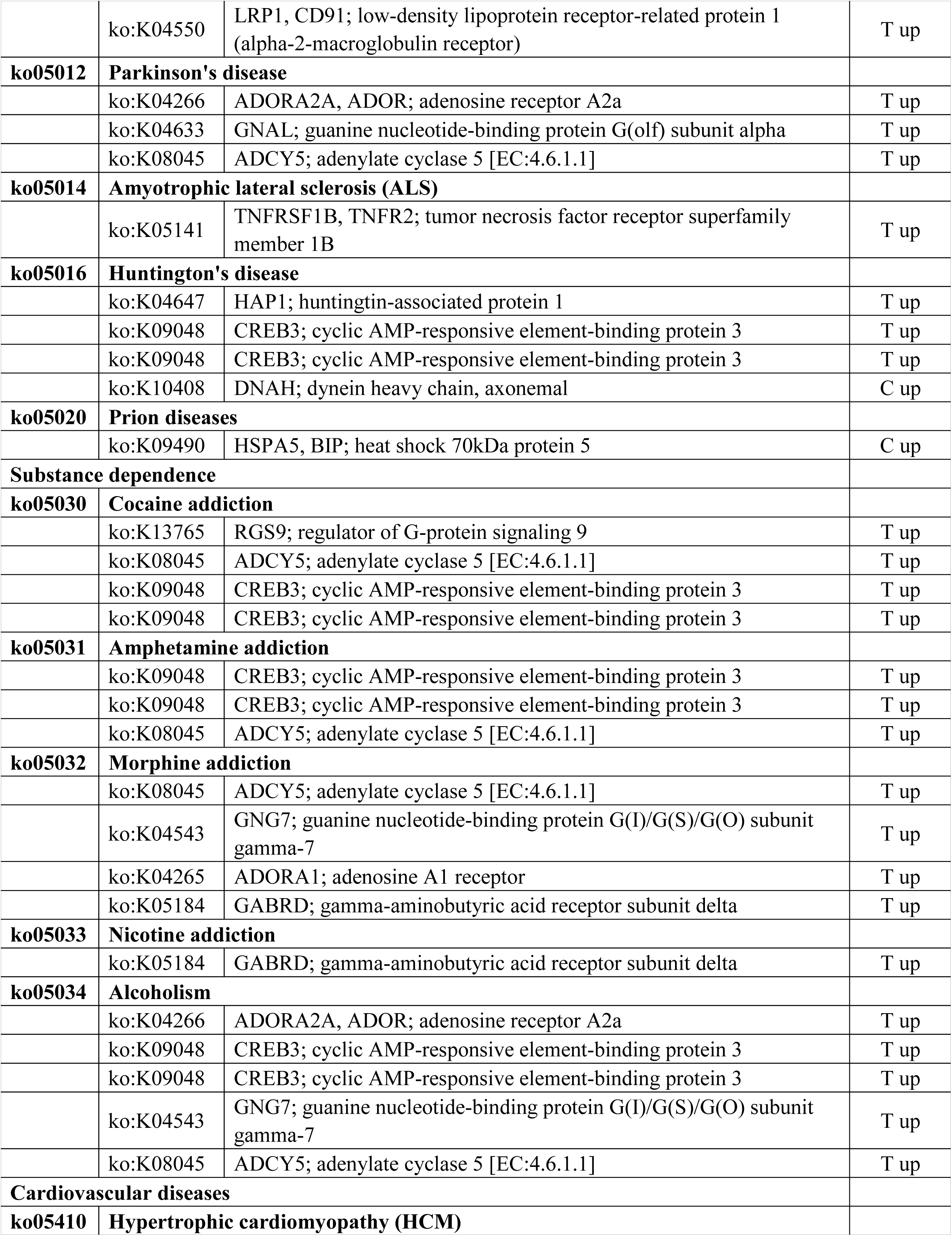
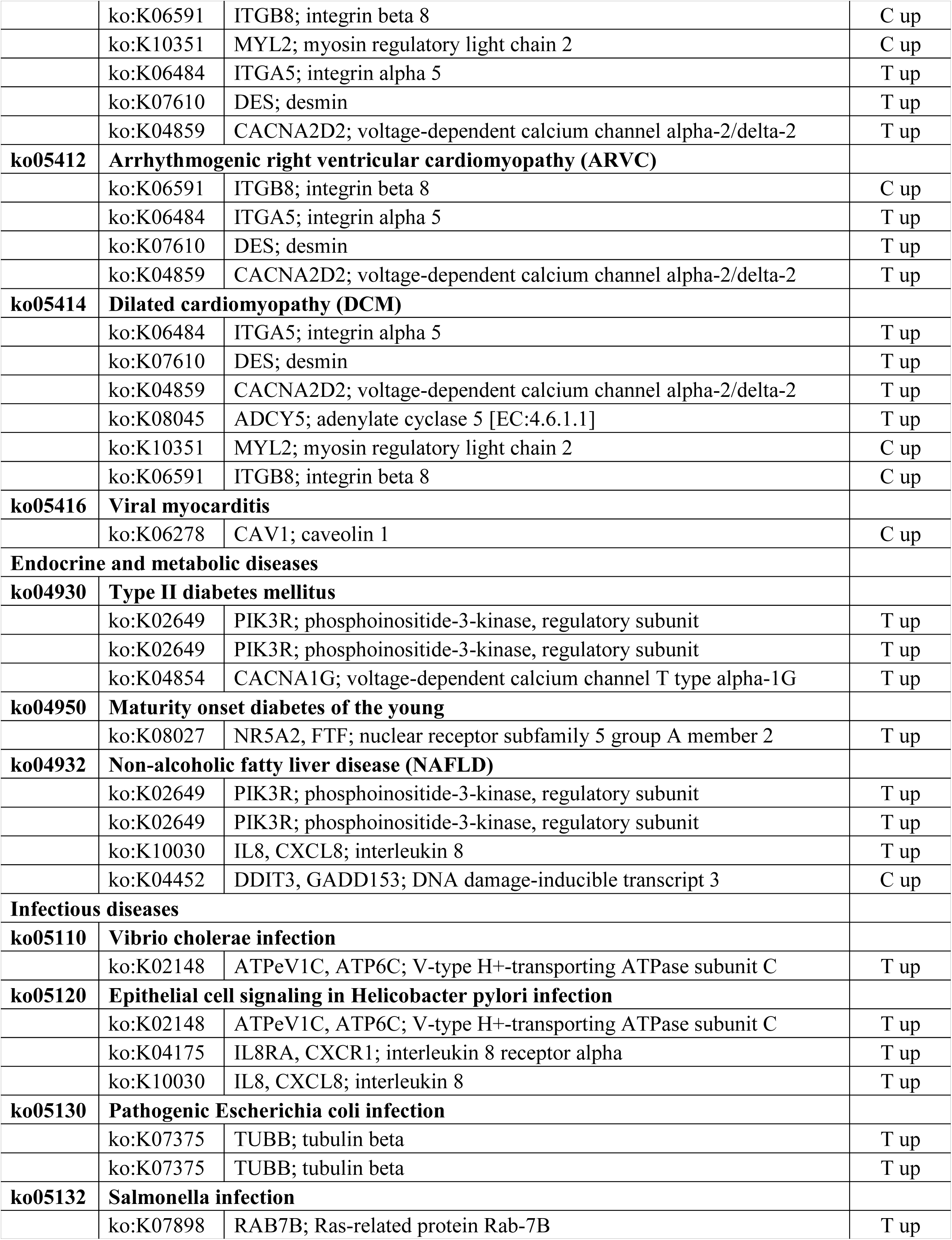
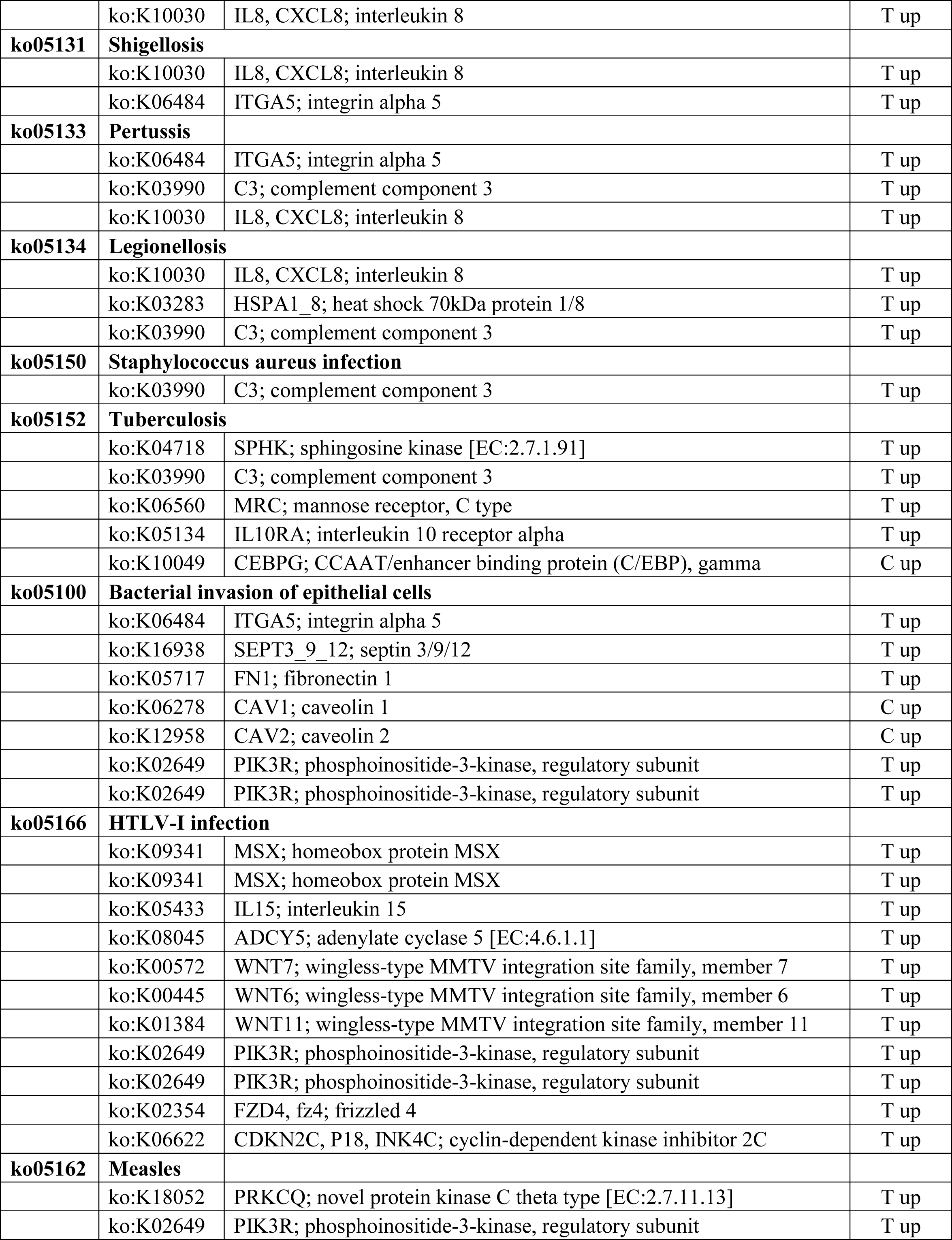
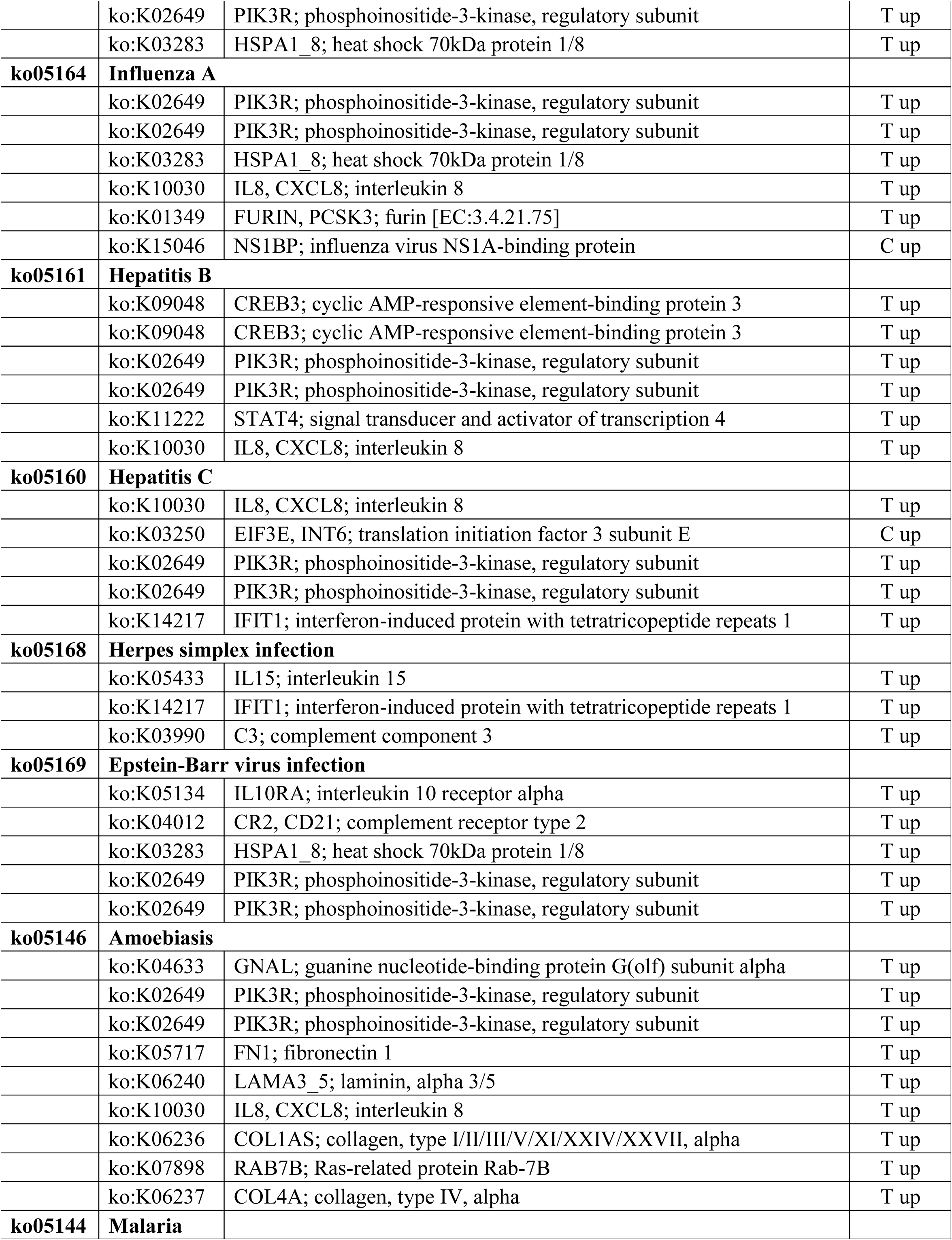
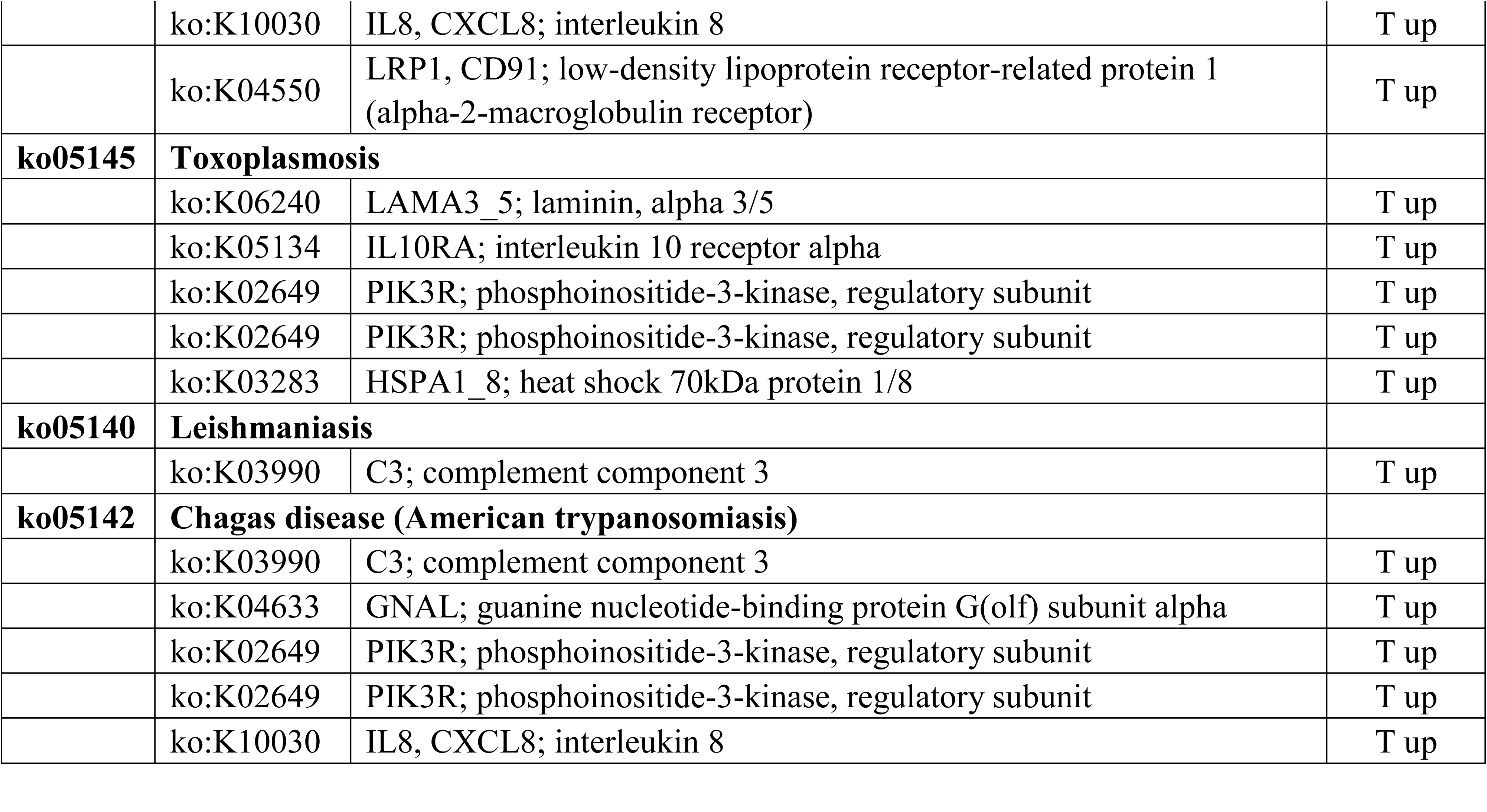
Differentially expressed genes and KEGG pathway analysis

**Fig. S1.**
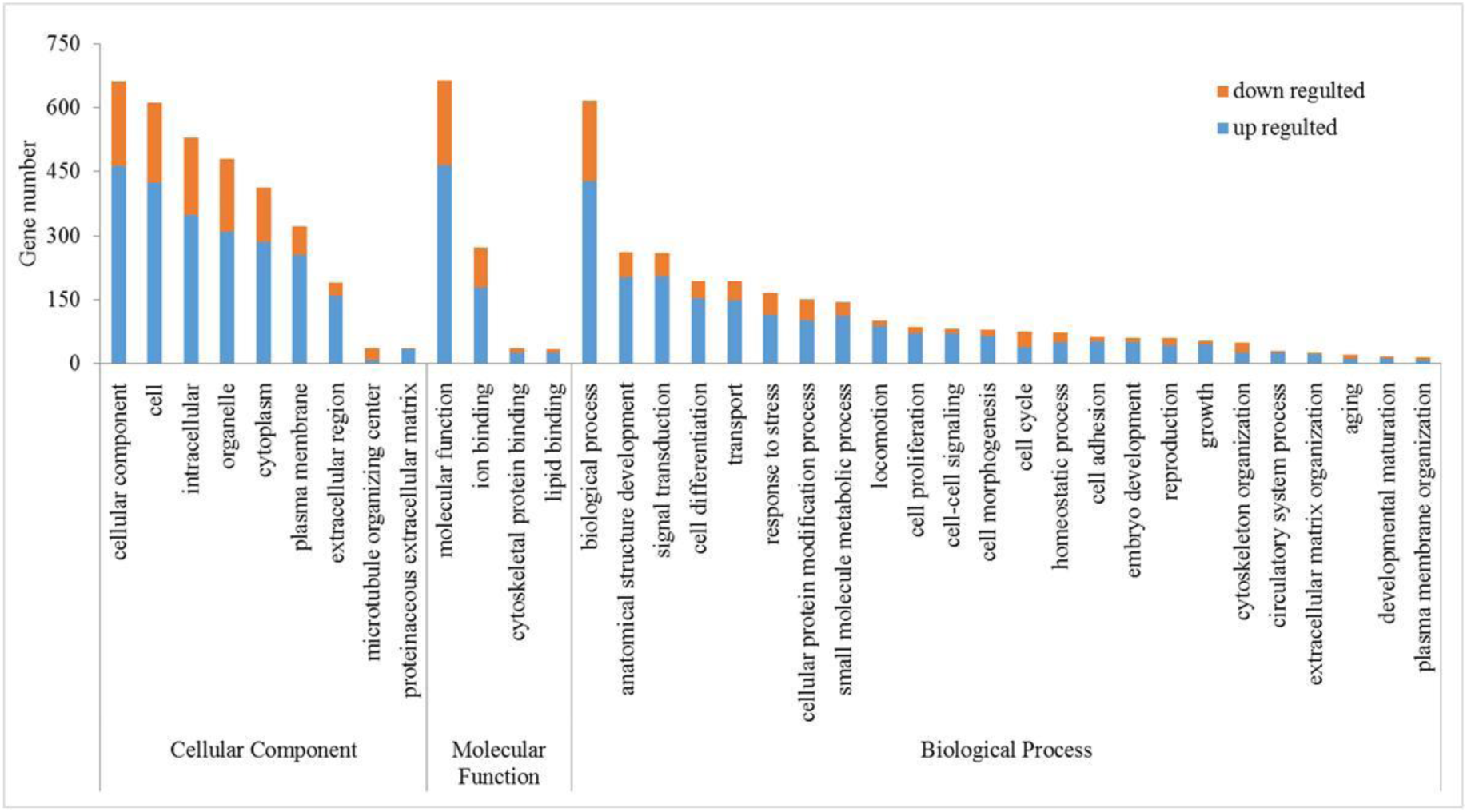
GO annotation of the differentially expressed genes. The genes are enriched in different terms in three functional categories.

**Fig. S2.**
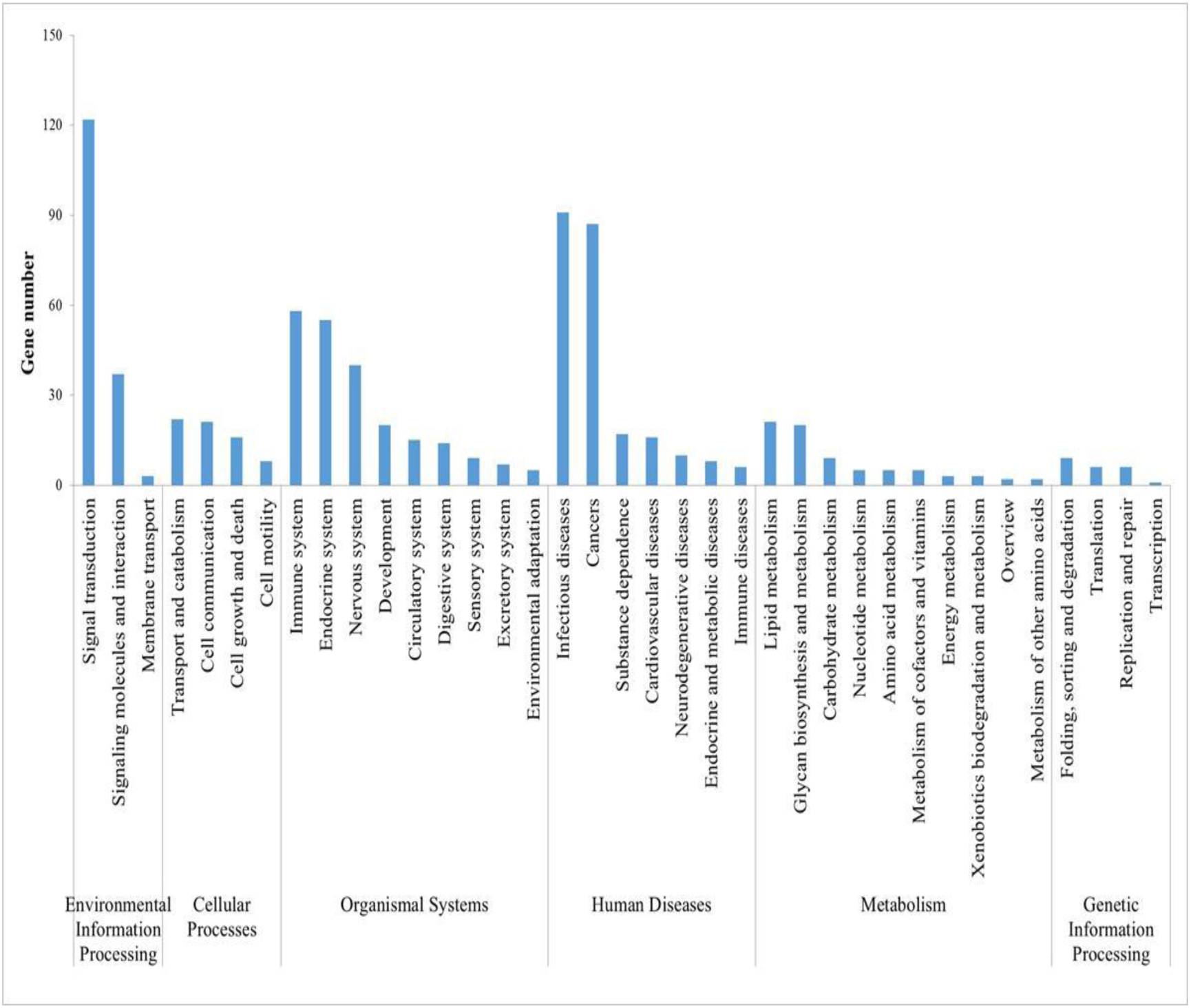
KEGG annotation of the differentially expressed genes. The genes are enriched in different pathways in different functional groups.

**Fig. S3.**
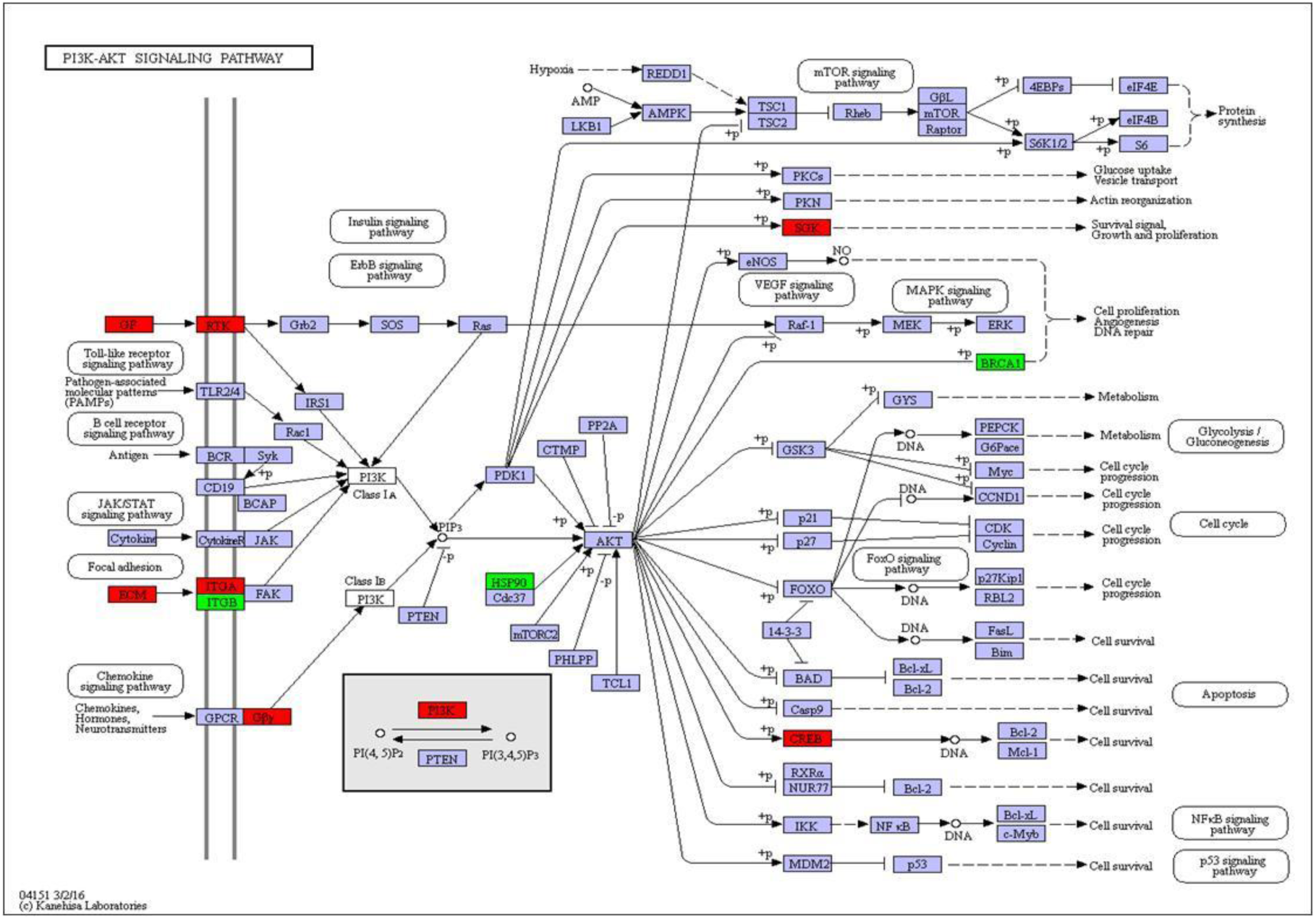
KEGG analysis reveals that AKT pathway is activated by high glucose, insulin and palmitic acid.

**Fig. S4.**
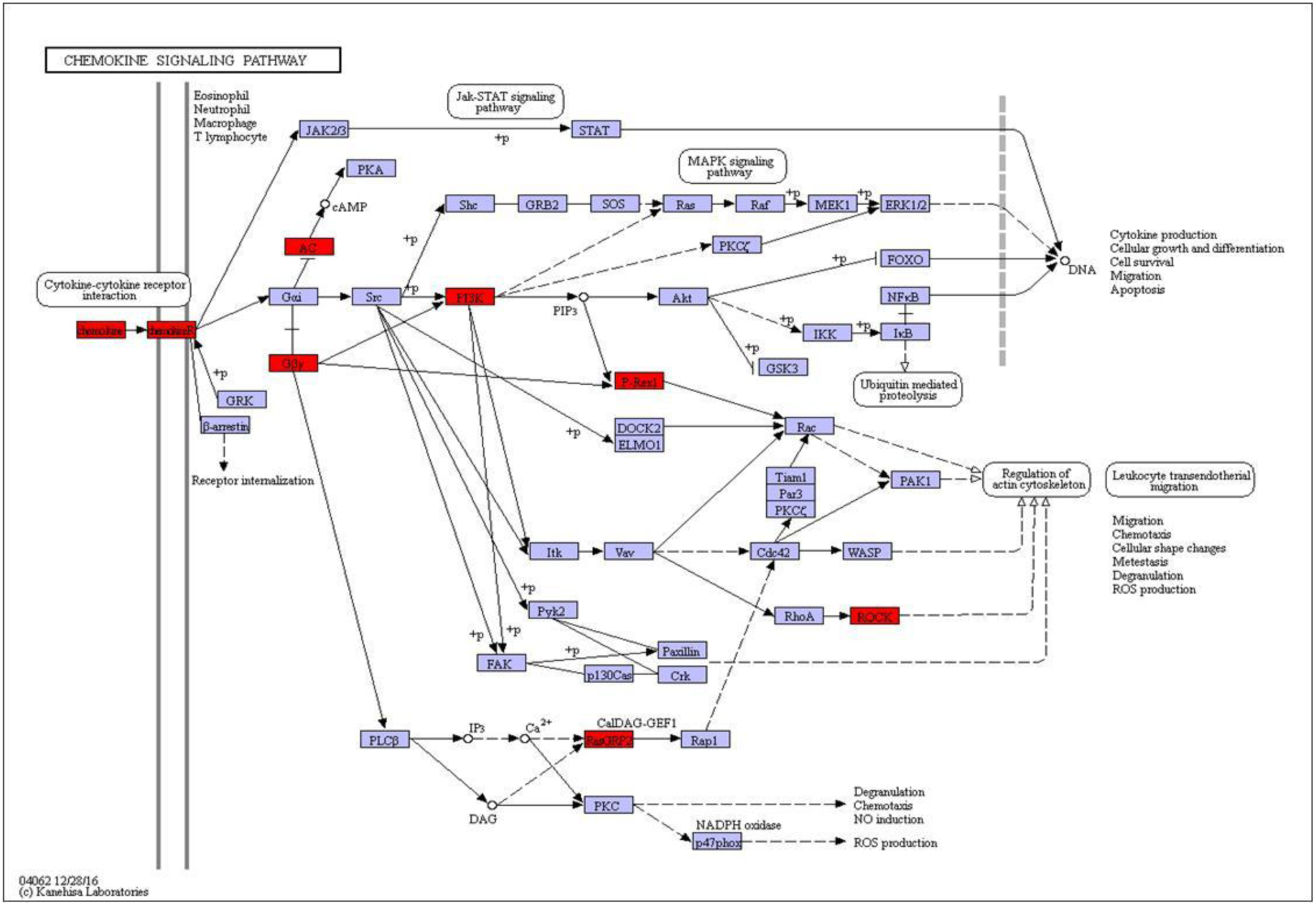
KEGG analysis reveals that ROCK pathway is activated by high glucose, insulin and palmitic acid.

**Fig. S5.**
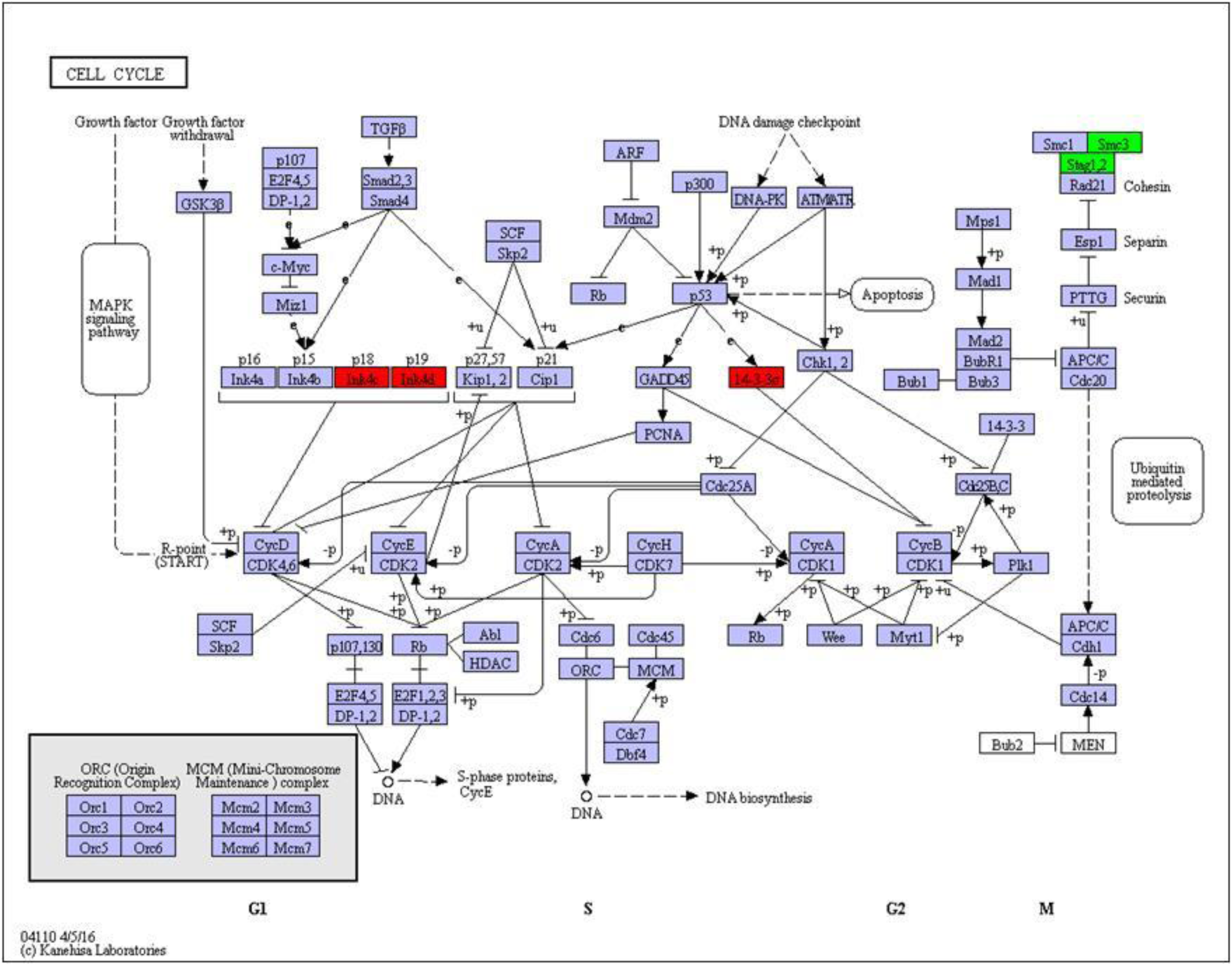
KEGG analysis reveals that 14-3-3σ pathway is activated by high glucose, insulin and palmitic acid.

